# Ayahuasca plant genomes reveal retrotransposon-derived centromeres and contrasting routes to DMT and β-carbolines

**DOI:** 10.64898/2026.10.06.754717

**Authors:** Ruiyi Qiu, Scott Galeung Alexander Mann, Thiago Nascimento, Jacob Owen Perley, Matthew Bailey Richardson, Daniel André Ramey Deschênes, Mirza Bezerra, Arielen Barreto de Carvalho Alves, Anderson Ortiz Alves, Juliana Schietti, Charles R. Clement, John T. Lovell, Victor A. Albert, Gustavo Souza, Cicero Almeida, Yang Qu, André Marques

## Abstract

Ayahuasca, used religiously for centuries by Amazonian Indigenous peoples and increasingly studied for its therapeutic potential, owes its psychoactivity to two plants: *Psychotria viridis* supplies *N,N*-dimethyltryptamine (DMT), whereas *Banisteriopsis caapi* supplies β-carbolines that render DMT orally active. Here we show that the two plants have distinct genome histories: *P. viridis* is hexaploid (2*n* = 6× = 66), consistent with autopolyploidy, and *B. caapi* is diploid (2*n* = 2× = 20). CENH3-bound regions of both species are nonetheless enriched in CRM-type Ty3/Gypsy retrotransposons, with additional LINE elements and CRM-derived satellites in *B. caapi*. In *P. viridis*, a single multifunctional methyltransferase catalyzes both *N*-methylations of tryptamine to DMT. In *B. caapi*, β-carboline biosynthesis combines dedicated enzymes with an acetaldehyde-dependent Pictet-Spengler cyclization that can proceed non-enzymatically, followed by enantioselective oxidation to harmaline. These resources link the ethnobotanical history of ayahuasca to its molecular basis, providing genomes of both source plants, the genetic structure of traditionally recognized cultivated varieties, and the enzymes behind the alkaloids long combined in this decoction.

## Introduction

Hoasca (hereafter ayahuasca) is a psychoactive decoction traditionally prepared from the stems of the liana *Banisteriopsis caapi* (Malpighiaceae) and the leaves of the tree *Psychotria viridis* (Rubiaceae). For centuries, ayahuasca has been central to the ritual, medicinal and cosmological life of Indigenous peoples of the western Amazon, and during the twentieth century it became the sacrament of syncretic Brazilian religions, most prominently the União do Vegetal (UDV) and Santo Daime, whose ceremonial use is now legally protected in several countries^1,2^. This ceremonial use has since spread worldwide. Yet neither plant has a sequenced nuclear genome, and how *B. caapi* produces its β-carboline alkaloids remains unresolved. For the communities that hold ayahuasca sacred, the brew is a “living relative” and a vehicle of healing and spiritual insight rather than a pharmacological curiosity. Genomic study of these plants therefore carries responsibilities as well as opportunities, a point we return to in the Discussion.

The pharmacological activity of ayahuasca arises from a synergy between two classes of tryptophan-derived alkaloids contributed by two distantly related plant species. *P. viridis* supplies *N*,*N*-dimethyltryptamine (DMT), a potent but orally inactive tryptamine, whereas *B. caapi* supplies β-carboline (harmala) alkaloids, chiefly harmine, harmaline and tetrahydroharmine (THH), which act as reversible inhibitors of monoamine oxidase A (MAO-A). By blocking peripheral oxidative deamination of DMT, these β-carbolines allow ingested DMT to reach the central nervous system, so that neither plant alone reproduces the effect of the combination^3,4^. This two-plant pharmacology has made ayahuasca a focus of neuroscience and psychiatry; controlled and observational studies report rapid and sustained antidepressant effects in treatment-resistant depression, with parallel interest in anxiety, substance-use disorders and post-traumatic stress, although the clinical evidence base remains early-stage and demands cautious interpretation^5–7^.

β-Carboline alkaloids are a widespread class of indole-derived metabolites with diverse neuroactive and enzyme-inhibitory activities^8,9^. Their biosynthesis is generally assumed to proceed from tryptamine-derived intermediates, but its enzymatic basis in plants remains poorly resolved^10^. Formation of the β-carboline scaffold requires a Pictet–Spengler condensation between an indoleamine and an aldehyde, which may be catalyzed by enzymes such as strictosidine synthases or proceed spontaneously in aqueous solution^11–14^. Whether plants produce β-carbolines through a strictly enzyme-driven route or through a system that also exploits intrinsic chemical reactivity has therefore remained an open question. By contrast, DMT can be produced by a minimal enzymatic route of tryptophan decarboxylation followed by sequential N-methylation, as recently reconstituted in microbes and plants^15–17^. Structurally related tryptamine alkaloids may therefore arise through different biosynthetic organizations, from compact, enzyme-constrained systems to pathways in which enzymatic and non-enzymatic steps act together.

Despite their functional interdependence, *P. viridis* and *B. caapi* have remained genomically underexplored. Existing resources are limited to organellar genomes, including mitochondrial and chloroplast sequences for *B. caapi*^18,19^ and plastid resources for *P. viridis* and other *Psychotria* taxa^20,21^, with no chromosome-scale nuclear assembly for either species. Consequently, their genome architecture, repeat and centromere biology, polyploid history and the genomic context of alkaloid biosynthesis have remained uncharacterized.

Here we present high-quality haplotype-phased chromosome-scale genome assemblies of *P. viridis* and *B. caapi* and combine them with CENH3 ChIP-seq, transcriptomics, population resequencing and functional biochemistry. We show that *P. viridis* is a hexaploid whose six monoploid chromosome sets correspond one-to-one to the chromosomes of the diploid *Coffea canephora*, and that two traditionally recognized *B. caapi* varieties share a conserved diploid genome architecture. In both species, centromeres are enriched in CRM-type Ty3/*Gypsy* retrotransposons, whereas *B. caapi* centromeres additionally accumulate LINE elements and CRM-derived satellites. Genetic structure among 135 *B. caapi* accessions tracks geographic origin rather than varietal designation. Finally, DMT production in *P. viridis* proceeds through a compact pathway centered on a multifunctional methyltransferase, whereas β-carboline biosynthesis in *B. caapi* combines dedicated enzymes with an acetaldehyde-dependent cyclization that can proceed non-enzymatically. We propose that the two plants combined in ayahuasca rely on contrasting pathway organizations to elaborate tryptamine.

## Results

### *P. viridis* is a hexaploid with six collinear monoploid chromosome sets

Using PacBio HiFi long reads assembled with hifiasm in Hi-C phasing mode, we generated a chromosome-scale assembly of the *P. viridis* genome, resolving all 66 chromosomes (2*n* = 6× = 66; *x* = 11) into six highly contiguous, well-oriented monoploid sets (g1–g6; Figures 1A and S1A–S1D; Tables S1 and S2). BUSCO completeness (embryophyta_odb12) was 98.7% for the complete assembly and 86.4%–94.8% for the six monoploid sets (Figure S1E; Tables S1 and S2). *K*-mer-based analyses with GenomeScope were consistent with a hexaploid model, with the *k*-mer pair-coverage distribution showing the multiplicity structure expected for six homologous copies, an estimated monoploid genome size of ≈1.47 Gb, only 33.9% unique sequence, and a GenomeScope heterozygosity parameter of 8.24%, indicative of substantial divergence among the six homologous copies (Figures S1B and S1C). The assembled monoploid sets (1.39–1.51 Gb each; Figure 1A; Table S2) are slightly larger than the *k*-mer estimate; this difference may reflect improved repeat resolution as well as uncertainty in *k*-mer-based modeling. The six monoploid sets show extensive collinearity along their full length, with fine-scale structural variation unevenly distinguishing individual groups and indicating localized divergence among homologous chromosome copies, for example a large inversion on Chr02 (Figure 1B). Syntenic paralogous gene pairs among the monoploid sets showed a single prominent mode of synonymous substitutions per synonymous site (*K_s_*) at ≈0.025, whereas orthologous gene pairs between *P. viridis* and *Coffea canephora* peaked at *K_s_* ≈ 0.7 (Figure 1C). Thus, the six monoploid sets diverged long after the separation of the *Psychotria* and *Coffea* lineages, consistent with a recent polyploid origin. Further comparison with the diploid Rubiaceae reference species *C. canephora* (*x* = 11) using GENESPACE revealed clear chromosome-scale one-to-one correspondence across all 11 linkage groups, with individual *P. viridis* chromosomes tracing to single *Coffea* chromosomes plus additional intragenomic translocation blocks (e.g. Chr02/Chr09 and Chr06/Chr10; Figures 1D and 1E), consistent with an autopolyploid interpretation on the shared Rubiaceae paleopolyploid background^22,23^ (Figures S2A and S2B). SubPhaser analysis did not resolve distinct subgenome-specific k-mer groups among the six monoploid sets, consistent with an autopolyploid origin (Figure S3). For clarity, g1–g6 are treated here as reconstructed monoploid chromosome sets rather than as inferred parental subgenomes. While the collinearity and SubPhaser results are consistent with an autopolyploid interpretation, they are not diagnostic and do not exclude allopolyploidy involving closely related parental lineages.

**Figure 1.**
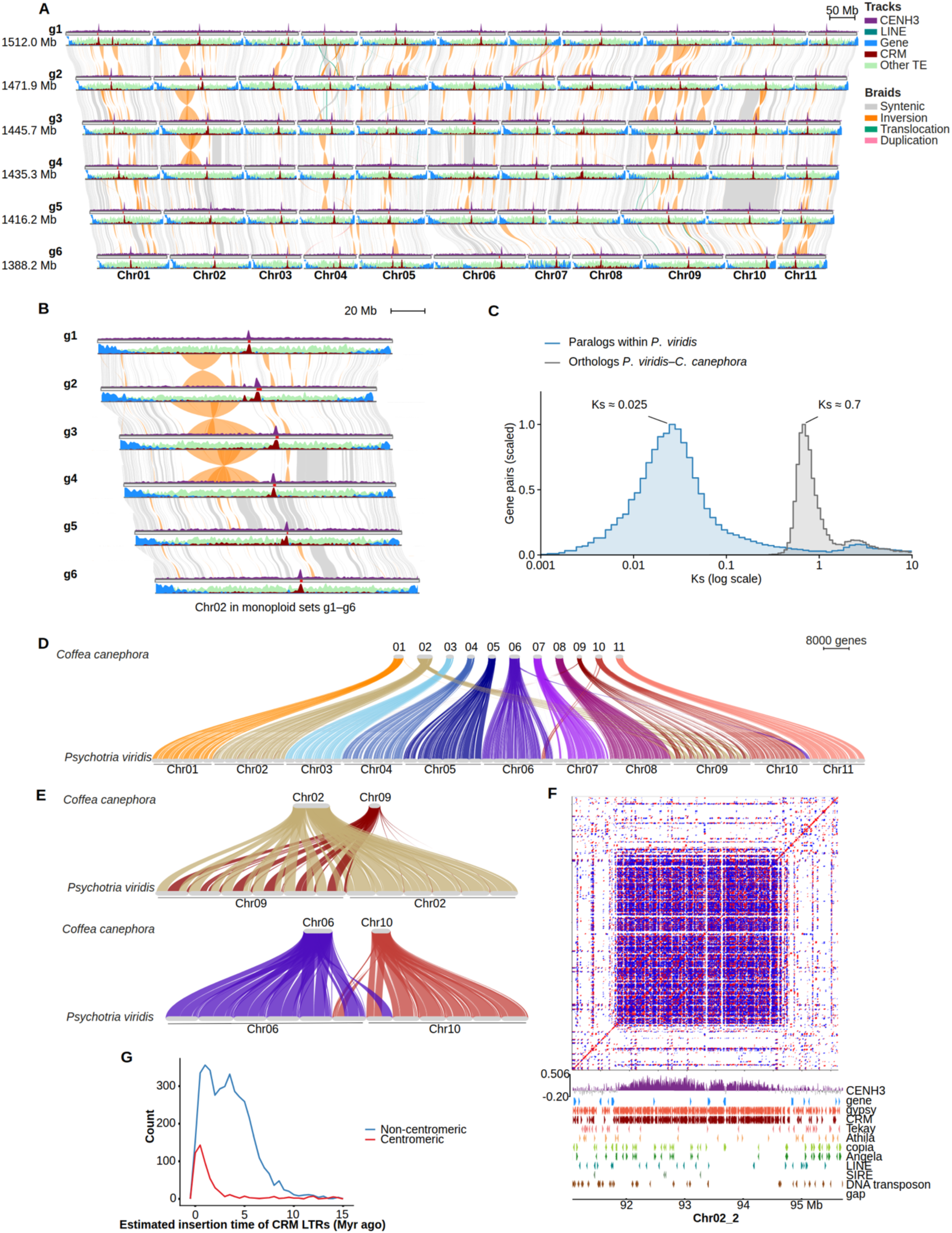
Chromosome-scale architecture of the hexaploid *Psychotria viridis* genome. **(A)** Genome-wide feature and synteny landscape of the six monoploid sets g1–g6. For each monoploid set, tracks show CENH3 ChIP-seq enrichment, density of genes, CRM retrotransposons, LINE elements and other transposable elements; braids between monoploid sets denote syntenic blocks (gray) and structural variants (inversions, orange; translocations, green; duplications, pink); pink strips on the chromosome ideograms represent the identified centromere regions. **(B)** Zoom-in view of Chr02 g1–g6. **(C)** Distributions of *K_s_* (log scale) for syntenic paralogous gene pairs within *P. viridis* and for orthologous gene pairs between *P. viridis* and *Coffea canephora*, computed with CoGe SynMap; each distribution is scaled to its mode. **(D)** GENESPACE riparian plot showing one-to-one chromosome-scale synteny between *P. viridis* and the diploid Rubiaceae reference species *Coffea canephora* (x = 11). **(E)** Detailed depiction of synteny for representative chromosome pairs (Chr02/Chr09 and Chr06/Chr10) illustrating intragenomic translocation blocks. **(F)** Self-similarity dotplot of a representative centromere (Chr02, g2) aligned with CENH3, gene and repeat-lineage tracks (Ty3/*Gypsy*, CRM, Tekay, Athila, Ty1/*Copia*, Angela, LINE, SIRE, DNA transposons), showing CRM-dominated centromeric organization. **(G)** Distribution of estimated CRM LTR insertion times for centromeric versus non-centromeric copies. See also Figures S1–S7 and Tables S1 and S2.

Interspersed repeats make up 70.86% (6.14 Gb) of the 8.67-Gb *P. viridis* assembly, dominated by Ty3/Gypsy (31.20%) and Ty1/Copia (13.86%) elements, with 18.46% of LTRs unclassified and smaller contributions from DNA transposons (2.01% CACTA, 2.09% Mutator, 1.19% hAT, 0.89% Tc1/Mariner, 0.25% PIF/Harbinger, 0.91% Helitron; Figure S1F). The high repeat fraction is consistent with the large genome size, but the timing of repeat accumulation relative to polyploidization remains unresolved^24^. Across the genome, gene density is reduced in regions of CENH3 ChIP-seq enrichment, which marks the functional centromeric regions, and is higher across the chromosome arms (Figures 1A, 1B and S4). Among Ty3/Gypsy lineages, CRM elements tend to co-localize with CENH3 signal peaks, whereas Tekay and Athila are distributed more broadly across chromosome arms. Ty1/Copia lineages (Angela, SIRE) show a comparatively even distribution along chromosome length. DNA transposons and LINEs occur at lower density than LTR retrotransposons across all chromosomes (Figure S4).

Zoomed-in views of individual centromeres confirm that CRM elements and associated LTR retrotransposons dominate the CENH3-enriched regions with little LINE intrusion (Figures 1F and S5), consistent with the paradigm of CRM dominance in angiosperm centromeres^25,26^. Centromeric CRM LTR pairs showed higher similarity and younger insertion ages than genome-wide copies (Figures 1G and S6A; Mann–Whitney *U* test for g2, *W* = 513,868, *P* < 2.2 × 10⁻¹⁶). Phylogenetic analysis of the RT domain further revealed that centromeric CRM copies cluster into distinct clades significantly enriched for centromeric insertions (Figure S7; one-sided Fisher’s exact test, BH-adjusted q < 0.05, enrichment ratio > 0.35). Thus, these results indicate centromere-biased amplification or preferential retention of specific CRM lineages.

### Two *B. caapi* varieties share a conserved diploid genome architecture

We assembled two culturally recognized cultivated varieties of *B. caapi*, Tucunacá (TCN) and Caupuri (CPR), each to chromosome scale with two phased haplotypes (2*n* = 20; Figures 2A and 2B). Hi-C contact maps of CPR show a well-resolved diploid structure of 10 chromosomes with strong on-diagonal contiguity and successful haplotype separation (Figure S8A). TCN contigs were scaffolded onto the CPR assembly (STAR Methods). *K*-mer modeling revealed a bimodal spectrum typical of a diploid genome, with an estimated genome size of ≈1.16–1.20 Gb, low heterozygosity (≈0.51–1.18%) and moderate repeat content (≈57%) (Figures S8B–S8E); assembled haplotypes ranged from 1.10 to 1.15 Gb (Figure 2A; Tables S1 and S2). BUSCO completeness (embryophyta_odb12) was 98.9% and 98.8% for the TCN and CPR assemblies, respectively, and 98.2%–98.3% for the individual haplotypes (Figure S8F; Table S1). SNPs between the phased haplotypes, identified with SyRI, were less frequent within CPR (2.6 per kb of aligned sequence) than within TCN (6.1 per kb), consistent with the k-mer estimates, and SNP density between TCN_h2 and CPR_h1 (6.6 per kb) was similar to that within TCN (Figure 2C). In CPR, windows with few or no SNPs formed tracts of several megabases, which covered almost all of chromosome 5 and most of chromosomes 1 and 6, whereas such tracts were rare in TCN (Figure 2C). These tracts suggest extended runs of homozygosity in the CPR genotype.

**Figure 2.**
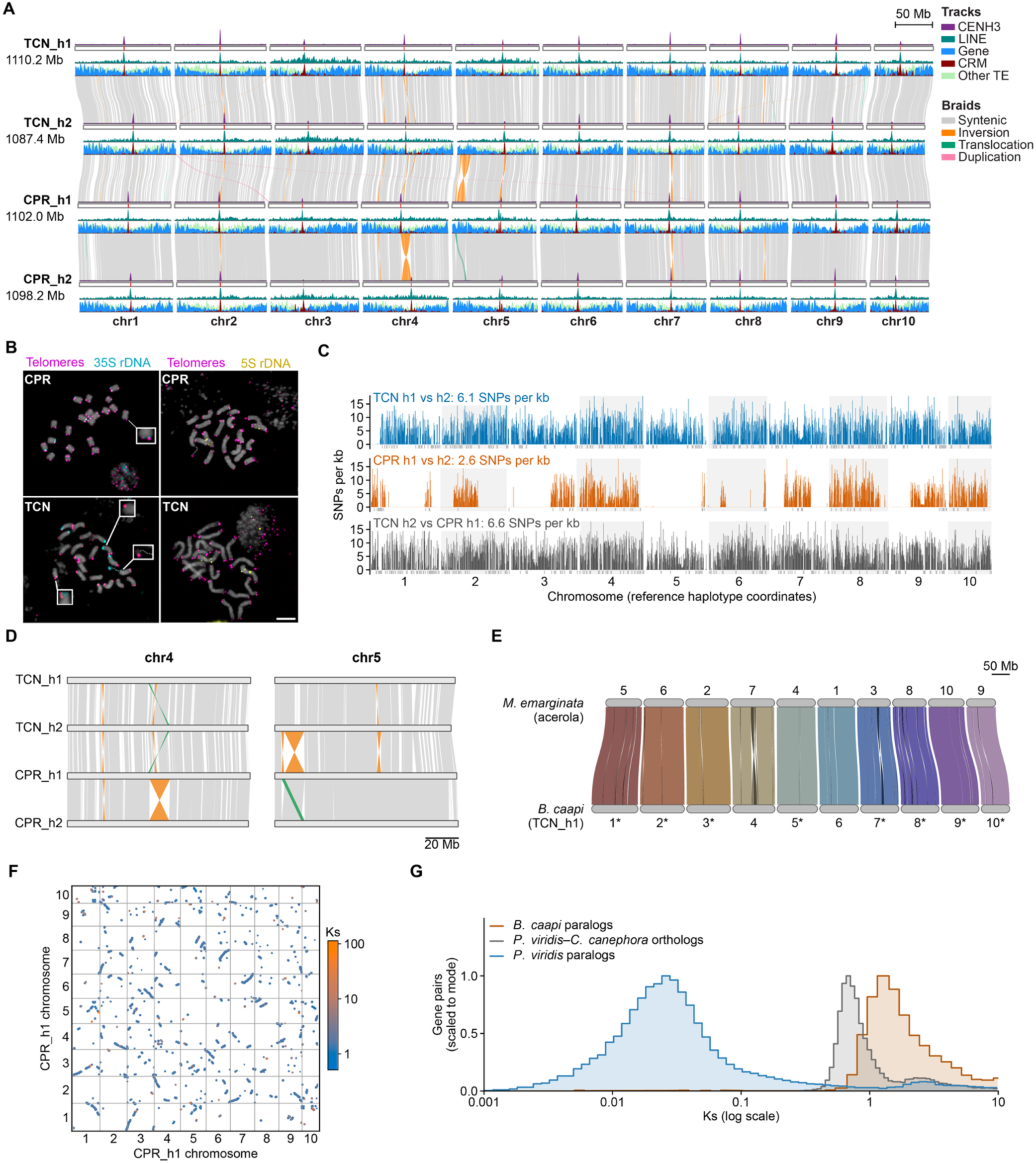
Conserved genome architecture and an ancient whole-genome duplication in two *Banisteriopsis caapi* varieties (Tucunacá, TCN; Caupuri, CPR). **(A)** pyGenomeTracks comparison of the four phased haplotypes (TCN_h1, TCN_h2, CPR_h1, CPR_h2) across the 10 chromosomes, with tracks for CENH3 signals, density of LINE, gene, CRM and other TE, and synteny braids (syntenic, gray; inversion, orange; translocation, green; duplication, pink); pink strips on the chromosome ideograms represent the identified centromere regions. **(B)** Mitotic metaphase chromosomes of CPR (top) and TCN (bottom) after FISH with telomeric (magenta) and 35S rDNA (cyan; left) or 5S rDNA (yellow; right) probes; chromosomes are shown in gray. Insets show 35S rDNA signals close to telomeres (CPR) and small terminal telomeric and 35S rDNA signals (TCN). Scale bar, 5 µm. **(C)** Density of SNPs between phased haplotypes, identified with SyRI, in 500-kb windows along the reference haplotype of each comparison (TCN_h1, CPR_h1 and TCN_h2), expressed per kb of aligned sequence; genome-wide values are given for each comparison. Gray ticks below the axis mark windows with less than 20% aligned sequence; the y axis is truncated at 18 SNPs per kb. **(D)** Synteny and structural rearrangements between the four haplotypes on chromosomes 4 and 5, identified with SyRI (syntenic, gray; inversion, orange; translocation, green; duplication, pink; rearrangements shorter than 200 kb are not shown). **(E)** GENESPACE riparian plot showing chromosome-scale synteny between *B. caapi* (TCN haplotype 1) and acerola (*Malpighia emarginata*, Malpighiaceae). Chromosomes are scaled by physical position and numbered as in each assembly; *B. caapi* chromosomes marked with an asterisk are shown in reverse orientation to match *M. emarginata*. Braids represent syntenic blocks of anchor genes and are colored by *B. caapi* chromosome; darker, crossed braids indicate inversions. **(F)** Self-synteny dot plot of *B. caapi* CPR haplotype 1, redrawn from the CoGe SynMap output (Figure S9A); syntenic gene pairs are colored by *K_s_*. **(G)** Distributions of *K_s_* (log scale) for syntenic paralogous gene pairs within *B. caapi* and *P. viridis*, and for orthologous gene pairs between *P. viridis* and *Coffea canephora*, computed with CoGe SynMap; each distribution is scaled to its mode. *K_s_* values are not directly comparable between lineages because of possible differences in substitution rate. See also Figures S8–S10 and Tables S1 and S2.

Comparison of the TCN and CPR genomes revealed strongly conserved chromosomal architecture, with collinearity maintained across all ten chromosomes and relatively few localized structural variants interrupting otherwise syntenic blocks, including inversions and translocations on chromosomes 4, 5 and 7 (Figure 2A). SyRI identified two large inversions spanning 13.1 Mb on chromosome 4, where CPR_h2 differs from CPR_h1, which is collinear with both TCN haplotypes in this region, and 12.4 Mb on chromosome 5, where TCN_h2 differs from CPR_h1 and neither variety carries the inversion between its own haplotypes (Figure 2D). CPR is therefore heterozygous for the chromosome 4 inversion, whereas the chromosome 5 arrangement distinguishes the two accessions assembled. These events are limited in scale and distribution, suggesting that structural divergence between genotypes is modest rather than genome-wide. Genome-wide annotation of gene and repeat features in the two haplotype-resolved *B. caapi* assemblies (TCN and CPR) shows a broadly similar chromosomal organization between the two genotypes. In both genotypes, gene density is highest toward the chromosome ends and lowest across proximal regions (Figures 2A and S8G–S8H).

Chromosome-scale synteny was also conserved within Malpighiaceae: comparison with the genome assembly of acerola (*Malpighia emarginata*, 2*n* = 20)^27^ showed one-to-one correspondence between the ten chromosomes of the two species, with collinearity maintained along entire chromosomes and interrupted only by a few inversions, the largest on chromosomes 4 and 7 (about 8 and 7 Mb, respectively; Figure 2E). Self-synteny analysis of *B. caapi* revealed numerous large off-diagonal syntenic blocks between non-homologous chromosomes, a pattern indicating the retention of duplicated regions from a whole-genome duplication (WGD) (Figures 2F and S9A). *K_s_* between syntenic gene pairs showed a single prominent mode at log_10_(*K_s_*) ≈ 0.1 (*K_s_* ≈ 1.3), with no distinct peak at lower *K_s_* values, indicating that this WGD is ancient and that no more recent duplication occurred in the *B. caapi* lineage (Figures 2G and S9A). This mode lies above the *K_s_* of orthologous gene pairs between *P. viridis* and *C. canephora* (≈0.7), which diverged within Rubiaceae, and far above that of *P. viridis* paralogs (≈0.025; Figure 2G). Because substitution rates can differ between lineages, this comparison indicates relative rather than absolute age. Consistent with this, fractionation bias analysis showed that genes along each chromosome are retained in syntenic regions distributed across more than one other chromosome, as expected after a WGD followed by fractionation (Figure S9B). Because the chromosomes of *B. caapi* and *M. emarginata* correspond one-to-one (Figure 2E), this WGD predates the divergence of the two genera and is likely shared with *M. emarginata*.

### CRM retrotransposons, LINEs and CRM-derived satellites occupy *B. caapi* centromeres

We next characterized the centromere organization of *B. caapi*. As in *P. viridis*, CRM elements are enriched within the CENH3 domains of individual *B. caapi* centromeres (Figures 2A, 3A, S10 and S11A), where centromeric CRM LTR pairs showed higher sequence similarity and younger insertion ages than genome-wide copies (Figures 3B and S6B; Mann–Whitney U test: TCN haplotype 1, W = 8,644.5, P < 2.2 × 10⁻¹⁶) and clustered into RT-domain clades enriched for centromeric insertions (Figure S12; one-sided Fisher’s exact test, BH-adjusted q < 0.05, enrichment ratio > 0.6). Thus, centromere-biased amplification or preferential retention of specific CRM lineages is shared by both species.

**Figure 3.**
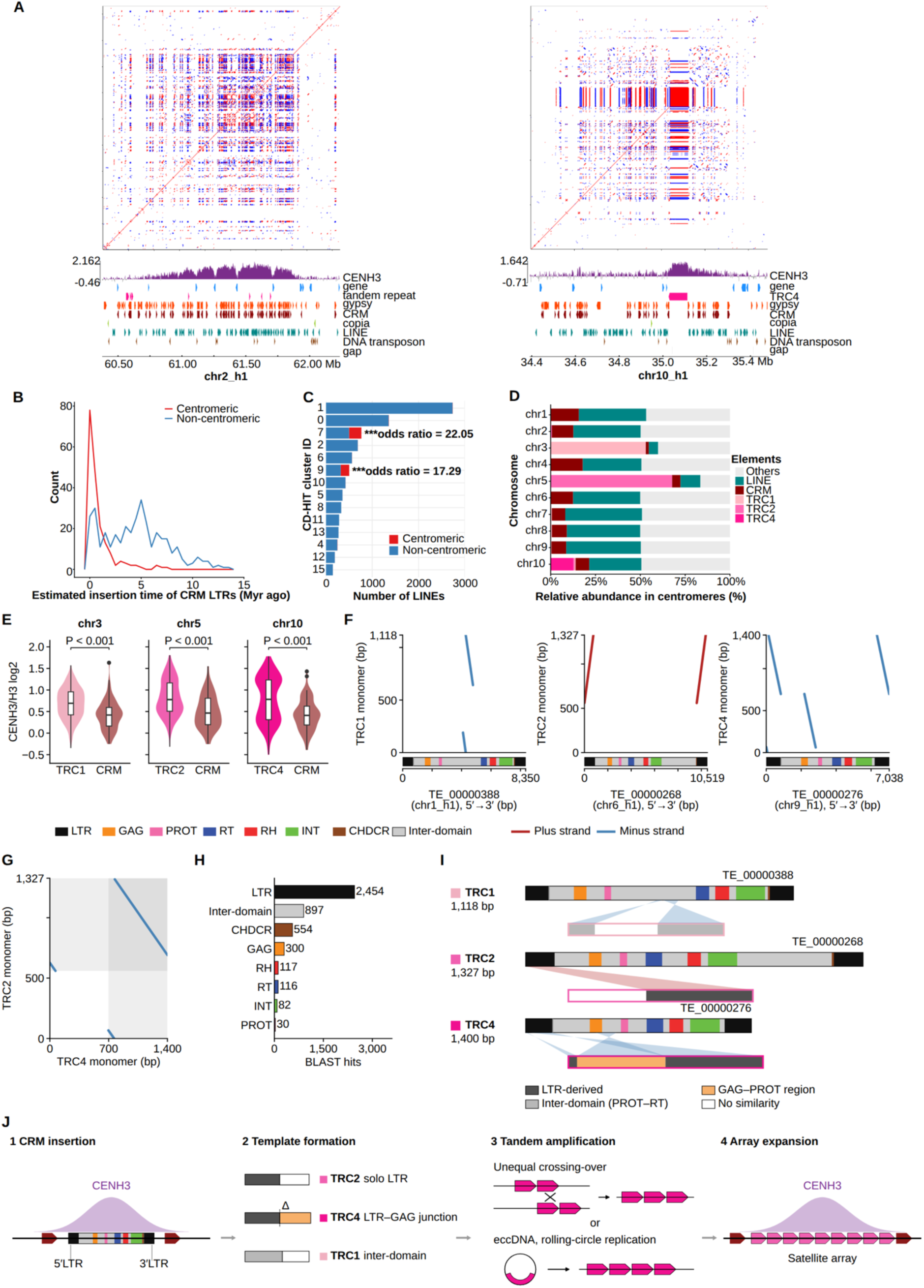
Centromeres of *Banisteriopsis caapi* combine CRM retrotransposons, LINEs and CRM-derived satellites. **(A)** Self-similarity dotplots of representative centromeres (chr2_h1 and chr10_h1) with CENH3, gene, satellite (tandem repeat/TRC4), Ty3/*Gypsy*, CRM, Ty1/*Copia*, LINE and DNA-transposon tracks, showing interspersion of LINE arrays with CRM on chr2 and, on chr10, a TRC4 satellite block in the centromeric region. **(B)** Estimated CRM-LTR insertion times for centromeric versus non-centromeric copies. **(C)** Number of LINE copies per CD-HIT family cluster, split by centromeric (red) versus non-centromeric (blue) location; two clusters are significantly enriched in centromeres (Fisher’s exact test with Benjamini–Hochberg (BH) correction, adjusted *P* < 0.001; odds ratios 22.05 and 17.29, respectively). **(D)** Relative abundance of repeat classes (LINE, CRM, satellite families TRC1/TRC2/TRC4, others) within each centromere of *B. caapi* TCN haplotype 1. **(E)** CENH3/H3 log2 enrichment in 1-kb windows over TRC arrays and CRM elements within the satellite-containing centromeres of chromosomes 3 (TRC1), 5 (TRC2) and 10 (TRC4) of TCN haplotype 1, redrawn from the original violin plots; boxes show the median and interquartile range. Windows were assigned to TRC where TRC co-occurred with CRM or LINE; Wilcoxon rank-sum test (Table S7). **(F)** Dot plots comparing the TRC1, TRC2, and TRC4 monomer consensus sequences (y axis) with representative full-length CRM elements (x axis; domain structure shown below each plot). Elements are shown 5′→3′; TE_00000388 and TE_00000268 are shown as the reverse complement of their genomic orientation. Red, plus strand; blue, minus strand. **(G)** Dot plot comparing the TRC2 and TRC4 consensus sequences. Shaded areas mark the LTR-derived segments of each monomer, as defined in **(I)**. **(H)** Number of BLAST hits of TRC monomer consensus sequences on CRM domains in the *B. caapi* Tucunacá haplotype 1 assembly (n = 4,550 hits; LTR, 53.9%; inter-domain regions, 19.7%). **(I)** Positions of the CRM-derived segments within each TRC monomer consensus, derived from the alignments in **(F)**. Ribbons link aligned segments of the element (top) and of the monomer (bottom); crossed ribbons indicate inverted orientation. For TRC2 and TRC4, only alignments with the 5′ LTR are shown; the 3′ LTR aligns with the same monomer segments **(F)**. Coordinates are approximate. The main centromeric arrays of TRC1, TRC2, and TRC4 lie on chromosomes 3, 5, and 10, respectively, of TCN haplotype 1 (D). **(J)** Proposed model for the origin of CRM-derived centromeric satellites. (1) A CRM element inserts into the CENH3 domain; dark red arrows indicate other CRM copies. (2) A CRM-derived template forms, for example a solo LTR generated by LTR–LTR recombination together with adjacent sequence (TRC2), an internally deleted element in which the LTR is joined to the GAG–PROT region (Δ; TRC4), or an inter-domain segment between PROT and RT (TRC1). (3) The template is amplified in tandem, for example by unequal crossing-over or by rolling-circle replication of extrachromosomal circular DNA (eccDNA) followed by reintegration. (4) Array expansion and homogenization produce chromosome-specific centromeric satellite arrays. Steps 2 and 3 are hypothetical and were not tested here; the alternative routes are not mutually exclusive. See also Figures S6 and S10–S12 and Tables S7 and S8.

However, CRM elements do not dominate the *B. caapi* centromeres exclusively: LINE (long interspersed nuclear element, non-LTR) retrotransposons are also strongly enriched, with self-similarity dot-plot analysis showing CRM and LINE sequences arranged in an intercalated pattern and LINE copies occasionally nested within CRM elements (Figures 3A, S10 and S11A). Clustering of all annotated LINE copies by sequence similarity using CD-HIT identified two clusters significantly over-represented among centromeric LINEs relative to non-centromeric copies (Figure 3C; Fisher’s exact test with Benjamini–Hochberg correction; odds ratios of 22.05 and 17.29, respectively; adjusted *P* < 0.001 for both). In addition, chromosome-specific satellite arrays (tandem-repeat consensus families TRC1, TRC2 and TRC4) contribute substantially to particular centromeres, for example TRC1 on chromosome 3, TRC2 on chromosome 5, and TRC4 on chromosome 10 of *B. caapi* TCN haplotype 1 (Figures 3D, S10 and S11A).

TRC1 and TRC2 make up about half and two-thirds of the centromeric sequence of chromosomes 3 and 5, respectively, exceeding the combined contribution of CRM and LINE elements on these chromosomes (Figure 3D). On chromosome 10, the highest CENH3 signal of the centromere coincides with the TRC4 array rather than with the flanking CRM- and LINE-rich sequence (Figure 3A). Quantification of CENH3/H3 enrichment in 1-kb windows supported this pattern in all three satellite-containing centromeres: median log2 enrichment was higher over TRC arrays than over CRM elements on chromosome 3 (TRC1, 0.69 versus 0.42), chromosome 5 (TRC2, 0.78 versus 0.46) and chromosome 10 (TRC4, 0.78 versus 0.41; Wilcoxon rank-sum test, P < 2 × 10⁻⁶ for each; Figure 3E and Table S7). By contrast, CENH3 enrichment over CRM and LINE elements did not differ consistently between chromosomes: LINE windows were more enriched on chromosome 1, CRM windows on chromosomes 3, 5, 7 and 10, and differences on the remaining five chromosomes were not significant (P > 0.1; Table S8). Centromere metaplots showed CENH3 peaks coinciding with CRM coverage on most chromosomes and with TRC coverage on chromosomes 3, 5 and 10 (Figure S11B).

Dot-plot comparisons of TRC1, TRC2 and TRC4 monomer consensus sequences with CRM sequences revealed syntenic segments, particularly within LTR and non-coding regions, revealing sequence homology between these satellites and CRM elements (Figures 3F and 3G). BLAST analysis of TRC monomer consensus sequences against annotated CRM domains showed that hits were concentrated in the LTR and inter-domain regions, with internal coding domains (RT, RH, INT, GAG) accounting for comparatively few matches (Figure 3H), suggesting that TRC sequences are most closely related to the structural, non-coding portions of CRM elements and may have arisen through ‘satellitization’ of CRM LTR sequences^28,29^. Mapping of these alignments onto the monomer consensus sequences showed that CRM-derived segments cover about 60% of the TRC1 and TRC2 monomers and nearly the entire TRC4 monomer; TRC2 and TRC4 contain LTR-derived segments, whereas the TRC1 segment corresponds to the inter-domain region between PROT and RT (Figure 3I). The similarity between TRC2 and TRC4 is confined to their LTR-derived segments (Figure 3G). We propose that these satellites originated from CRM-derived templates that were amplified in tandem within the CENH3 domain (Figure 3J). Because TRC1 derives from a different CRM region than TRC2 and TRC4, and each satellite forms the main array of a different centromere (Figures 3D and 3I), CRM-derived satellites appear to have originated more than once in *B. caapi* rather than from a single ancestral event. This mixed CRM + LINE + satellite architecture is uncommon in plants, where centromeric non-LTR LINEs have been reported clearly only in sunflower^30^ and poplar^31^, and is consistent with the view that centromere identity is maintained epigenetically by CENH3 rather than by a single conserved sequence class^32^.

### *B. caapi* genetic structure tracks geography rather than varietal designation

To test whether the TCN and CPR varieties represent genetically distinct lineages, we resequenced 135 *B. caapi* accessions from 18 localities in Brazil and analyzed them together with the TCN and CPR reference genomes (137 samples; Table S3). On average, reads mapped to *B. caapi* TCN haplotype 1 reference genome at 98.27 ± 2.08 %, with a mean sequencing depth of 5.97 ± 2.10 × and mean genome coverage breadth of 77.79 ± 12.02 % (Figure S13). After SNP calling and filtering, we reconstructed a phylogenetic network, performed principal component analysis (PCA), and estimated ancestry proportions (Figure 4). In the phylogenetic network, accessions from the northwestern part of the sampled range (RR, PF, CS, TF, RA, SL) formed long, early-diverging branches distinct from a second, less differentiated group comprising the remaining localities (NE, AC, CX, EX, PC, OP, VP, AF, Cba, TR) together with TCN and CPR. Although the TCN and CPR accessions are morphologically distinct, they were sampled from the same plantation and did not form separate, variety-specific clades. Instead, both were nested within the same clade as Cba and other inland accessions (Figure 4A). PCA was consistent with this pattern: most of the genetic variance (PC1, 47.3%) separated accessions along a geographic axis, and k-means-derived clusters (Cluster0–4) corresponded to discrete sampling localities rather than to variety identity (Figures 4B and S14A–S14C). Ancestry analysis across K = 2–8 supported hierarchical, geography-associated ancestry structure: at K = 2, accessions from the northwestern localities separated from all remaining accessions, and additional ancestry components resolved further substructure among the remaining localities with increasing K (Figure 4C). At K = 8, ancestry components were largely correlated with sampling locality, and TCN and CPR still shared a nearly identical ancestry profile, indicating that fine-scale genetic structure closely tracks geographic origin (Figures 4C and 4D). Thus, genetic differentiation among the sampled *B. caapi* accessions is structured primarily by geographic origin rather than by the TCN/CPR varietal designation. However, the TCN and CPR references represent single accessions from the same plantation, and although the ancestry results are consistent with vegetative propagation, they do not by themselves establish clonality among named accessions.

**Figure 4.**
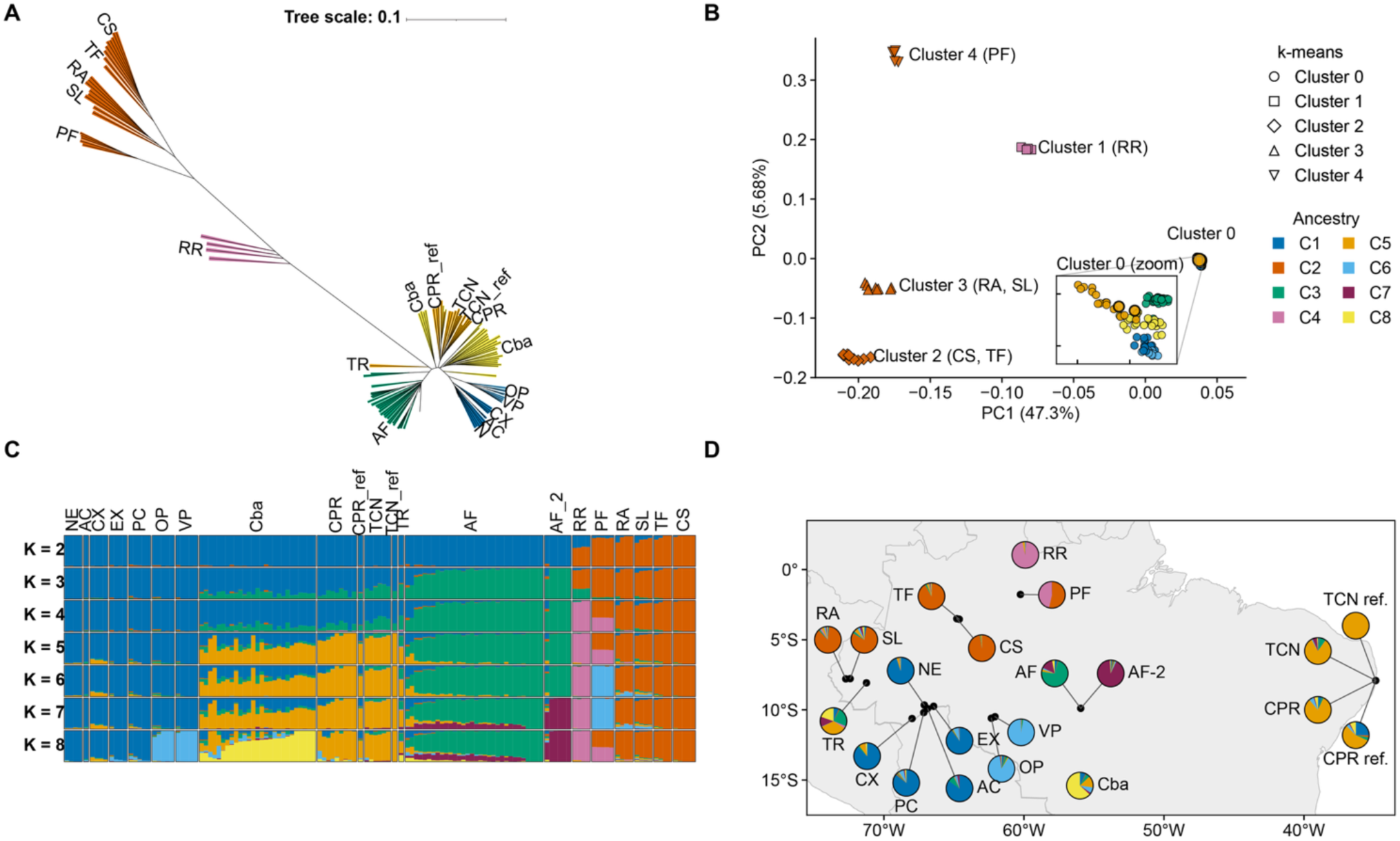
Population structure of *Banisteriopsis caapi*. **(A)** Phylogenetic network of 135 resequenced accessions and the TCN and CPR reference genomes (137 samples); tips colored by the predominant ancestry component (K = 8) of their sampling locality. **(B)** Principal component analysis (PCA) of genetic variation, PC1 versus PC2; colors as in **(A)**, symbols denote k-means-derived genetic clusters (Clusters 0 to 4), and the reference genomes are outlined. The inset enlarges Cluster 0. **(C)** Ancestry proportions for K = 2–8. **(D)** Sampling localities with pie charts showing mean ancestry proportions at K = 8; lines link each pie to the sampling coordinates. The resequenced TCN and CPR accessions and the two reference genotypes were collected from one plantation in Abreu e Lima, Pernambuco, Brazil. See also Figures S13 and S14 and Table S3.

### A single multifunctional methyltransferase completes DMT biosynthesis in *P. viridis*

To determine the DMT pathway in *P. viridis*, we generated high-depth RNA-seq and Iso-Seq datasets from leaves, stems and roots. DMT biosynthesis is expected to proceed through tryptophan decarboxylation to tryptamine by a tryptophan decarboxylase (TDC), followed by one or two N-methylations catalyzed by N-methyltransferases (NMTs) (Figure 5A). Given the high DMT content of *P. viridis* leaves (6.97 mg g^−1^ dry weight; Figure S15), we expected the biosynthetic genes to be strongly expressed in this tissue.

**Figure 5.**
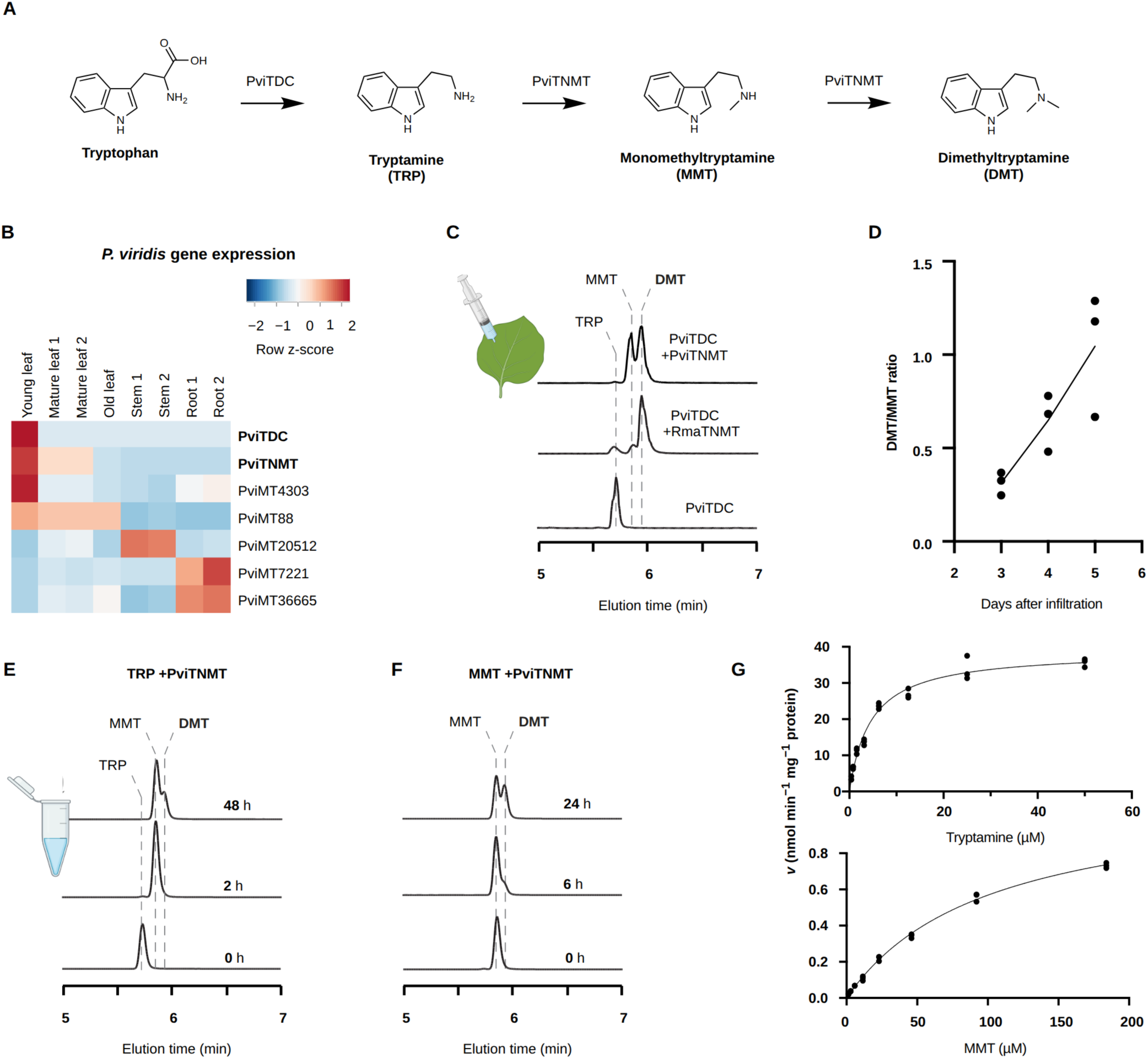
Biosynthesis of *N,N*-dimethyltryptamine (DMT) in *Psychotria viridis*. **(A)** DMT is produced from tryptophan by a tryptophan decarboxylase (PviTDC) and sequential *N*-methylation of tryptamine (TRP) to monomethyltryptamine (MMT) and DMT by a single tryptamine *N*-methyltransferase (PviTNMT). **(B)** Tissue expression heatmap (row *Z*-score) of PviTDC, PviTNMT, and the five next most highly expressed MT candidates across young leaf, mature leaf, old leaf, stem and root; values in Table S4. **(C)** LC–MS/MS extracted-ion chromatograms after *Agrobacterium*-mediated transient co-expression of PviTDC with PviTNMT or the cane-toad RmaTNMT in *Nicotiana benthamiana*, showing de novo MMT and DMT. **(D)** DMT/MMT ratio in *N. benthamiana* leaves 3–5 days post-infiltration (points, biological replicates; line, linear fit). **(E)** In vitro assay of purified His-PviTNMT with tryptamine, showing rapid MMT and delayed DMT formation (0, 2, 48 h). **(F)** In vitro assay with MMT as sole substrate, showing DMT formation over 0–24 h. **(G)** Michaelis–Menten saturation kinetics for tryptamine and MMT; parameters in Table S5. See also Figures S15–S20 and Tables S4 and S5.

RNA-seq identified a young-leaf–enriched TDC-like transcript sharing 75% and 67% amino-acid identity with characterized TDCs from *Camptotheca acuminata* and *Catharanthus roseus*, respectively, and a highly expressed methyltransferase-like transcript with 63% identity to an *O*-methyltransferase from *Carapichea ipecacuanha* and low similarity to other MTs (Figures 5B and S16; Table S4). This methyltransferase transcript was expressed at least 11.5-fold higher than any other MT candidate in young leaf (Figure 5B; Table S4). Heterologous expression of the TDC-like gene in *Saccharomyces cerevisiae* produced tryptamine, and co-expression with the MT-like gene yielded monomethyltryptamine (MMT), confirmed by LC–MS/MS (Figures S17 and S18). We therefore named these enzymes PviTDC and PviTNMT (tryptamine N-methyltransferase). Co-expression of PviTDC with the characterized RmaTNMT from the cane toad *Rhinella marina*^17^ yielded both MMT and DMT, albeit at lower levels (Figures S17 and S18), and transient expression of PviTDC and PviTNMT in *Nicotiana benthamiana* produced both MMT and DMT, with the DMT/MMT ratio rising from 0.31 at day 3 to 1.04 at day 5 (Figures 5C and 5D).

To test whether PviTNMT catalyzes both methylation steps, we purified recombinant His-tagged PviTNMT from *Escherichia coli* (Figure S19). In vitro, MMT formed rapidly (within 2 h) with little DMT, but substantial DMT accumulated by 48 h; using MMT as sole substrate also yielded DMT over 6–24 h (Figures 5E and 5F). No other tested MT candidate was active toward tryptamine or MMT. Kinetic analysis showed markedly higher catalytic efficiency toward tryptamine (*K*_M_ = 4.56 μM; *k*_cat_ = 28.8 × 10−3 s−1) than toward MMT (*K*_M_ = 100.4 μM; *k*_cat_ = 0.84 × 10−3 s−1), a 751-fold difference in *k*_cat_/*K*_M_ (Figure 5G; Table S5). Relative to RmaTNMT, PviTNMT was 1,169-fold more efficient for the first methylation but only 14.8% as efficient for the second (Table S5). Homology modeling provided a structural rationale: tryptamine fits efficiently in the PviTNMT active site, whereas steric constraints limit productive binding of MMT (Figures S20A and S20B). Thus, PviTNMT is a single, multifunctional enzyme that is highly efficient at the first methylation and slower at the second, suggesting that the in vivo balance between MMT and DMT may depend on substrate availability and flux.

### Dedicated enzymes and an acetaldehyde-dependent cyclization produce harmala alkaloids in *B. caapi*

We next investigated harmala-alkaloid biosynthesis in *B. caapi* using high-depth RNA-seq and Iso-Seq from multiple tissues of the TCN and CPR varieties. Metabolite profiling showed that leaves are the primary site of accumulation, with harmine and THH being the two dominant alkaloids and harmaline and 6-methoxytryptamine present at lower levels (Figure S21). We hypothesized a pathway in which tryptamine is methoxylated and then undergoes cyclization to THH, which is oxidized to harmaline and harmine (Figure 6A).

**Figure 6.**
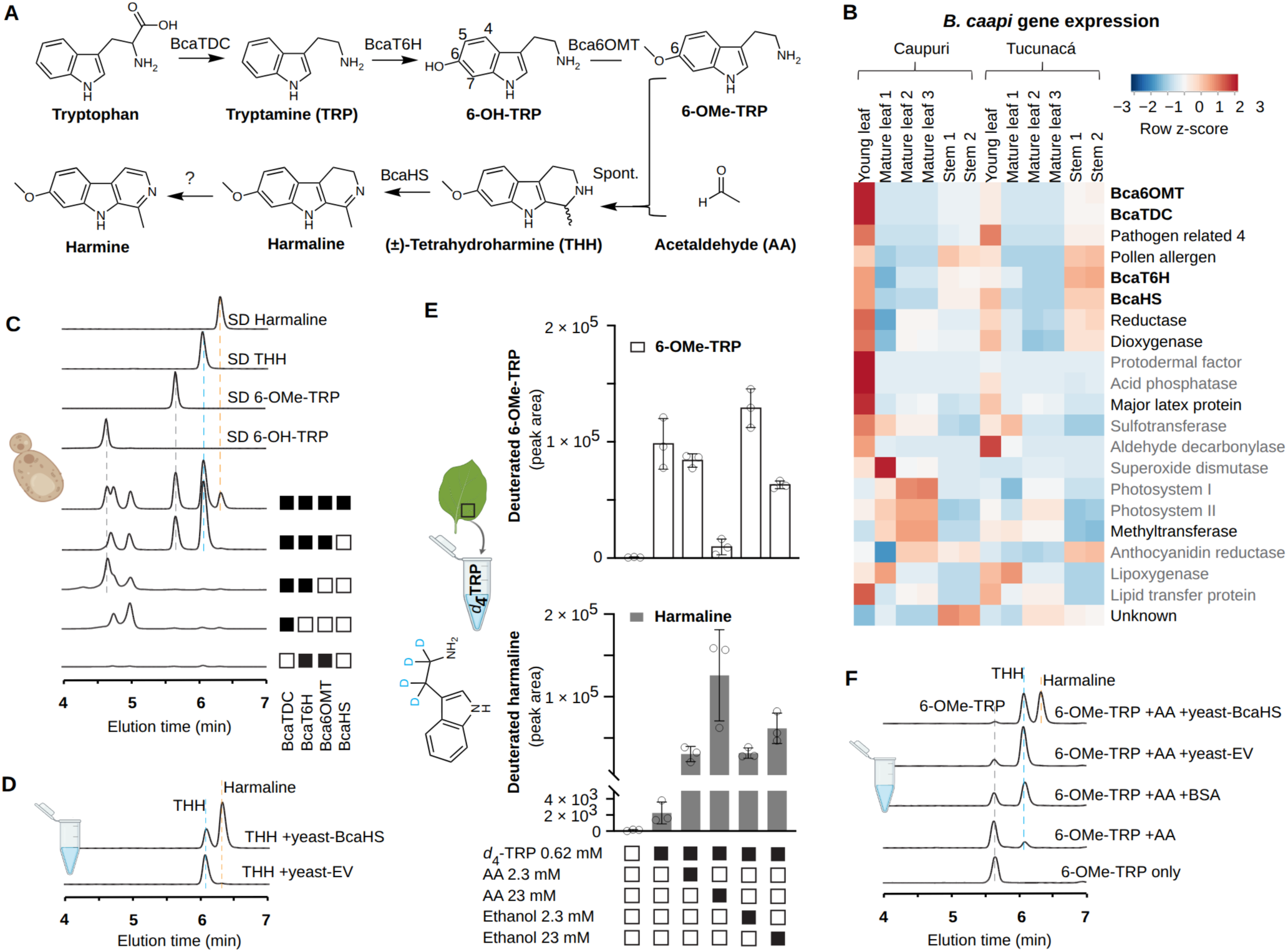
Biosynthesis of β-carboline (harmala) alkaloids in *Banisteriopsis caapi*. **(A)** Proposed pathway: tryptophan → tryptamine (BcaTDC) → 6-hydroxytryptamine (BcaT6H) → 6-methoxytryptamine (Bca6OMT), followed by acetaldehyde-dependent Pictet-Spengler cyclization that can proceed non-enzymatically to (±)-tetrahydroharmine (THH), enantioselective oxidation of *S*-THH to harmaline by harmaline synthase (BcaHS), with the terminal oxidation to harmine unresolved. **(B)** Tissue expression heatmap (row *Z*-score) of the 21 candidate genes in the Caupuri and Tucunacá varieties (young/mature leaf, stem); pathway genes highlighted. **(C)** LC–MS/MS showing stepwise reconstitution in *Saccharomyces cerevisiae*: sequential addition of BcaTDC, BcaT6H, Bca6OMT and BcaHS yields 6-OH-TRP, 6-OMe-TRP, THH and harmaline (filled/open squares indicate enzymes present). **(D)** In vitro oxidation of THH to harmaline by yeast lysate expressing BcaHS versus empty-vector control. **(E)** Isotope feeding of *d*₄-tryptamine to *B. caapi* leaf discs: deuterated 6-OMe-TRP and harmaline with/without acetaldehyde (AA) or ethanol supplementation (bars, mean; points, three biological replicates; error bars, standard deviation). **(F)** In vitro acetaldehyde-dependent formation of THH from 6-OMe-TRP and its BcaHS-dependent conversion to harmaline (with BSA/yeast-lysate acceleration). See also Figures S16 and S21–S28 and Table S6.

To find candidate genes we excluded transcripts sharing >70% amino-acid identity with *Arabidopsis thaliana* proteins and took the 21 most highly expressed young-leaf transcripts, curating these to 11 candidates including a TDC-like enzyme, two methyltransferases, a cytochrome P450 and a berberine-bridge-enzyme (BBE)-like oxidase (Figure 6B; Table S6). In *S. cerevisiae*, the TDC-like enzyme produced tryptamine; addition of the P450 yielded 6-hydroxytryptamine, and a methyltransferase produced 6-methoxytryptamine (Figure 6C), activities also confirmed in *N. benthamiana* (Figure S22). We named these BcaTDC, BcaT6H (tryptamine 6-hydroxylase) and Bca6OMT (6-*O*-methyltransferase). Docking supported regioselective C6 hydroxylation by BcaT6H and 6-O-methylation by Bca6OMT (Figures S24E and S24F). BcaT6H is related to characterized tryptamine 5-hydroxylases from tomato and rice (64.6% and 53.5% identity), and Bca6OMT is only distantly related to characterized alkaloid methyltransferases (closest homolog a *Capsicum annuum* catechol OMT, 52.3% identity) (Figures S16 and S23).

Yeast expressing only these upstream enzymes accumulated THH without any additional *B. caapi* gene, and co-expression of the BBE-like oxidase produced harmaline (Figure 6C); we named this enzyme harmaline synthase (BcaHS). In vitro, BcaHS oxidized THH to harmaline and converted only ≈50% of racemic THH, consistent with strong enantioselectivity (Figures 6D and 7A–7C); modeling and molecular-dynamics simulations supported preferential, stable binding of the *S*-enantiomer and rapid destabilization of *R*-THH (Figures S24A–S24D). Because *S. cerevisiae* generates abundant acetaldehyde during fermentation (intracellular concentration >11 mM)^33^ and tryptamines undergo spontaneous Pictet-Spengler condensation with aldehydes, we reasoned that the THH formed in yeast likely arose non-enzymatically. In planta, feeding deuterated (*d*₄) tryptamine to *B. caapi* leaf discs produced labeled 6-methoxytryptamine and harmaline, and supplementation with acetaldehyde or ethanol (converted to acetaldehyde in vivo) increased harmaline production 12- to 50-fold in a dose-dependent manner, with a corresponding decrease in 6-methoxytryptamine (Figure 6E). Co-infiltration of 6-methoxytryptamine with acetaldehyde or ethanol in *N. benthamiana* yielded THH, whereas 6-methoxytryptamine alone did not (Figure S25).

**Figure 7.**
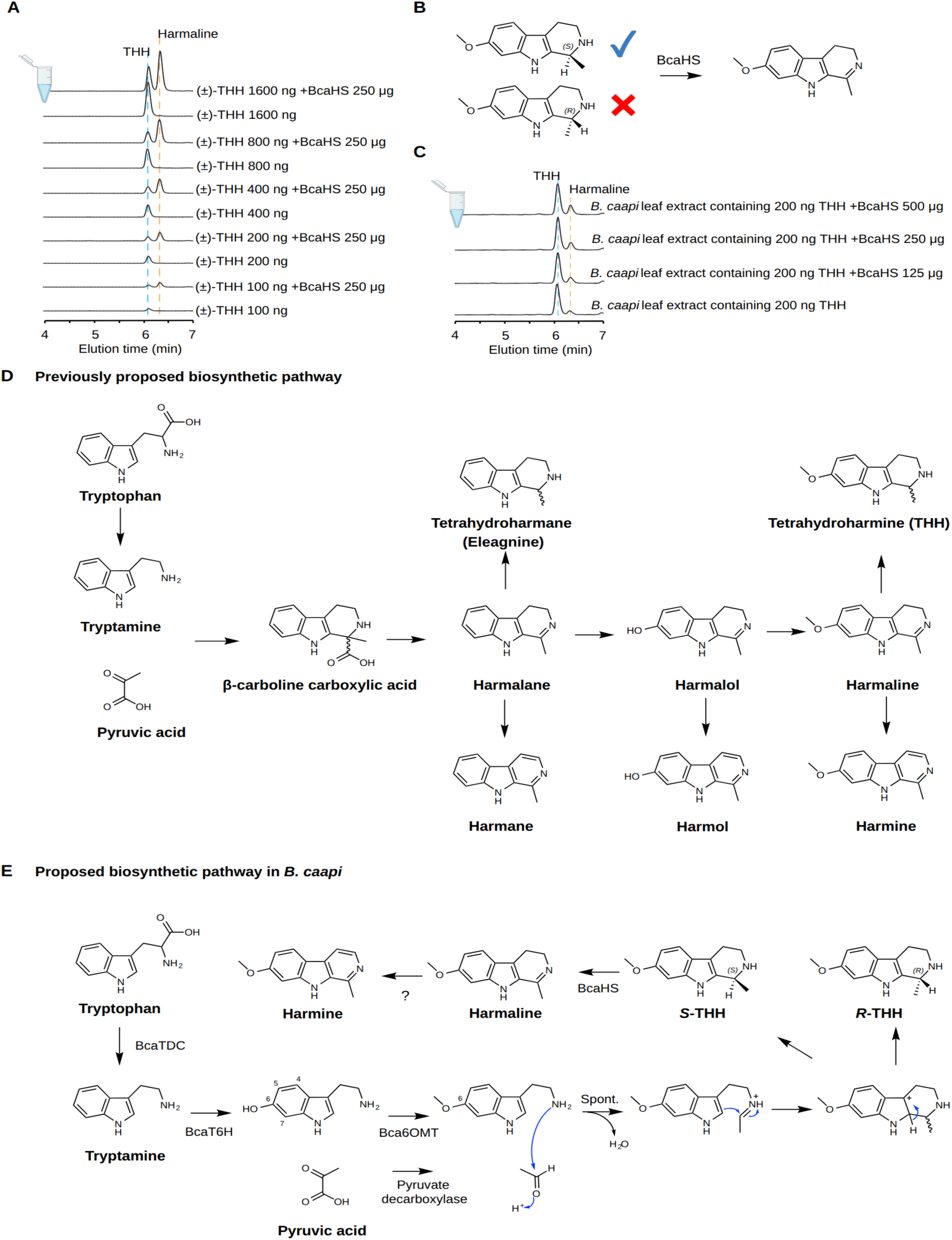
*Banisteriopsis caapi* harmaline synthase (BcaHS) selectively oxidizes the *S*-enantiomer of tetrahydroharmine (THH). **(A)** Approximately 50% of racemic THH was oxidized to harmaline, independent of the amount of THH supplied (100–1600 ng/100 µL reaction), using constant amounts of yeast total lysate (250 µg protein) expressing BcaHS. **(B)** Schematic representation of BcaHS stereoselectivity inferred from homology modeling of BcaHS and docking simulations with either *S*-or *R*-THH (see Figure S24). **(C)** THH present in *B. caapi* leaf extracts (200 ng/100 µL reaction) could not be oxidized to harmaline by yeast total lysate (from 125 to 500 µg protein) expressing BcaHS under identical assay conditions, suggesting that the THH accumulating in *B. caapi* leaves corresponds predominantly to the *R*-enantiomer. All reactions were incubated at 30 °C overnight to ensure completion. LC-MS/MS analysis shows extracted-ion chromatograms (EICs) corresponding to the following [M+H]⁺ ions: *m/z* 217 for THH and *m/z* 215 for harmaline. (**D** and **E**) Proposed biosynthetic pathways for harmala alkaloid biosynthesis. **(D)** Based on feeding experiments with isotope-labeled tryptamine and pyruvic acid, it was previously proposed that harmala alkaloid biosynthesis proceeds through formation of a β-carboline carboxylic acid intermediate generated by a cycloaddition reaction between tryptamine and pyruvic acid, followed by decarboxylation to harmalane and subsequent modifications into various alkaloids^34–38^. **(E)** In contrast, our study identified a tryptophan decarboxylase (TDC), tryptamine 6-hydroxylase (T6H), and 6-hydroxytryptamine *O*-methyltransferase (6OMT) that collectively convert tryptophan to 6-methoxytryptamine, which accumulates in all examined tissues, including young leaf, old leaf, stem, and root (Figure S21). Both in vitro and in vivo evidence supports a spontaneous Pictet-Spengler reaction between 6-methoxytryptamine and acetaldehyde, suggesting that this reaction underlies harmala alkaloid biosynthesis in *B. caapi*. Following THH formation, the *S*-tetrahydroharmine (THH) enantiomer is selectively oxidized by harmaline synthase (HS), whereas *R*-THH accumulates due to its poor substrate compatibility with BcaHS. Harmaline is subsequently oxidized by an unknown oxidase to form harmine, which accumulates at levels comparable to *R*-THH (Figure S21). This metabolite distribution is consistent with a non-enzymatic cyclization step that generates racemic THH, and supported by in vitro assays using BcaHS and either racemic THH or THH accumulated in *B. caapi* (**A** and **C**). In this proposed pathway, acetaldehyde may be generated through pyruvate decarboxylase activity; however, the biosynthetic origin and flux of this intermediate in *B. caapi* remain to be determined and will require further investigation. See also Figures S21 and S24–S28.

Consistent with these results, in vitro reactions showed acetaldehyde-dependent cyclization of tryptamine derivatives to THH under the tested aqueous conditions, accelerated by acetaldehyde concentration and, non-specifically, by proteins such as bovine serum albumin or yeast lysate (Figures 6F, S26A, S26B, and S27). Addition of BcaHS then converted THH to harmaline (Figures 7A and S28). BcaHS showed negligible activity toward native THH extracted from *B. caapi* leaves, a result consistent with, but not by itself establishing, enrichment of the *R*-enantiomer in planta (Figure 7C). Thus, these data support a model in which acetaldehyde-dependent, non-enzymatic cyclization may contribute to THH formation in planta, followed by selective enzymatic oxidation of the *S*-enantiomer by BcaHS (Figures 7D and 7E); the terminal oxidation of harmaline to harmine remains unresolved, as none of the remaining candidates performed this step. Because candidate selection was guided by expression and homology, these experiments do not exclude an additional, untested enzymatic contribution to the cyclization step.

## Discussion

Here we report haplotype-phased chromosome-scale genome assemblies for *P. viridis* and *B. caapi*, two species that are functionally linked in ayahuasca yet genomically distinct: a structurally complex hexaploid and a diploid, respectively. The high collinearity among the six *P. viridis* monoploid sets and the almost perfect one-to-one correspondence with *Coffea* chromosomes support extensive retention of homologous genome structure and are consistent with, but not fully diagnostic of, an autopolyploid interpretation^23,39^. The well-characterized allotetraploidy of cultivated *Coffea arabica*^39^, which retains extensive subgenomic synteny derived from closely related parental species, provides a useful Rubiaceae comparison, while *P. viridis* provides a tractable system for studying the genomic consequences of hexaploidy. Accordingly, while the current data favor an autopolyploid interpretation, allopolyploidy involving closely related parental lineages cannot be excluded, should subgenome dominance or appreciable post-WGD diploidization be inconspicuous.

The centromeres of both species are dominated by the same retrotransposon lineage, CRM of the Ty3/*Gypsy* superfamily. The younger age of centromeric CRM copies is consistent with ongoing centromere-biased amplification or preferential retention^25,28,40^. In *B. caapi*, however, centromeres additionally accumulate specific LINE families and chromosome-specific satellites, some sharing high sequence homology with CRM LTRs, likely derived from them^28^. Similar LTR-derived centromeric satellite repeats have also been reported in pentaploid dogroses^29^. Satellitization appears to have occurred at least twice in *B. caapi*, and probably three times: TRC1 derives from a CRM inter-domain region, whereas TRC2 and TRC4 derive from different LTR-containing templates and share sequence only within their LTR-derived segments. In all three satellite-containing centromeres, CENH3 enrichment was higher over the TRC arrays than over CRM elements, whereas CRM and LINE elements showed no consistent difference across chromosomes. This suggests that, once amplified, a CRM-derived satellite can become a preferred substrate for CENH3 deposition, as reported for TE-derived satellite repeats in pentaploid dogroses^29^. Centromeric LINEs are rare in plants, with clear examples reported in sunflower^30^ and poplar^31^, so the *B. caapi* CRM + LINE + satellite configuration extends the growing picture of centromeres as dynamic, epigenetically defined compartments capable of recruiting distinct repeat lineages. Poplar, one of the two other plants with reported centromeric LINEs, also belongs to the Malpighiales, raising the possibility that centromeric LINE accumulation is more frequent in this order.

The two plants elaborate tryptamine through different biosynthetic organizations. In *P. viridis*, DMT is produced by a minimal pathway in which the single multifunctional methyltransferase PviTNMT performs both *N*-methylations, strongly favoring the first step; pathway output may therefore be strongly influenced by substrate availability. The same two enzymes were independently reported while this work was in progress^15^, and the agreement between studies strengthens the conclusion that a compact, enzyme-constrained route suffices for DMT. The functional data do not require distinct methyltransferases for the two N-methylation steps, pointing to enzyme multifunctionality rather than separate enzymes for each reaction. The retention and expression of PviTNMT homologous copies across the six chromosome sets remain to be resolved. Phylogenetically, PviTNMT falls within a clade of Rubiaceae/Apocynaceae methyltransferases associated with monoterpenoid indole and ipecac alkaloid metabolism and is only distantly related to barley gramine synthase, consistent with repeated, independent evolution of tryptamine N-methylation across plants, amphibians, fungi and humans^15,41,42^.

In *B. caapi*, by contrast, β-carboline biosynthesis combines enzymatic steps with chemistry that can proceed non-enzymatically. Dedicated enzymes (BcaTDC, BcaT6H, Bca6OMT) build 6-methoxytryptamine, but the key Pictet-Spengler cyclization to THH can proceed with acetaldehyde without an added *B. caapi* cyclase, yielding a racemate in the experimental systems; BcaHS then selectively oxidizes *S*-THH to harmaline, while *R*-THH is not oxidized in the assays; the terminal conversion of harmaline to harmine remains unresolved. This organization differs from earlier proposals invoking a β-carboline-carboxylic-acid intermediate in *Peganum harmala*^35^, *Peganum multisectum*^43^ and *Elaeagnus angustifolia*^35,37^, and it provides a plausible biochemical basis for the co-accumulation of THH and harmine in the plant. The methoxy group installed by Bca6OMT increases indole nucleophilicity and accelerates the non-enzymatic cyclization under the tested conditions, illustrating how an enzyme can “prime” a non-enzymatic step. That a culturally and pharmacologically central metabolite may depend partly on non-enzymatic chemistry is consistent with the view that metabolic complexity need not scale with gene number or genome size, but can emerge from the interplay of enzyme properties, substrate availability and intrinsic reactivity^44,45^. The *P. viridis* pathway is tightly enzyme-controlled, whereas the *B. caapi* pathway includes more flexible chemistry. Thus, the two species exhibit contrasting pathway organizations within a single ethnobotanical system, but the present comparison cannot distinguish whether these differences are consequences of their different ploidy states or of their distant phylogenetic relationship.

Beyond its interest as a case of ploidy divergence, shared centromere architecture and biochemical complementarity, ayahuasca is a sacrament. For many Indigenous Amazonian peoples and for the members of religious communities such as the UDV and Santo Daime, *B. caapi* and *P. viridis* are sacred beings central to healing, worship and community life, and their ceremonial use is a protected expression of religious freedom in several jurisdictions^1,2^. In presenting the molecular foundations of these plants and of the alkaloid biosynthesis that characterizes them, we are conscious that a genome can amount to more than data when it is part of a living heritage. We therefore frame this work as a resource whose stewardship should respect the communities for whom these plants are sacred and the governance obligations associated with such material.

Beyond this cultural context, the genomes, population data and enzymes reported here are resources for several research areas. The six collinear monoploid sets of *P. viridis* provide a system for studying the genomic consequences of hexaploidy, and the CRM-, LINE- and satellite-containing centromeres of *B. caapi* add a plant case to comparative studies of centromere evolution. The resequencing panel provides a genetic reference for *B. caapi* accessions from Brazil. Finally, the two-plant pharmacology of ayahuasca, in which MAO inhibition by β-carbolines enables oral DMT activity, is now being pursued therapeutically, with early controlled and open-label trials reporting rapid antidepressant effects of ayahuasca and of standardized DMT in treatment-resistant depression, alongside investigation in anxiety, addiction and PTSD^5,6,46^. The biosynthetic enzymes reported here, including a compact DMT route and a partly spontaneous β-carboline route, provide defined parts for studying and reconstituting the production of these molecules in heterologous systems.

Several questions now become experimentally tractable: the identity of the oxidase that converts harmaline to harmine, the enantiomeric composition of THH in planta and the retention and expression of PviTNMT copies across the six monoploid sets of *P. viridis*. In conclusion, the integration of chromosome-scale genomics, population analysis and functional biochemistry provides a molecular framework for the study of ethnobotanically important plants while recognizing their cultural and ecological contexts.

### Limitations of the study

The autopolyploid interpretation of *P. viridis* rests on collinearity among the six monoploid sets, one-to-one correspondence with *C. canephora* chromosomes and the absence of subgenome-specific k-mer groups. None of these observations is diagnostic, and allopolyploidy involving closely related parental lineages cannot be excluded without independent evidence of parental origin. The TCN and CPR reference genomes each derive from a single accession sampled from the same plantation, and the population panel comprises 135 accessions from 18 localities in Brazil, so the genetic structure reported here may not capture the full diversity of wild and cultivated *B. caapi*.

Candidate enzymes were selected by expression level and sequence homology, and an additional enzymatic contribution to the Pictet-Spengler cyclization cannot be excluded. The enantiomeric composition of THH in *B. caapi* leaves was inferred from the lack of BcaHS activity toward native THH rather than measured by chiral separation, and the enzyme that oxidizes harmaline to harmine remains unidentified. The source and concentration of acetaldehyde in *B. caapi* tissues were not determined, so the relative contributions of non-enzymatic and enzymatic steps in planta remain unresolved.

## Resource availability

### Lead contact

Requests for further information and resources should be directed to and will be fulfilled by the lead contact, André Marques.

### Materials availability

Plasmids generated in this study are available from the lead contact upon request. Plant material was collected under the access and benefit-sharing authorizations required by Brazilian law (SisGen); its transfer to third parties is subject to these authorizations and may require a material transfer agreement.

### Data and code availability

Raw sequencing reads (HiFi, Hi-C, RNA-seq, Iso-Seq, ChIP-seq, resequencing) and annotations have been deposited at ENA Project PRJEB123950 and will be publicly available as of the date of publication. Custom scripts used for centromere boundary refinement and for the geographic mapping of genetic clusters and ancestry have been deposited at GitHub page: https://github.com/RuiyiQiu/ayahuasca. Any additional information required to reanalyze the data reported in this paper is available from the lead contact upon request.

## Acknowledgments

We are grateful to M. Geraldo Franco for providing us the *P. viridis* and *B. caapi* material used for generating the reference genomes. We thank our friends who allowed and contributed to the collections in the Xingu: Duarte Guerra, Karin Juruna and Yakarawa Juruna; we are grateful for all the good companionship that provides peace. To Prof. Carolina Joana da Silva (in memoriam) for the enriching suggestions and inspiration in the development of the work, always innovating with her ideas, providing us with professional and personal development. We thank the Centro Espírita Beneficente União do Vegetal (CEBUDV) for logistical and operational support of the field collections. We are grateful to José B. B. Figueiredo (Universidade Federal de Roraima) and Marciel J. Ferreira (Laboratório de Silvicultura, Universidade Federal do Amazonas, UFAM) for guidance on field sampling and for logistical support during the collections.

RQ, TN and AM thank the Max Planck Society. SGAM received support from the Canada Graduate Research Scholarship. YQ acknowledge support from the Natural Sciences and Engineering Research Council of Canada (NSERC) Discovery Grant, ResearchNB Research Professional Initiative and Research Assistantship Initiative grants. This work was supported by the Fundação de Amparo à Pesquisa do Estado do Amazonas (FAPEAM, POSGRAD program; grants 062.00133/2020 and 062.00148/2020).

## Author contributions

Conceptualization: CA, YQ, AM; Data curation: RQ, CA, YQ, AM; Formal analysis: RQ, SGAM, TN, JOP, MBR, DARD; Funding acquisition: CA, YQ; Investigation: RQ, CA, YQ, AM; Methodology, Resources: MB, ABCA, AOA, JS, CRC; Supervision: RQ, JTL, GS, VAA, CA, YQ, AM; Validation, Visualization, Writing – original draft: RQ, CA, YQ, AM and Writing – review & editing: RQ, GS, VAA, CA, YQ, AM.

## Declaration of generative AI and AI-assisted technologies in the writing process

During the preparation of this work, the authors used Claude (Anthropic) to format the manuscript and figures. After using this tool, the authors reviewed and edited the content as needed and take full responsibility for the content of the publication.

## Methods

### Experimental model and study participant details

#### Plant material and ethics

Leaf tissue from a single, well-defined individual of *Psychotria viridis* was collected to minimize chimerism. For *Banisteriopsis caapi*, stem/vine material of the cultivated varieties Tucunacá (TCN) and Caupuri (CPR) was obtained from representative accessions. For the population panel, leaf tissue from 135 *B. caapi* individuals was collected from 18 localities in Brazil (Table S3); the TCN and CPR reference genotypes were analyzed as two additional samples (137 in total). Plant material was obtained with the access and benefit-sharing authorizations required under Brazilian law (SisGen registration no. AB63C5D, A9B4381, ABF4610, AF3C936) and with the consent of the communities and collectors acknowledged in the manuscript.

### Method details

#### Nucleic acid extraction and sequencing

High-molecular-weight DNA was extracted using modified CTAB and Nanobind-based protocols, with size selection enriching fragments >50 kb. Total RNA was extracted with Qiagen Plant RNA kits, DNase-treated, and quality-checked on a Bioanalyzer. PacBio HiFi long-read sequencing was performed to approximately 90× coverage per monoploid genome equivalent for *P. viridis*, and approximately 22×–23× coverage per haplotype for *B. caapi*. Hi-C libraries (Arima Genomics) were prepared for chromosome-scale scaffolding and haplotype phasing. Illumina short-read RNA-seq and PacBio Iso-Seq libraries were prepared from multiple tissues (young leaf, mature leaf, <u>old leaf,</u> stem, root). For population genomics, Illumina whole-genome resequencing was performed with a mean depth of approximately 6× mean coverage per accession. Sequencing libraries were sent to BGI Genomics (Hong Kong) for sequencing.

#### Genome assembly and phasing

Genome characteristics (size, heterozygosity, repeat content, ploidy) were estimated with GenomeScope 2.0 and Smudgeplot^47,48^. For *P. viridis*, genome assembly were performed using hifiasm in Hi-C phasing mode^49^, generating six reconstructed monoploid chromosome sets. Hi-C scaffolding used HapHiC^50^ and C-Phasing^51^ followed by manual curation to correct misjoins. The six monoploid sets (g1–g6) were ordered arbitrarily by decreasing total sequence length. For *B. caapi*, the Caupuri (CPR) reference assembly was generated from HiFi and Hi-C data using hifiasm in Hi-C phasing mode, with Hi-C scaffolding performed using HapHiC, yielding two phased haplotypes. The Tucunacá (TCN) assembly was generated from HiFi reads using hifiasm, and the resulting contigs were scaffolded onto the CPR reference assembly using RagTag^52^. Assembly completeness was assessed with BUSCO^53^ (embryophyta_odb12 lineage dataset) and base and phasing accuracy with Merqury^54^. Structural variation between haplotypes and varieties was identified with MUMmer^55^ and SyRI^56^.

#### Repeat and gene annotation

Repeats were annotated with the RepeatExplorer2 TideCluster, DANTE and DANTE-LTR pipelines, identifying and classifying LTR retrotransposons, non-LTR elements (LINEs, SINEs), DNA transposons and satellites^57^. A comprehensive repeat library was additionally constructed using EDTA^58^, and assembly continuity within repeat-rich regions was assessed with the LTR Assembly Index (LAI). Gene structure was predicted with Helixer^59^. Distribution of genomic features was visualized using R package *circlize*^60^.

#### Synteny and whole-genome duplication

Genomic synteny and structural rearrangements were characterized and visualized using SyRI^56^. Haplotypes were aligned with minimap2 (-ax asm20 --eqx) before SyRI analysis; SNP densities in 500-kb windows, normalized to aligned sequence, and inversion lengths were summarized from the SyRI output. Protein-coding gene collinearity and synteny were analyzed with GENESPACE^61^, using *Coffea canephora* as the diploid Rubiaceae reference^39^. For *B. caapi*, the chromosome-scale assembly of *Malpighia emarginata* (NRA309_r1.0)^27^ was used as the Malpighiaceae reference. Self-synteny dot plots, fractionation bias and K*s* distributions of syntenic gene pairs were obtained with CoGe SynMap^62^ and FractBias^63^ for the *P. viridis* and *B. caapi* self-comparisons and for *P. viridis* versus *C. canephora*, and were used to assess whole-genome duplication. Subgenome-specific k-mers among the 66 *P. viridis* pseudochromosomes were assessed with SubPhaser^64^ (k = 19, minimum k-mer frequency q = 50, fold-enrichment threshold f = 2).

#### Population structure

Resequencing reads were mapped to the *B. caapi* TCN reference genome with BWA-MEM, and variants were called with bcftools. Biallelic SNPs were retained using VCFtools (indels excluded, minor allele frequency ≥ 0.05, genotype call rate ≥ 90%, quality ≥ 30). Population structure was analyzed via PCA with k-means clustering using VCF2PCACluster^65^, and phylogenetic network reconstruction using VCF2Dis^66^ and SplitsTree^67^. Ancestry proportions were estimated using sNMF in the R package LEA^68^ based on PLINK-formatted genotypes^69^. Clustering and ancestry were mapped geographically using custom R scripts.

#### Slides preparation and Fluorescent in situ Hybridization (FISH)

Meristematic tissue from young roots of *B. caapi* and *P. viridis* was pre-treated with 2 mM 8-Hydroxyquinoline solution for 4 h at 18 °C to accumulate metaphases, and then fixed in Carnoy solution [Methanol:Acetic Acid, 3:1 (v/v)] for 2–24 h at room temperature, before being used for slide preparation or kept at −20 °C until use. Slides were prepared using an air-drying technique with a 30 min bath in 60% acetic acid for cytoplasm removal. Briefly, the meristems were treated with an enzymatic solution containing 2% cellulase R-10 Onozuka (Serva), 2% pectolyase Y-23 (Serva) and 20% pectinase (Sigma) in 1× citrate buffer (pH 4.8) in a humid chamber for 1 h at 37 °C. The material was then macerated in one drop of fresh cold Carnoy solution, spread and air-dried, incubated in 60% acetic acid, and left to dry at room temperature. The best slides were selected by epi-fluorescence microscopy after mounting with 2 µg/mL DAPI in Vectashield H-100 antifade solution (Vector Laboratories). For FISH experiments, oligo probes were used for the *Arabidopsis thaliana* telomeric motif (TTTAGGGn) labelled in Cy5, and oligo pools described by Waminal et al.^70^ for the angiosperm 35S and 5S rDNA sites labelled in FAM and Cy3, respectively. The hybridization mix consisted of 50% formamide, 2× SSC (Saline Sodium Citrate), 10% dextran sulphate, and 25 ng of each probe (15 µL per slide). Slides were denatured at 75 °C for 5 min and hybridized at 37 °C in a dark humid chamber for at least 18 h. After hybridization, slides were subjected to stringency washes in 2× and 0.1× SSC at 42 °C, yielding a final stringency of ∼76%. Slides were mounted with 2 µg/mL DAPI in Vectashield antifade medium (Vector Laboratories) and imaged using a Zeiss fluorescence microscope equipped with a Zeiss Axio Imager 2 camera system and ZEN Blue software (v.3.1). Selected images were adjusted for brightness and contrast in Adobe Photoshop software.

#### CENH3 ChIP-seq and centromere analysis

To detect the functional centromeres of accessions of both *B. caapi* and *P. viridis*, we generated a specific polyclonal antibody raised in rabbit against the respective CENH3 proteins (*B. caapi*: RIKHQATRSQARSNFCGV; *P. viridis*: RSKHPATRGGPSRRSSAN), previously identified by BLASTP using the CENH3 protein sequence of A. thaliana as a reference and synthesized by LifeTein. For the ChIP experiment, young leaves (around 2–5 g) of each plant were collected and cross-linked in 4% formaldehyde in 1× PBS on ice, with vacuum infiltration applied for 30 min, then quenched with 1 M glycine under vacuum infiltration at room temperature for 15 min. The material was then macerated in liquid nitrogen and chromatin was extracted according to Reimer and Turck^71^. After extraction, the chromatin was sonicated for 30 min using a Bioruptor (Diagenode) until fragments of ∼200–600 bp were obtained (30 s on; 30 s off; high mode). The sonicated chromatin was incubated overnight at 4 °C with 1 µg of each polyclonal antibody [anti-CENH3 antibody raised in rabbit and anti-histone H3 (Active Motif, 39064) raised in mouse]. Samples without primary antibodies were also incubated as input controls, and at least two experimental replicates were used for each ChIP combination. After incubation, Sepharose beads [anti-rabbit: rProtein A Sepharose FastFlow 50% slurry; anti-mouse: rProtein G Sepharose FastFlow 50% slurry (GE Healthcare)] were washed and added to each protein–antibody complex and incubated for at least 2 h at 4 °C with slow rotation (12 rpm). The chromatin was then eluted from the beads, followed by decrosslinking and DNA extraction. After quality assessment using a 4200 TapeStation System (Agilent Technologies), the samples were submitted for 150 bp paired-end Illumina sequencing.

Chromatin immunoprecipitation sequencing was performed with antibodies against CENH3 alongside H3 as a control, with two biological replicates each. Reads were mapped to the reference assemblies using bowtie2^72^, and peaks were called with MACS3^73^. The CENH3/H3 log2 enrichment signal was calculated with deepTools^74^ bamCompare (parameters: --normalizeUsing RPKM --scaleFactorsMethod None --operation log2). For quantification, the CENH3/H3 log2 enrichment signal and repeat element coverage were computed in 1-kb non-overlapping windows using bedtools^75^. Each window within centromeres was assigned to the element class with non-zero coverage, with TRC taking priority where it co-occurred with CRM or LINE, and CENH3/H3 log2 ratios were compared between element classes using the Wilcoxon rank-sum test in R. For centromere metaplots, CENH3/H3 enrichment and LINE, CRM and TRC coverage were summarized in 1-kb windows across each centromere and the flanking 5% of chromosome length on either side. To define functional centromere regions at high resolution, we used custom scripts to refine MACS3 peaks: MACS3 peaks were first merged within 100 kb gaps to retain the largest merged block per chromosome as a coarse candidate centromeric region; then this region was refined by scanning the continuous CENH3/H3 log2 signal with a sliding window. For both *B. caapi* varieties, boundaries were refined using a 50-kb window, 5-kb step, log2 threshold of 0.2, and 5 consecutive windows; for *P. viridis*, a 100-kb window, 10-kb step, log2 threshold of 0.1, and 3 consecutive windows were used. Genome-wide CRM phylogenetic trees were constructed by aligning CRM sequences with MAFFT^76^, trimming alignments with trimAl^77^, inferring trees with FastTree2.2^78^, and visualizing them with iTOL (https://itol.embl.de/). CRM LTR insertion ages were estimated from LTR identity provided by RepeatExplorer2 DANTE_LTR. LINE families were grouped by CD-HIT clustering (threshold: 40% amino acid identity), and enrichment was tested using odds ratios with Fisher’s exact test followed by Benjamini-Hochberg correction for multiple testing. Centromere self-similarity was visualized with re-DOT-able (https://github.com/s-andrews/redotable), and genomic tracks with pyGenomeTracks^79^.

#### Transcriptomics and pathway inference

RNA-seq reads were quantified with Salmon^80^; differential expression used DESeq2^81^. Iso-Seq reads were processed to full-length transcripts to refine gene models. Co-expression modules were identified with WGCNA^82^. Candidate metabolic genes were prioritized by homology, tissue-specific and co-expression patterns, genomic context and predicted enzyme domains.

#### Phylogenetic analysis

Protein sequences were aligned using ClustalW^83^. The evolutionary history was inferred in MEGA11^84^ using the Maximum Likelihood method and the JTT matrix-based model. Bootstrap support values (100 replicates) are shown next to the branches. Initial trees for the heuristic search were obtained by applying Neighbor-Join and BioNJ algorithms to a matrix of pairwise distances estimated using the JTT model, selecting the topology with the superior log likelihood value. Branch lengths are measured in the number of substitutions per site. The plant species and protein sequences used in the analyses are provided in Data S1.

#### Homology modeling, docking and molecular dynamics

Homology models of PviTNMT, BcaHS, BcaT6H and Bca6OMT were built and substrates were docked into their active sites using the Molecular Operating Environment (MOE) 2022.02. Docking poses were inspected manually to identify configurations conducive to the relevant catalytic reactions, and the pose with the lowest docking energy and a biochemically plausible orientation was selected. The stability of the selected S- and R-THH poses in BcaHS was assessed by molecular dynamics simulations, and ligand stability was evaluated from the RMSD over the simulation (Figure S24).

#### Chemical standards

Tryptamine, 4- and 5-hydroxytryptamine, racemic tetrahydroharmine, harmaline, harmine, harmane, harmol and harmalol were purchased from Cayman Chemical (Ann Arbor, MI, USA). Deuterated (d4) tryptamine and 6-hydroxytryptamine were purchased from Toronto Research Chemicals (Vaughan, ON, Canada). 6-methoxytryptamine was purchased from Santa Cruz Biotechnology (Dallas, TX, USA).

#### Cloning

All *P. viridis*, *B. caapi* and *R. marina* cDNA sequences were synthesized and subcloned into pESC-Ura/Leu vectors within *BamH*I/*Sal*I sites or in pESC-His vector within *EcoR*I/*Not*I sites (Twist Bioscience, South San Francisco, CA, USA). The resulting vectors were mobilized to *Saccharomyces cerevisiae* strain BY4741 (MATα his3Δ1 leu2Δ0 met15Δ0 ura3Δ0 YPL154c::kanMX4) using a standard lithium acetate/polyethylene glycol transformation procedure. For gene expression in *N. benthamiana*, the sequences were amplified using primers (1/2, from pESC-Ura/Leu constructs) or primers (3/4, from pESC-His constructs) and subcloned into pEAQ-HT vector within *Age*I site, using standard Gibson assembly (NEBuilder® HiFi DNA Assembly, New England Biolabs, Ipswich, MA, USA). The resulting vectors were introduced into *A. tumefaciens* strain LBA4404. For *PviTNMT* expression in *E. coli*, the sequence was amplified using primers (5/6) and subcloned into pET30b+ vector within *BamH*I/*Sal*I sites using Gibson assembly. The vector was introduced into *E. coli* BL21(DE3) cells.

#### Yeast culturing and biotransformation

Single yeast colonies carrying various vectors were inoculated in 1 mL synthetic complete (SC) medium with 2% (w/v) glucose, and incubated at 30 °C, 200 rpm overnight. The yeasts were pelleted by centrifugation, washed once with water, resuspended in 1 mL SC medium with 2% (w/v) galactose, and incubated at 30 °C, 200 rpm overnight. For alkaloid detection, cultures were mixed with equal volumes of methanol, centrifuged, and the supernatants were analyzed by LC-MS/MS.

#### Transient expression in N. benthamiana and biotransformation

Single colonies of *A. tumefaciens* strain LBA4404 carrying various vectors were inoculated in 1 mL LB medium supplemented with 0.1 mg/mL kanamycin and 0.02 mg/mL rifampicin at 28 °C, 200 rpm overnight. Cells were pelleted, resuspended in infiltration buffer (10 mM MES pH 5.6, 10 mM MgCl2, 0.2 mM acetosyringone) to an OD₆₀₀ of 0.8, and incubated at 28 °C, 200 rpm for 2-3 h. Suspensions were infiltrated individually or mixed in equal volumes into the abaxial side of *N. benthamiana* leaves using a syringe. Plants were maintained in a growth chamber (150 µmol/m²/s, 16 h light/8 h dark, 26 °C) for 3-5 days. Leaves were harvested, homogenized in 0.5 mL methanol using small pestles in 1.5 mL microcentrifuge tubes. After centrifugation, the supernatants were analyzed by LC-MS/MS. For tetrahydroharmine biosynthesis, substrate solutions (0.26 mM 6-methoxytryptamine, 2.3 mM acetaldehyde or ethanol in 20 mM Tris HCl pH 7.5) were infiltrated into *N. benthamiana* leaves. Samples were harvested after 1 day and processed as described above.

#### Isotope labeling in B. caapi leaves

Young *B. caapi* leaves (5-6 cm wide, 8-10 cm long) were cut into 1 × 1 cm pieces and incubated in 0.1 mL 20 mM Tris pH 7.5 containing 0.53 mM d4-tryptamine and/or 2.3 or 23 mM acetaldehyde/ethanol in 1.5 mL microcentrifuge tubes, covered with parafilm. Samples were maintained in a growth chamber under the same conditions as above. All experiments were performed in triplicate. After 1 day, 0.1 mL methanol was added, and tissues were homogenized with small pestles. After centrifugation, the supernatants were analyzed by LC-MS/MS.

#### Protein isolation from yeast and purification from E. coli

For yeast expressing BcaHS or harboring an empty vector, 0.5 mL of an overnight culture was used to inoculate 50 mL SC medium containing 2% (v/v) glucose. Cultures were incubated at 30 °C, 200 rpm overnight. Cells were pelleted, washed once with water, and resuspended in 50 mL SC medium containing 2% (v/v) galactose, followed by incubation at 30 °C, 200 rpm overnight. Cells were pelleted and resuspended in 1 mL sample buffer (20 mM Tris pH 7.5, 100 mM NaCl, 10% (v/v) glycerol). Cell lysis was performed using 1 mm glass beads in a 2 mL microcentrifuge tube with a BeadBug™ microtube homogenizer (30 Hz, 180 s; Benchmark Scientific, Sayreville, NJ, USA) for three cycles, with 2 min cooling intervals on ice at 4 °C. The slurry was briefly centrifuged to remove glass beads, followed by centrifugation at 20,000 g for 20 min at 4 °C to obtain the soluble protein fraction.

For PviTNMT purification, *E. coli* BL21(DE3) cells expressing N-terminal His-tagged proteins were grown in LB medium to an OD₆₀₀ of 0.7, then induced with 0.1 mM isopropyl β-D-1-thiogalactopyranoside (IPTG) at 15 °C, 200 rpm overnight. Cells were harvested and lysed by sonication in ice-cold sample buffer (20 mM Tris-HCl, pH 7.5, 100 mM NaCl, 10% (v/v) glycerol) and centrifuged at 10,000 g for 10 min at 4 °C to obtain the soluble protein fraction. The proteins were purified using Ni-NTA affinity chromatography (Cytiva, Marlborough, MA, USA) and desalted into sample buffer using a PD-10 desalting column (Cytiva). Purified proteins were stored at −80 °C until further use.

#### In vitro assays and enzyme kinetics

For testing tetrahydroharmine biosynthesis or BcaHS activity, the in vitro assays (100 µL) included 20 mM Tris HCl pH 7.5, 250 µg yeast crude lysate or 250 µg bovine serum albumin (Bio Basic, Markham, ON, Canada), and various amounts of substrates or total *B. caapi* leaf extracts. Reactions were incubated at 30 °C overnight for completion, and then quenched with an equal volume of methanol. Samples were centrifuged, filtered, and analyzed by LC-MS/MS.

For PviTNMT kinetics, each reaction (50 µL) contained 20 mM Tris-HCl (pH 7.5), 0.25 µg of purified PviTNMT, 0.06 mM S-adenosyl methionine, and either tryptamine or monomethyltryptamine substrate at various concentrations. Reactions with tryptamine were incubated at 30 °C for 15 min, whereas reactions with monomethyltryptamine were incubated for 2 h. All reactions were quenched by adding an equal volume of methanol, followed by centrifugation, filtration, and LC-MS/MS analysis. Kinetic parameters were determined by fitting the data to a non-linear regression model using GraphPad Prism version 10.4.2.

#### LC-MS/MS

The samples were analyzed using the Ultivo Triple Quadrupole LC-MS/MS system from Agilent (Santa Clara, CA, USA), equipped with an Agilent Poroshell EC-C18 column (100×4.6 mm, particle size 2.7 *μ*m) as well as a photodiode array detector and a mass spectrometer. The following solvent systems were used: solvent A, water (0.1% formic acid); solvent B, acetonitrile (0.1% formic acid). The following linear elution gradient was used: 0 min 99% A, 1% B; 0.5 min 99% A, 1% B; 10 min 1% A, 99% B;12 min 1% A, 99% B; 13 min 99% A, 1% B; 18 min 99% A, 1% B. The flow rate was held constant at 0.6 mL/min. The photodiode array detector recorded from 200 to 500 nm. The mass spectrometer was operated with the gas temperature at 300 °C and gas flow of 12 L/min. The capillary voltage was 4 kV over m/z 100 to 1000 with a 500 ms scan time, and the fragmentor voltage was 45 V with positive polarity. MS/MS acquisition used a fragmentor voltage of 45 V and collision energies of 20 or 30 V in positive-ion mode.

To separate harmaline and harmine properly for quantification, the samples were analyzed using an Avantor® ACE® UltraCoreTM SuperC18^TM^ column (2.5 μm, 50 × 3 mm) instead. The following solvent systems were used: solvent A, methanol: acetonitrile: ammonium acetate 1 M, water at 29:71:2:398; solvent B, methanol: acetonitrile: ammonium acetate 1 M: water at 130:320:0.25:49.7. The following linear gradient (8 min run, 0.6 mL/min) was used: 0 min 80% A, 20% B; 0.5 min, 80% A, 20% B; 5 min 1% A, 99% B; 5.8 min 1% A, 99% B; 6.5 min 80% A, 20% B; 8 min 80% A, 20% B. The photodiode array detector recorded from 200 to 500 nm. MS/MS acquisition used a gas temperature of 300 °C, gas flow of 10 L/min, capillary voltage of 4 kV, fragmentor voltage of 45 V, and collision energy of 30 V in positive-ion mode.

#### Quantification and statistical analysis

Statistical tests, the exact value of n and what n represents are reported in the Results and figure legends. Insertion ages of centromeric and non-centromeric CRM LTRs were compared with Mann–Whitney U tests. Enrichment of centromeric copies among CRM clades was tested with one-sided Fisher’s exact tests, and enrichment of centromeric LINE copies among CD-HIT clusters with Fisher’s exact tests; in both cases, p values were adjusted for multiple testing with the Benjamini–Hochberg method. CENH3/H3 log2 ratios of centromeric 1-kb windows assigned to different repeat classes were compared with two-sided Wilcoxon rank-sum tests. Enzyme kinetic parameters were estimated by non-linear regression of the Michaelis–Menten model in GraphPad Prism version 10.4.2. Metabolite levels are reported as mean ± standard deviation of biological replicates, with n given in the corresponding figure legends.

## Supplemental information

Document S1. Figures S1–S28 and Tables S1, S2, and S4–S8

Table S3. Collection metadata for all *Banisteriopsis caapi* samples included in the population genetic analysis, related to Figure 4 and STAR Methods.

Data S1. Amino acid sequences used in phylogenetic analysis, related to Figures S16 and S23 and STAR Methods.

