## Supplementary materials for "Ayahuasca plant genomes reveal retrotransposon-derived centromeres and contrasting routes to DMT and β-carbolines"

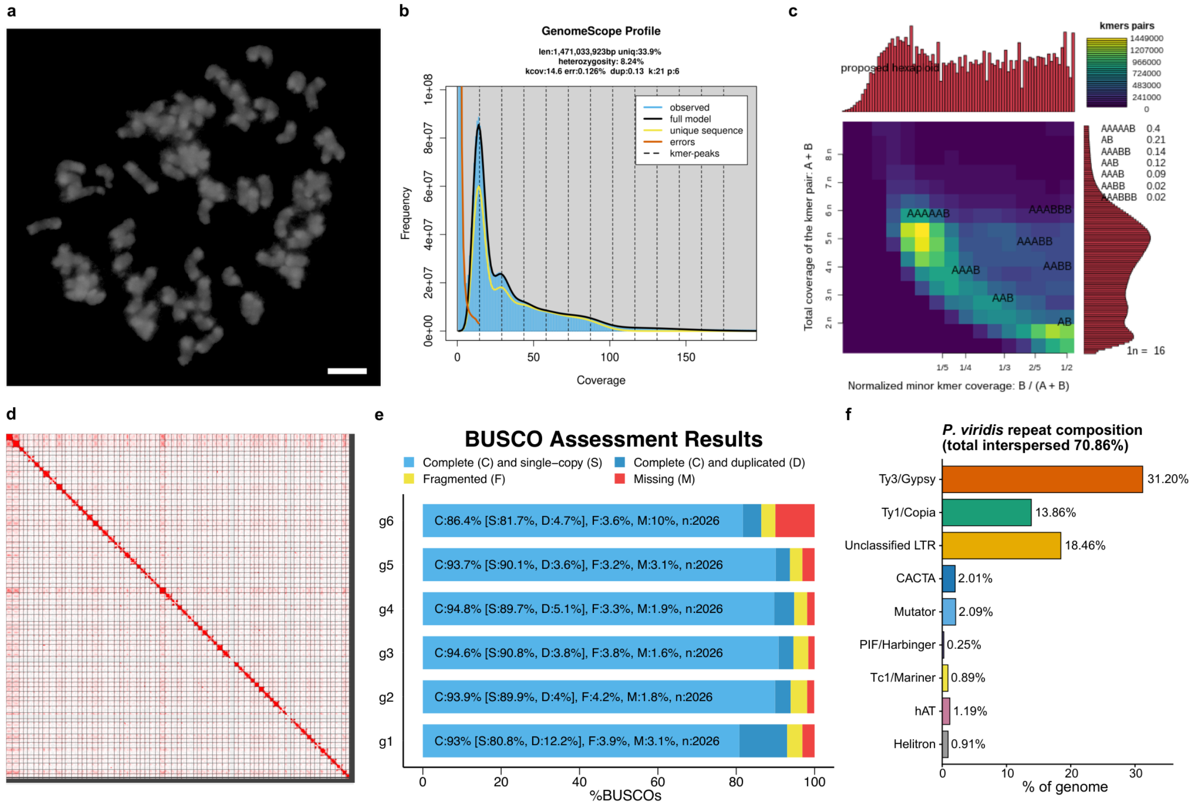


**Figure S1. Genome assembly assessment of *Psychotria viridis*, related to Figure 1 and Table S1. (A)** Chromosomes of *P. viridis* stained with DAPI. Bar corresponds to 5 µm. **(B)** Genome size and heterozygosity estimation based on *21*-mer distribution using GenomeScope2. **(C)** Ploidy and genome structure inference based on *21*-mer analysis using Smudgeplot. **(D)** Hi-C map of *P. viridis* genome assembly including 66 chromosomes. **(E)** BUSCO assessment results (embryophyta_odb12) of each monoploid set. **(F)** Interspersed repeat composition of the whole genome.

**a**

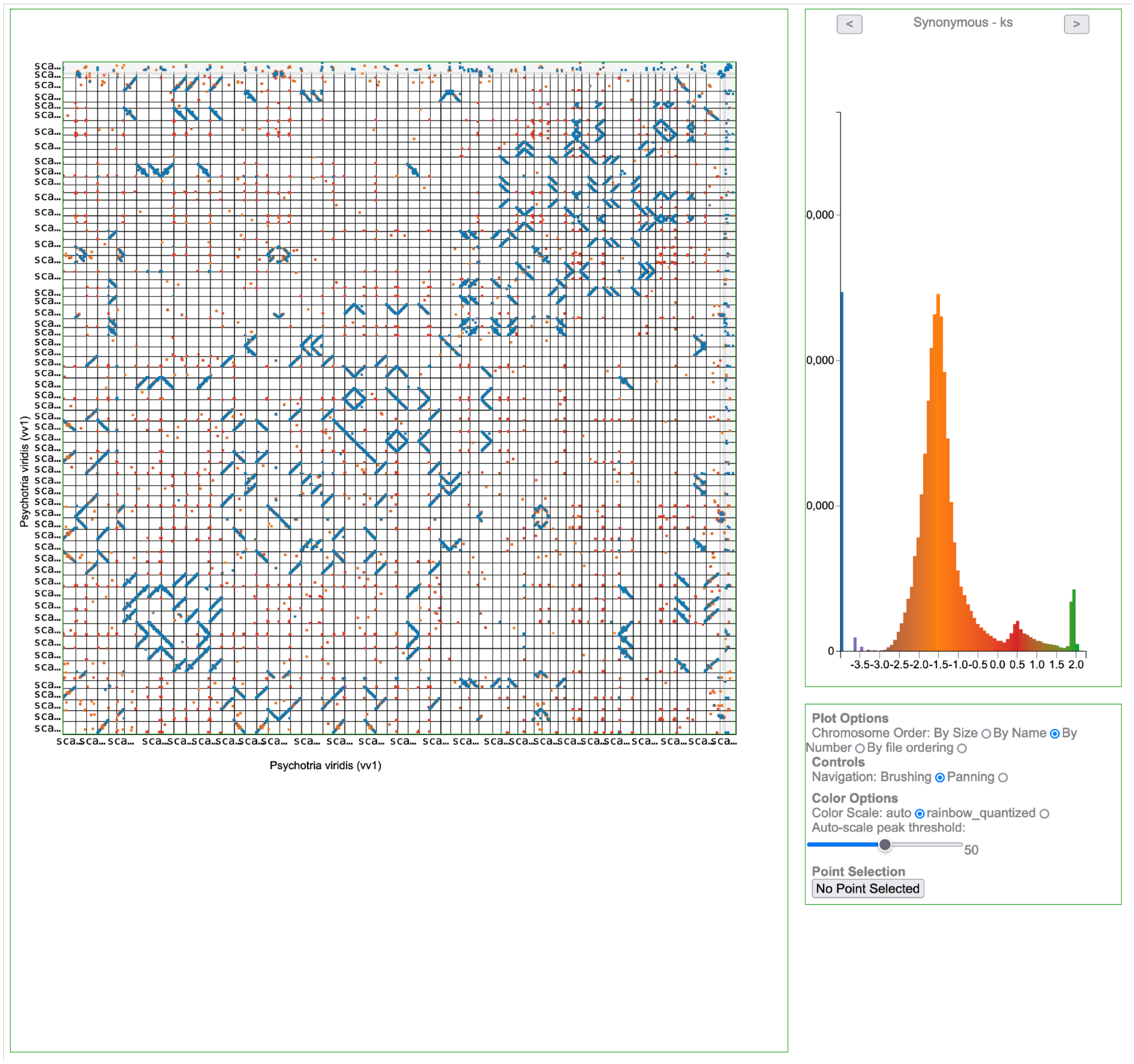

Psychotria viridis (vv1)
Psychotria viridis (vv1)
Synonymous - ks
-3.5-3.0-2.5-2.0-1.5-1.0-0.5 0.0 0.5 1.0 1.5 2.0
Plot Options
Chromosome Order: By Size By Name By Number By file ordering
Controls
Navigation: Brushing Panning
Color Options
Color Scale: auto rainbow_quantized
Auto-scale peak threshold:
50
Point Selection
No Point Selected

**b**

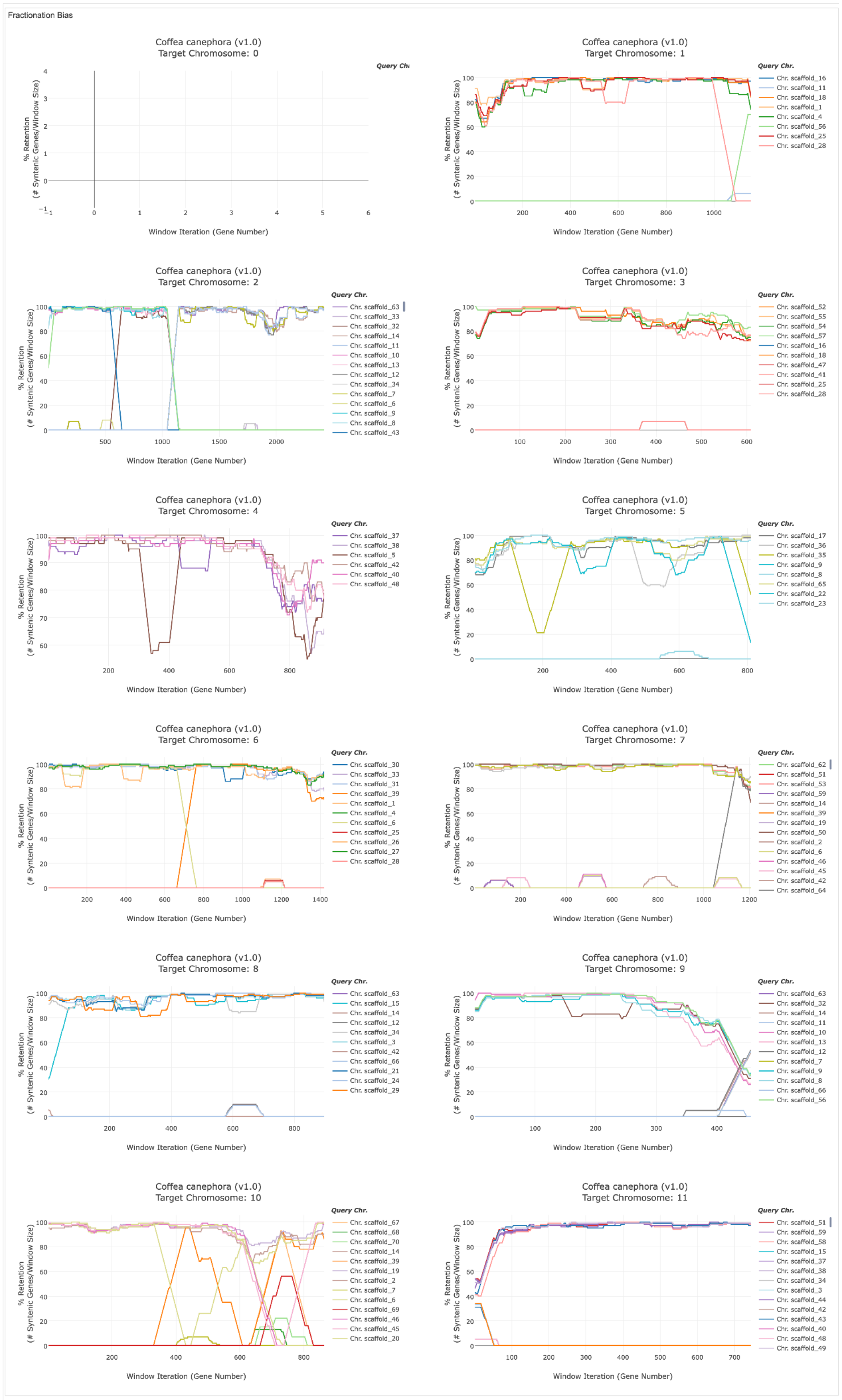

Fractionation Bias
Coffea canephora (v1.0) Target Chromosome: 0
Query Chr.
% Retention (# Syntenic Genes/Window Size)
Window Iteration (Gene Number)
Coffea canephora (v1.0) Target Chromosome: 1
Query Chr.
Chr. scaffold_16
Chr. scaffold_11
Chr. scaffold_18
Chr. scaffold_1
Chr. scaffold_4
Chr. scaffold_56
Chr. scaffold_25
Chr. scaffold_28
Coffea canephora (v1.0) Target Chromosome: 2
Query Chr.
Chr. scaffold_63
Chr. scaffold_33
Chr. scaffold_32
Chr. scaffold_14
Chr. scaffold_11
Chr. scaffold_10
Chr. scaffold_13
Chr. scaffold_12
Chr. scaffold_34
Chr. scaffold_7
Chr. scaffold_6
Chr. scaffold_9
Chr. scaffold_8
Chr. scaffold_43
Coffea canephora (v1.0) Target Chromosome: 3
Query Chr.
Chr. scaffold_52
Chr. scaffold_55
Chr. scaffold_54
Chr. scaffold_57
Chr. scaffold_16
Chr. scaffold_18
Chr. scaffold_47
Chr. scaffold_41
Chr. scaffold_25
Chr. scaffold_28
Coffea canephora (v1.0) Target Chromosome: 4
Query Chr.
Chr. scaffold_37
Chr. scaffold_38
Chr. scaffold_5
Chr. scaffold_42
Chr. scaffold_40
Chr. scaffold_48
Coffea canephora (v1.0) Target Chromosome: 5
Query Chr.
Chr. scaffold_17
Chr. scaffold_36
Chr. scaffold_35
Chr. scaffold_9
Chr. scaffold_8
Chr. scaffold_65
Chr. scaffold_22
Chr. scaffold_23
Coffea canephora (v1.0) Target Chromosome: 6
Query Chr.
Chr. scaffold_30
Chr. scaffold_33
Chr. scaffold_31
Chr. scaffold_39
Chr. scaffold_1
Chr. scaffold_4
Chr. scaffold_6
Chr. scaffold_25
Chr. scaffold_26
Chr. scaffold_27
Chr. scaffold_28
Coffea canephora (v1.0) Target Chromosome: 7
Query Chr.
Chr. scaffold_62
Chr. scaffold_51
Chr. scaffold_53
Chr. scaffold_59
Chr. scaffold_14
Chr. scaffold_39
Chr. scaffold_19
Chr. scaffold_50
Chr. scaffold_2
Chr. scaffold_6
Chr. scaffold_46
Chr. scaffold_45
Chr. scaffold_42
Chr. scaffold_64
Coffea canephora (v1.0) Target Chromosome: 8
Query Chr.
Chr. scaffold_63
Chr. scaffold_15
Chr. scaffold_14
Chr. scaffold_12
Chr. scaffold_34
Chr. scaffold_3
Chr. scaffold_42
Chr. scaffold_66
Chr. scaffold_21
Chr. scaffold_24
Chr. scaffold_29
Coffea canephora (v1.0) Target Chromosome: 9
Query Chr.
Chr. scaffold_63
Chr. scaffold_32
Chr. scaffold_14
Chr. scaffold_11
Chr. scaffold_10
Chr. scaffold_13
Chr. scaffold_12
Chr. scaffold_7
Chr. scaffold_9
Chr. scaffold_8
Chr. scaffold_66
Chr. scaffold_56
Coffea canephora (v1.0) Target Chromosome: 10
Query Chr.
Chr. scaffold_67
Chr. scaffold_68
Chr. scaffold_70
Chr. scaffold_14
Chr. scaffold_39
Chr. scaffold_19
Chr. scaffold_42
Chr. scaffold_2
Chr. scaffold_7
Chr. scaffold_6
Chr. scaffold_69
Chr. scaffold_46
Chr. scaffold_45
Chr. scaffold_20
Coffea canephora (v1.0) Target Chromosome: 11
Query Chr.
Chr. scaffold_51
Chr. scaffold_59
Chr. scaffold_58
Chr. scaffold_15
Chr. scaffold_37
Chr. scaffold_38
Chr. scaffold_34
Chr. scaffold_3
Chr. scaffold_44
Chr. scaffold_42
Chr. scaffold_43
Chr. scaffold_40
Chr. scaffold_48
Chr. scaffold_49

**Figure S2. Self-synteny and fractionation-bias analysis of the *Psychotria viridis* genome (one monoploid) using CoGe, related to Figure 1. (A)** self-versus-self SynMap dot plot showing intragenomic syntenic gene pairs, with the corresponding distribution of synonymous substitution rates ($K_s$) for syntenic gene pairs. **(B)** FractBias analysis showing percent gene retention across sliding windows of genes along each chromosome, relative to each syntenic query chromosome (colored lines).

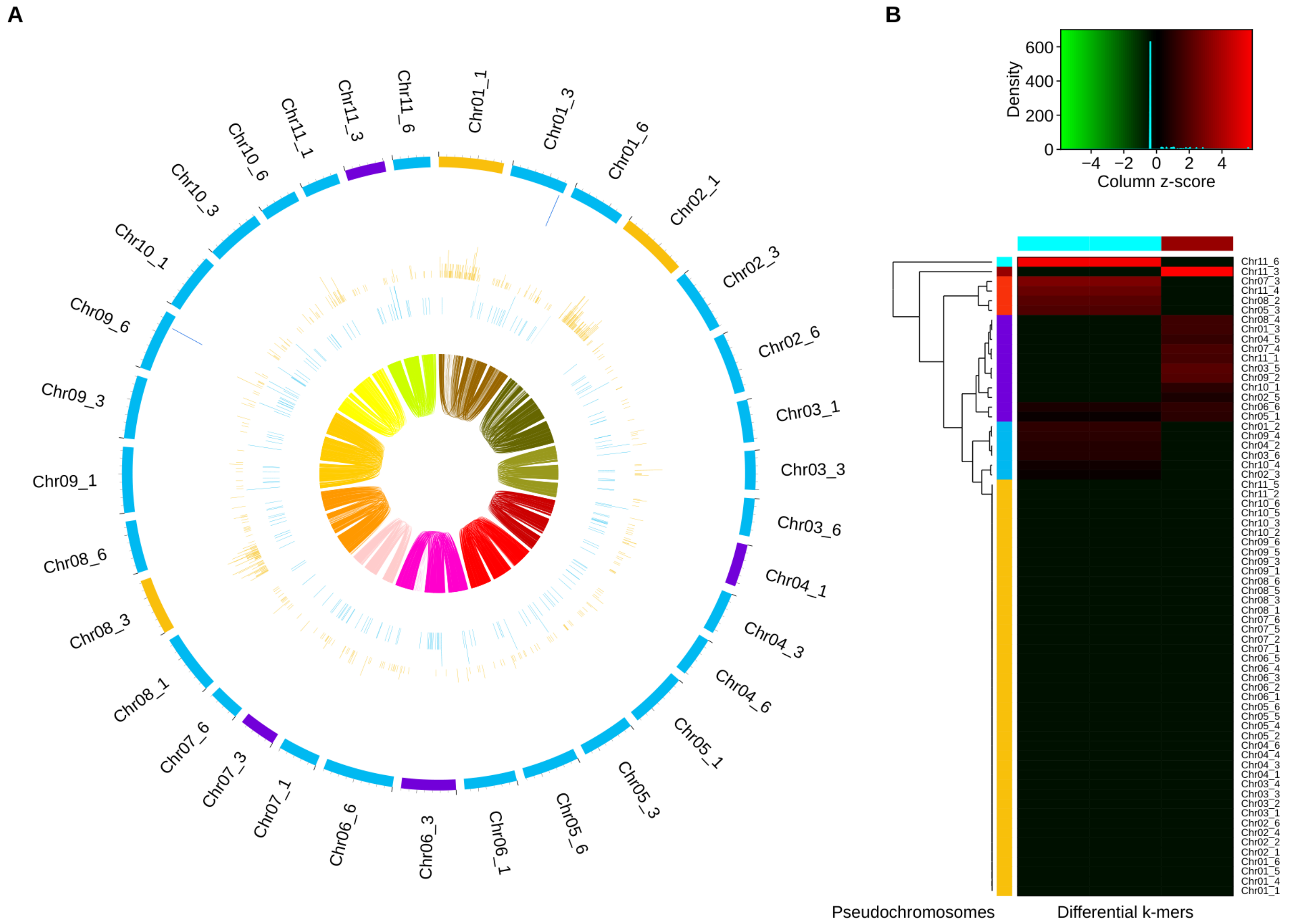


**Figure S3. *K*-mer-based clustering of the pseudochromosomes in *Psychotria viridis* genome using SubPhaser, related to Figure 1. (A)** Circos plot of the SubPhaser results. Only one half of the genome (33 of 66 pseudochromosomes) is included. From outer to inner circles (1–6): (1) pseudochromosomes colored by *k*-means clustering based on differential *k*-mers , with no grouping corresponding to complete monoploid sets; (2) significant enrichment of cluster-specific k-mers; (3–4) absolute count of each cluster-specific *k*-mer set; (5) density of long terminal repeat retrotransposons (LTR-RTs); (6) homeologous blocks (synteny). **(B)** k-mer matrix for the *Psychotria viridis* genome. Heatmap showing the abundance of differential 19-mers across the 66 pseudochromosomes of *P. viridis* ($k$ = 19, minimum frequency $q$ = 50, fold-enrichment threshold $f$ = 2). Columns represent the differential *k*-mers recovered from 60.9 million total 19-mers, grouped by *k*-means clustering into two sets (top color bar, cyan and dark red). Rows represent pseudochromosomes ordered by hierarchical clustering (dendrogram, left); the left color bar indicates the *k*-means cluster assigned to each pseudochromosome. Color indicates the column Z-score of *k*-mer count (green, depleted; red, enriched; black, absent or near-zero signal).

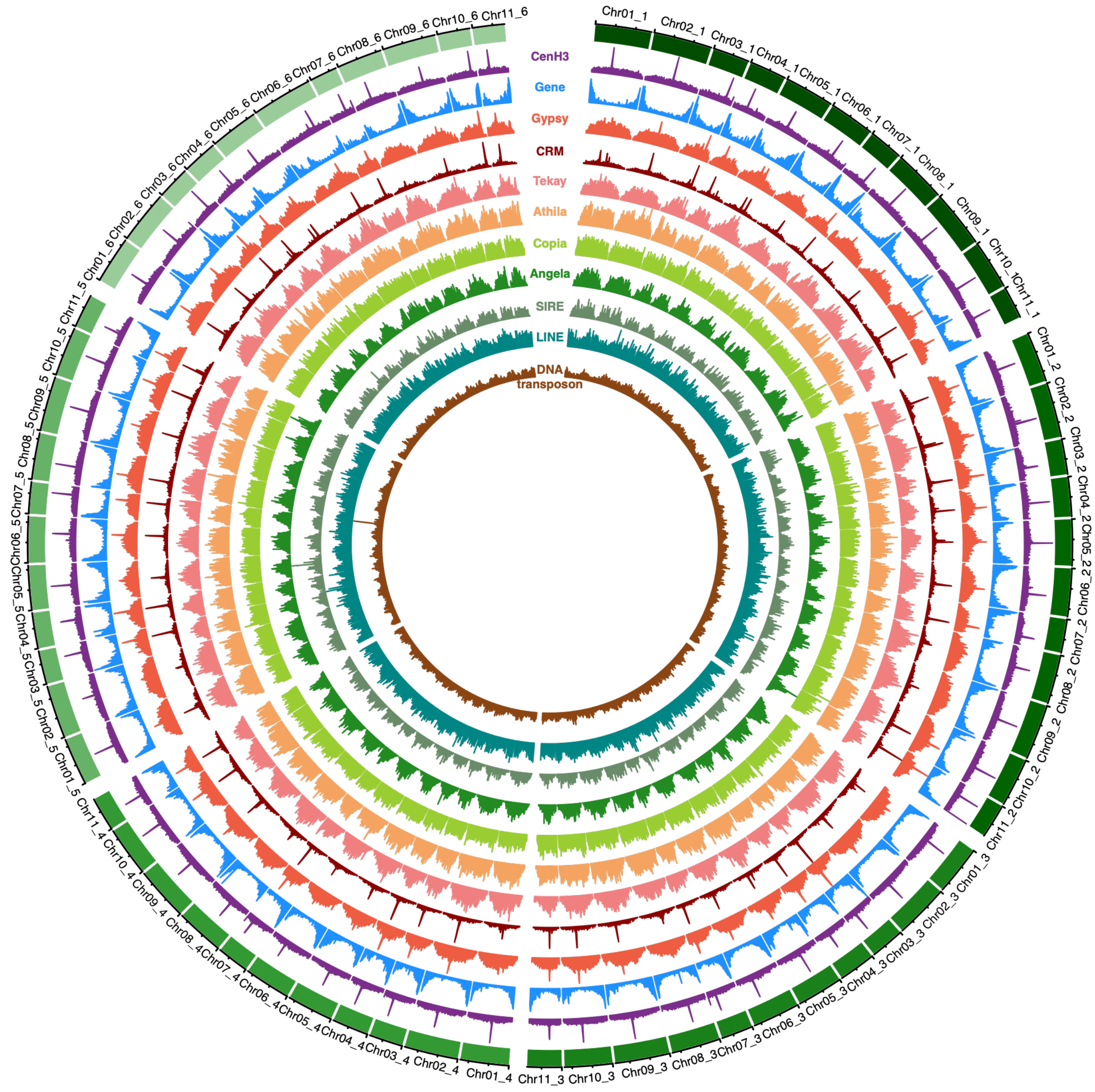


**Figure S4. Genome-wide distribution of genomic features of *Psychotria viridis*, related to Figure 1.** Circos plot showing the genomic distribution of CENH3, genes and repeats across the genome.

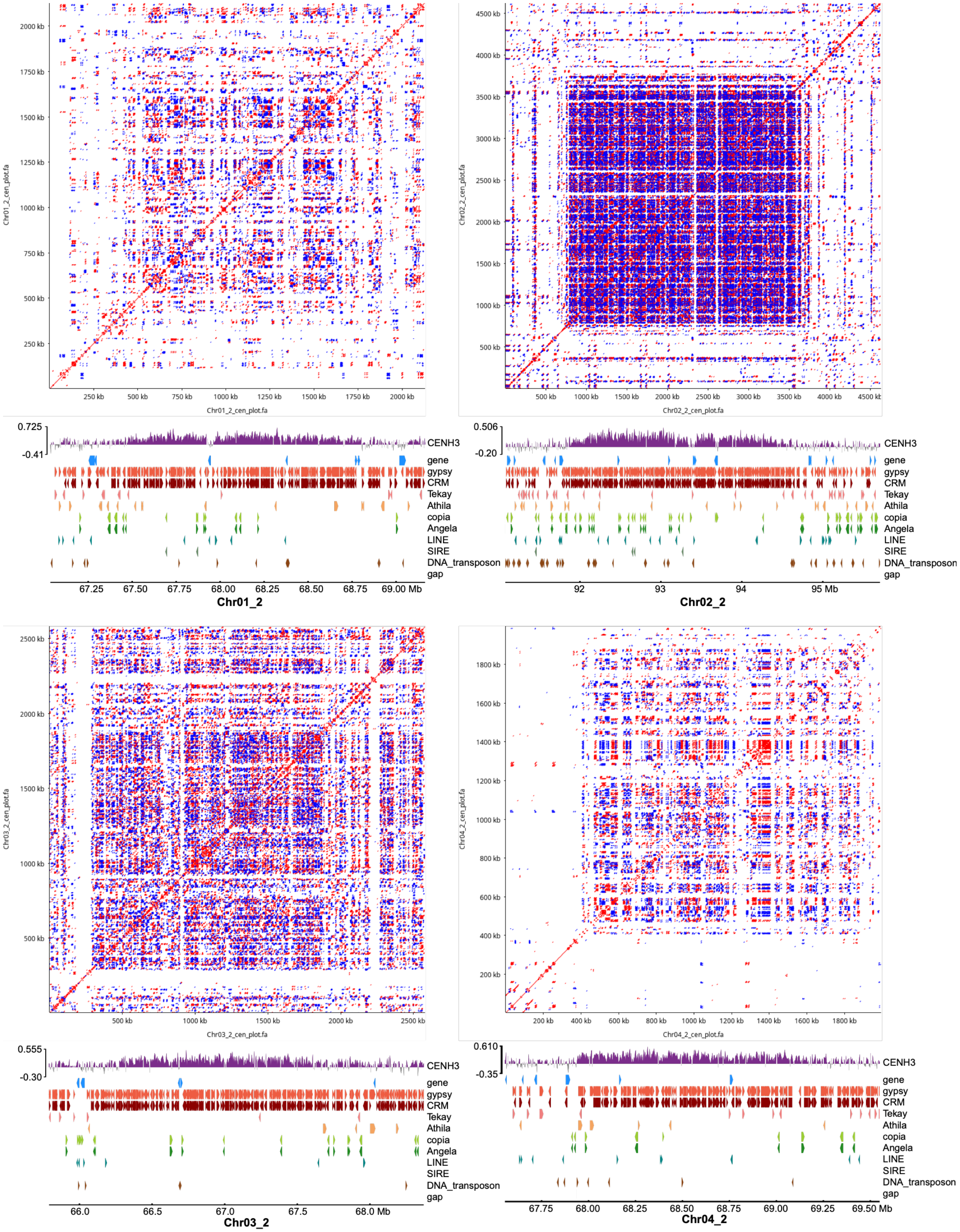

Chr01_2_cen_plot.fa
250 kb
500 kb
750 kb
1000 kb
1250 kb
1500 kb
1750 kb
2000 kb
0.725
-0.41
CENH3
gene
gypsy
CRM
Tekay
Athila
copia
Angela
LINE
SIRE
DNA_transposon
gap
67.25
67.50
67.75
68.00
68.25
68.50
68.75
69.00 Mb
Chr01_2
Chr02_2_cen_plot.fa
500 kb
1000 kb
1500 kb
2000 kb
2500 kb
3000 kb
3500 kb
4000 kb
4500 kb
0.506
-0.20
92
93
94
95 Mb
Chr02_2
Chr03_2_cen_plot.fa
0.555
-0.30
66.0
66.5
67.0
67.5
68.0 Mb
Chr03_2
Chr04_2_cen_plot.fa
200 kb
400 kb
600 kb
800 kb
1000 kb
1200 kb
1400 kb
1600 kb
1800 kb
0.610
-0.35
67.75
68.00
68.25
68.50
68.75
69.00
69.25
69.50 Mb
Chr04_2

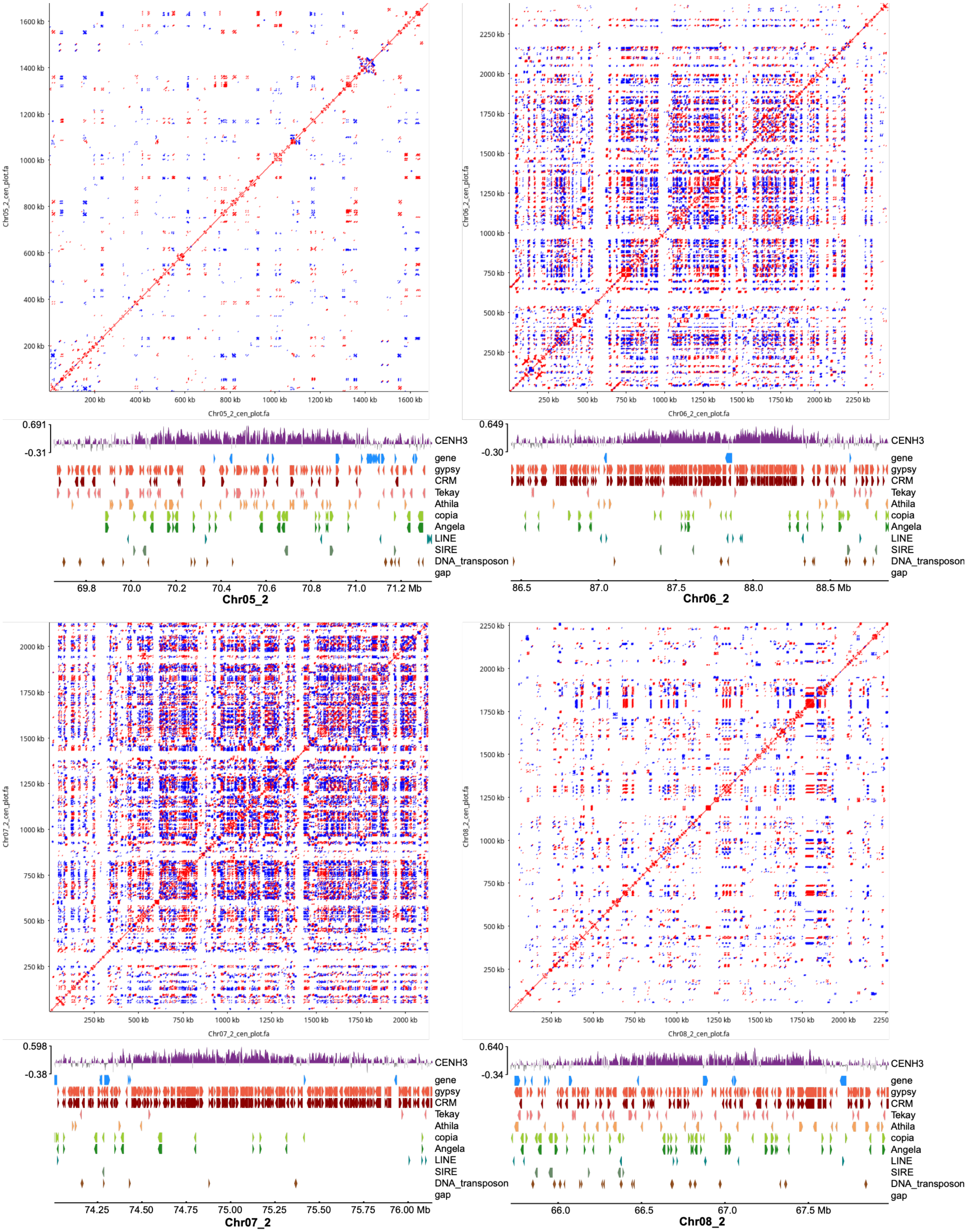

Chr05_2_cen_plot.fa
1600 kb
1400 kb
1200 kb
1000 kb
800 kb
600 kb
400 kb
200 kb
0.691
-0.31
CENH3
gene
gypsy
CRM
Tekay
Athila
copia
Angela
LINE
SIRE
DNA_transposon
gap
69.8
70.0
70.2
70.4
70.6
70.8
71.0
71.2 Mb
Chr05_2
Chr06_2_cen_plot.fa
2250 kb
2000 kb
1750 kb
1500 kb
1250 kb
1000 kb
750 kb
500 kb
250 kb
0.649
-0.30
86.5
87.0
87.5
88.0
88.5 Mb
Chr06_2
Chr07_2_cen_plot.fa
0.598
-0.38
74.25
74.50
74.75
75.00
75.25
75.50
75.75
76.00 Mb
Chr07_2
Chr08_2_cen_plot.fa
0.640
-0.34
66.0
66.5
67.0
67.5 Mb
Chr08_2

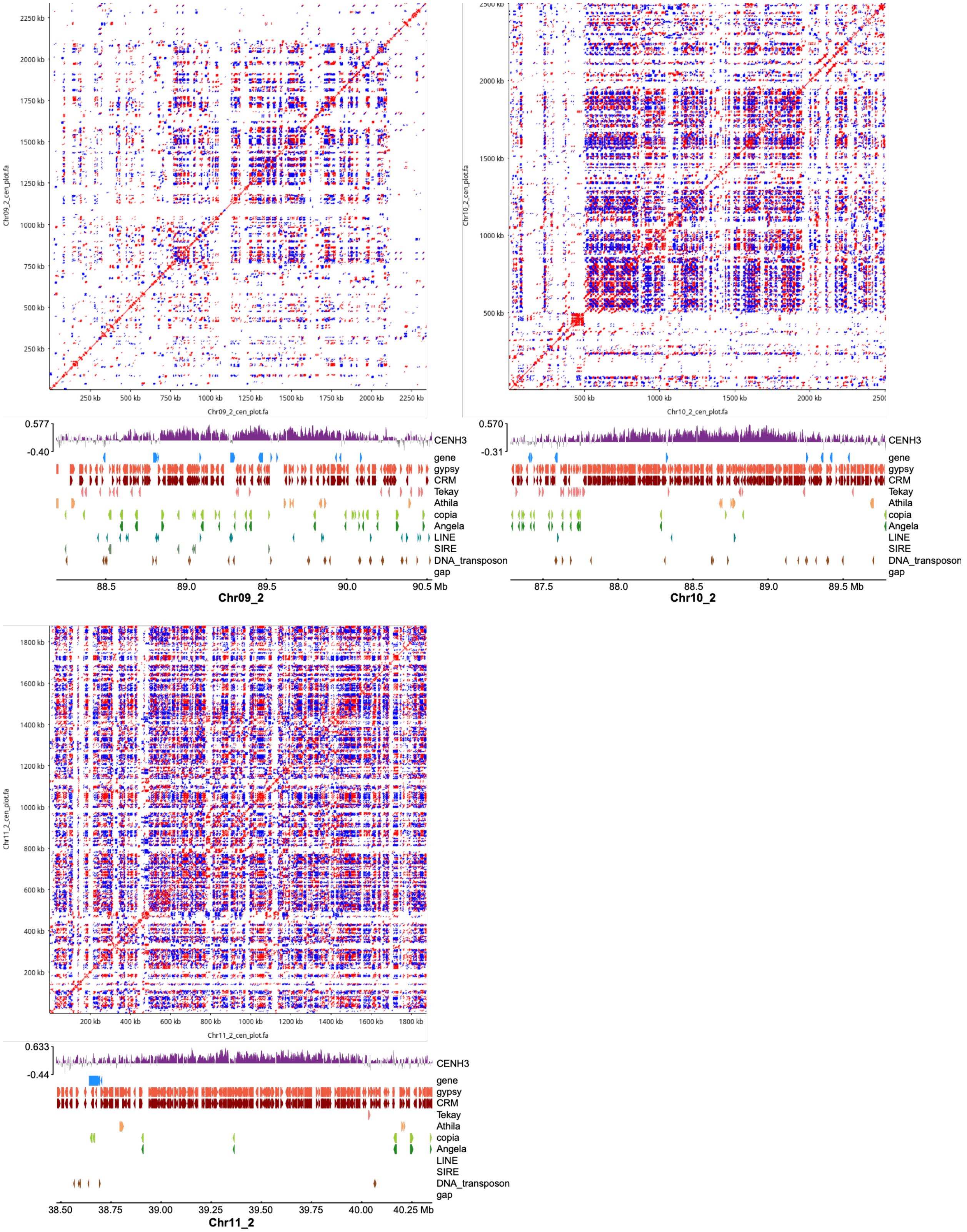


**Figure S5. Self-similarity dotplots and genomic features at centromere of each chromosome in the *Psychotria viridis* genome (monoploid set g2), related to Figure 1.**

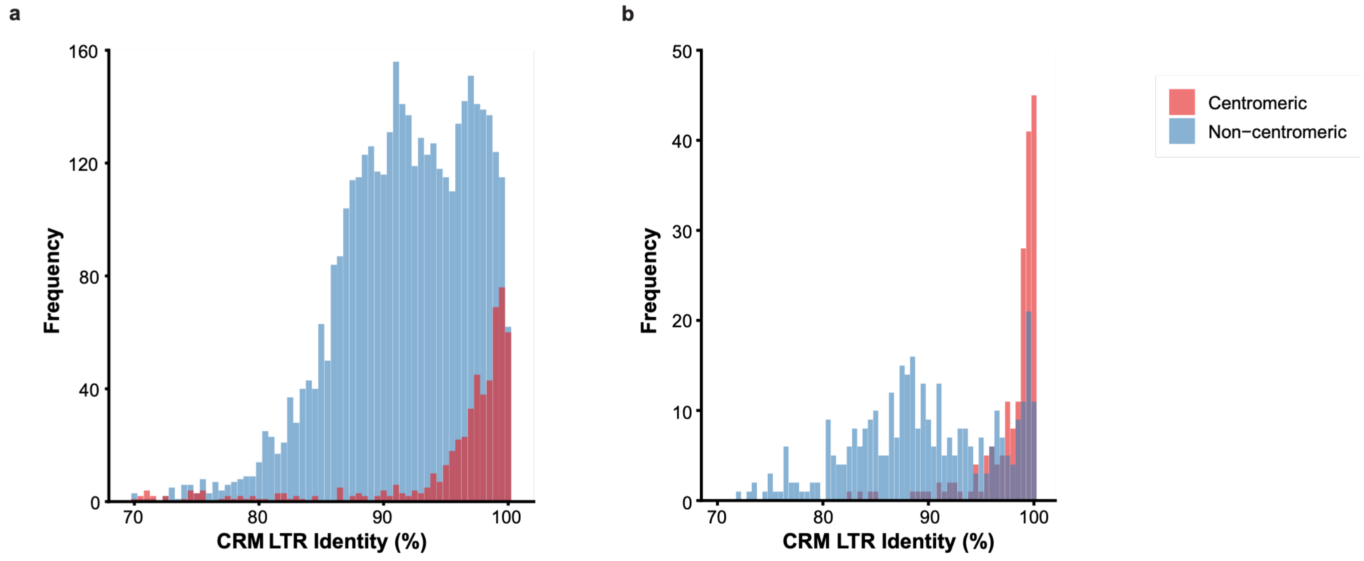


**Figure S6. Distribution of LTR identity (%) for centromeric versus non-centromeric CRMs, related to Figures 1 and 3. (A)** *Psychotria viridis* monoploid set g2. **(B)** *Banisteriopsis caapi* Tucunacá haplotype 1.

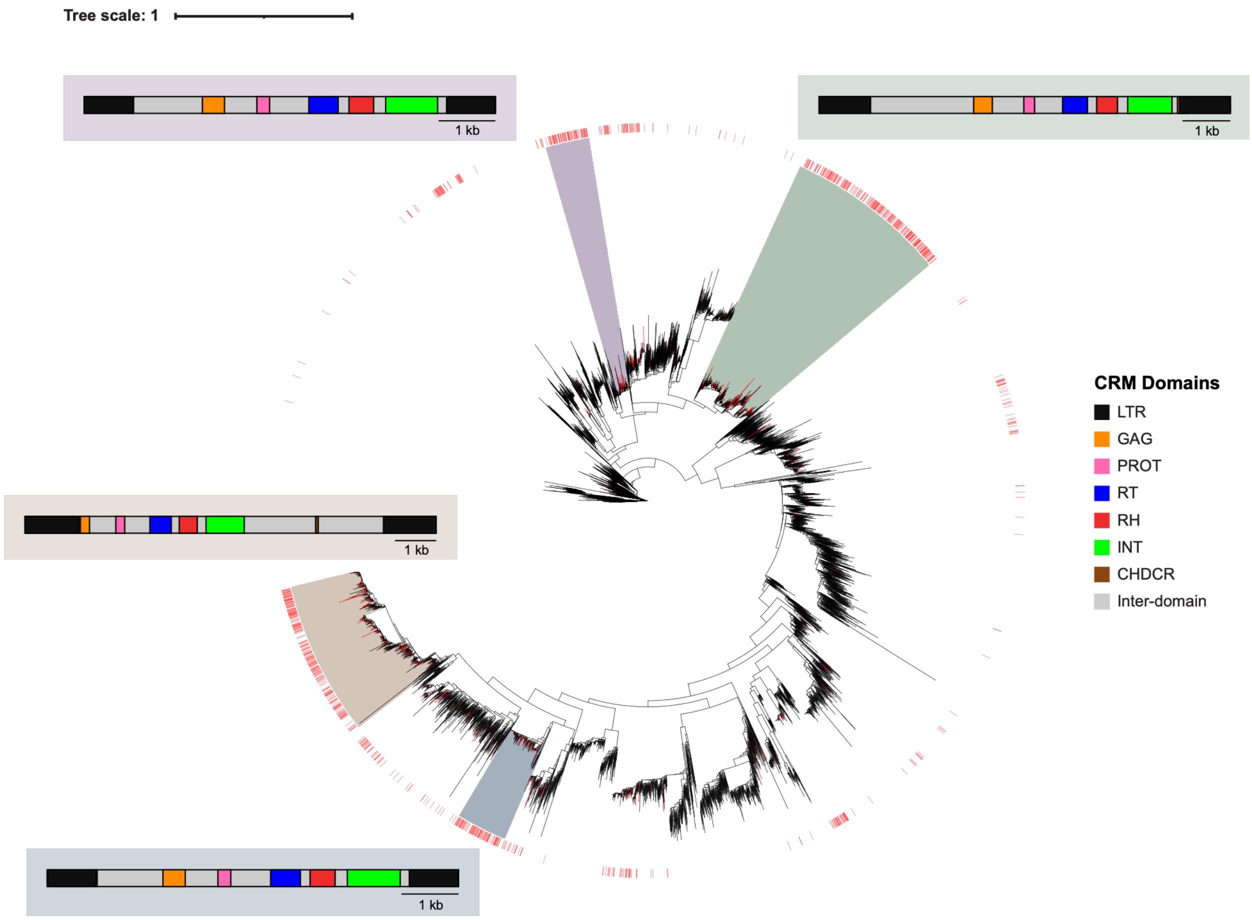


**Figure S7. Phylogenetic tree of CRM elements in the *Psychotria viridis* genome (monoploid set g2), related to Figure 1.** Maximum-likelihood phylogeny based on the RT domain of CRM elements. Red colorstrips mark CRM copies located within centromeres. Color ranges mark phylogenetic clades significantly enriched for centromeric CRMs (one-sided Fisher's exact test, BH-adjusted $q < 0.05$, enrichment ratio > 0.35). Boxed diagrams show the domain structure of a representative, full-length CRM element from each enriched cluster.

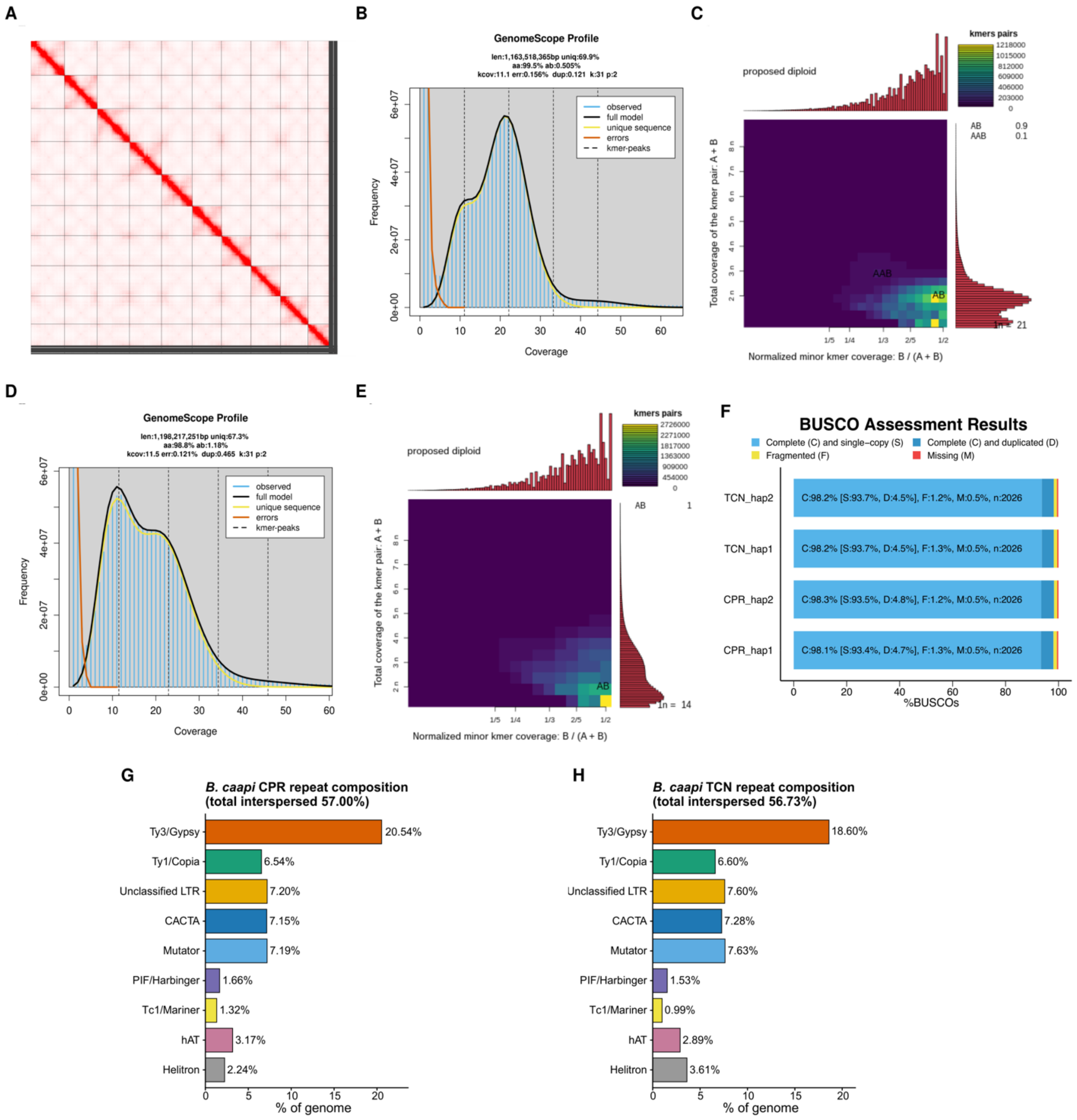


**Figure S8. Genome assembly assessment of *Banisteriopsis caapi* for the varieties Caupuri (CPR) and Tucunacá (TCN), related to Figure 2 and Table S1.** **(A)** Hi-C map of *B. caapi* CPR haplotype 1. **(B)** Genome size and heterozygosity estimation for *B. caapi* CPR based on *31*-mer distribution using GenomeScope2. **(C)** Ploidy and genome structure inference for *B. caapi* CPR based on *31*-mer analysis using Smudgeplot. **(D)** Genome size and heterozygosity estimation for *B. caapi* TCN based on *31*-mer distribution using GenomeScope2. **(E)** Ploidy and genome structure inference for *B. caapi* TCN based on *31*-mer analysis using Smudgeplot. **(F)** BUSCO assessment results (embryophyta_odb12) of each haplotype. **(G)** Interspersed repeat composition of CPR genome. **(H)** Interspersed repeat composition of TCN genome.

a

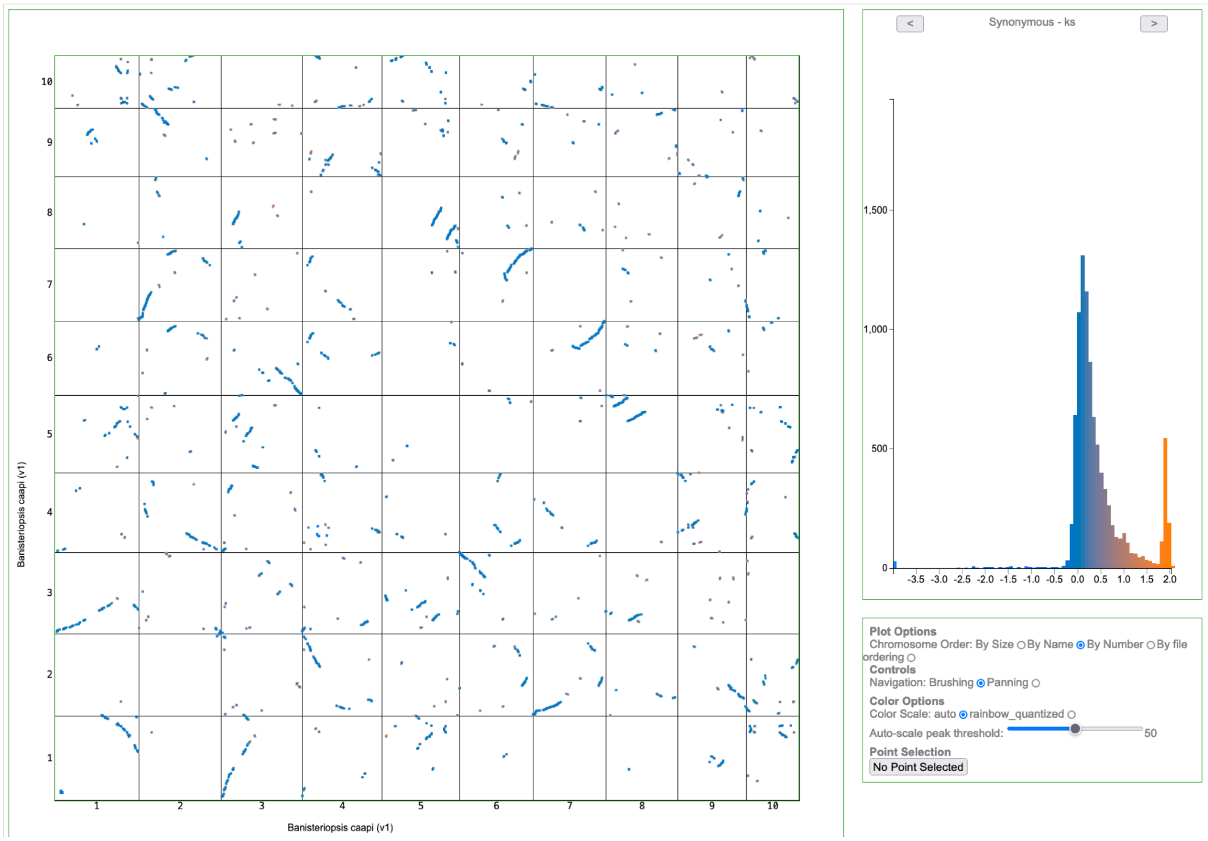

Banisteriopsis caapi (v1)
Banisteriopsis caapi (v1)
Synonymous - ks
Plot Options
Chromosome Order: By Size By Name By Number By file ordering
Controls
Navigation: Brushing Panning
Color Options
Color Scale: auto rainbow_quantized
Auto-scale peak threshold: 50
Point Selection
No Point Selected

b

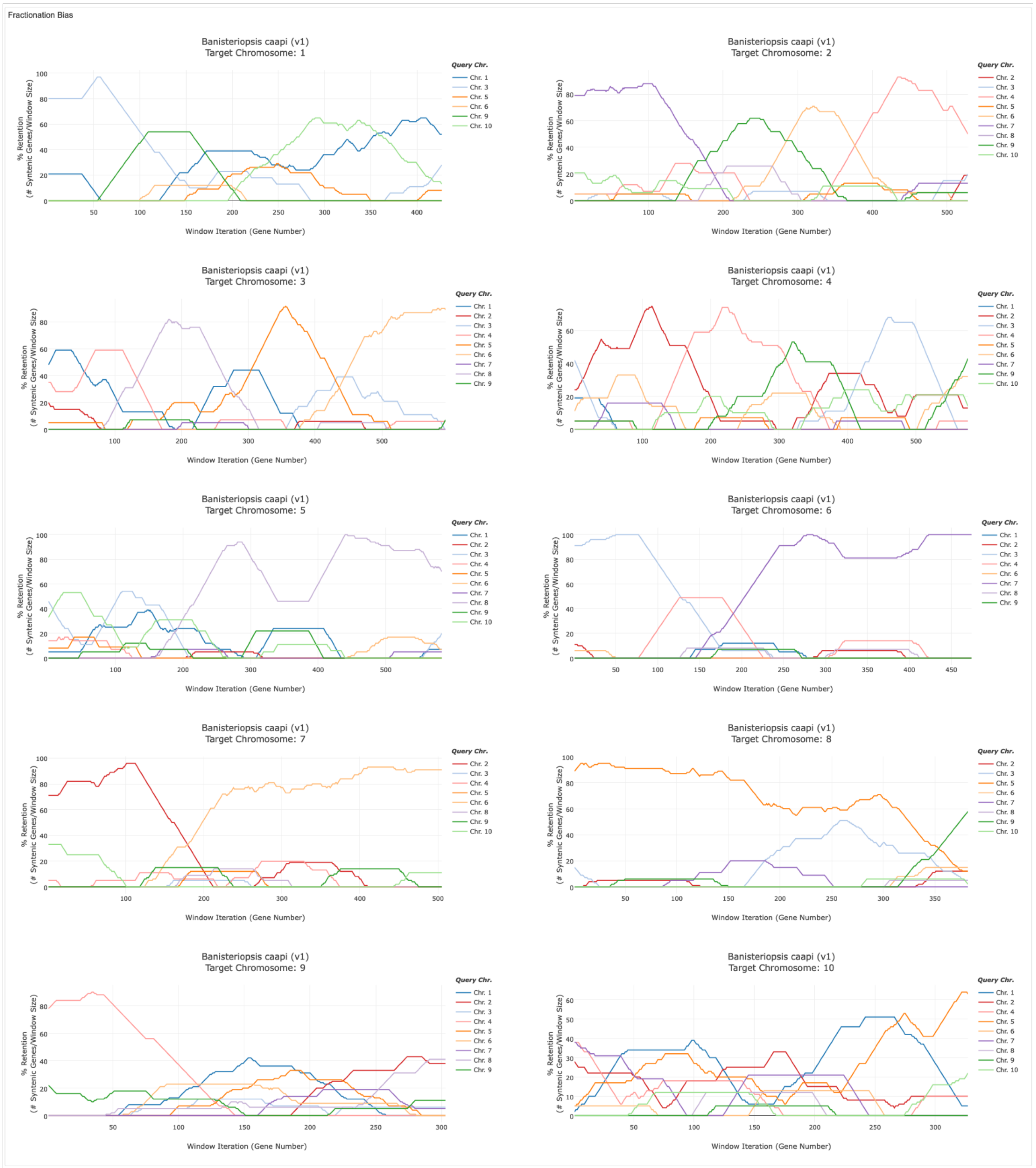


**Figure S9. Self-synteny and fractionation-bias analysis of the *Banisteriopsis caapi* genome (Caupuri haplotype 1) using CoGe, related to Figure 2. (A)** self-versus-self SynMap dot plot showing intragenomic syntenic gene pairs, with the corresponding distribution of synonymous substitution rates ($K_s$) for syntenic gene pairs. **(B)** FractBias analysis showing percent gene retention across sliding windows of genes along each chromosome, relative to each syntenic query chromosome (colored lines).

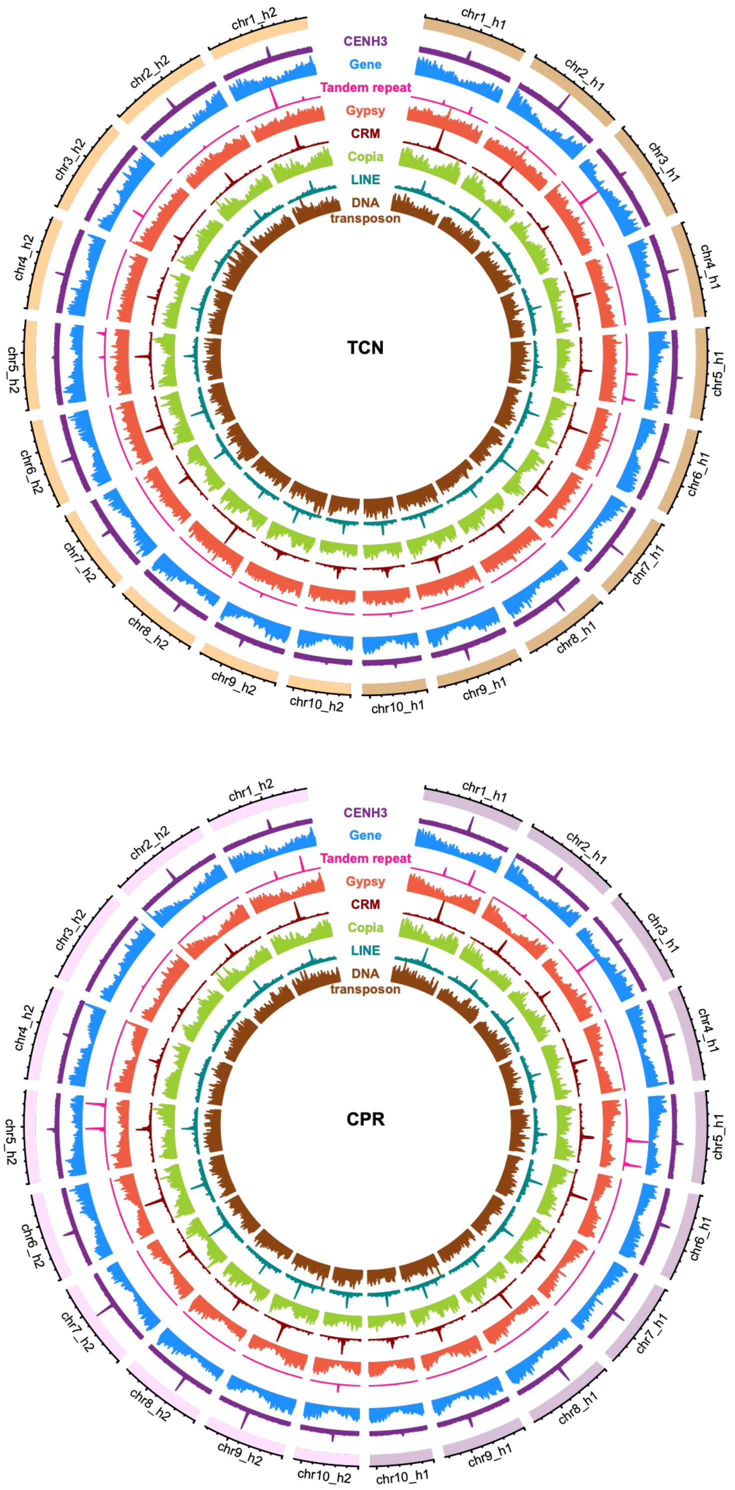


**Figure S10. Genome-wide distribution of genomic features of *Banisteriopsis caapi* Tucunacá (TCN) and Caupuri (CPR), related to Figures 2 and 3.** Circos plots of both haplotypes of *B. caapi* TCN and CPR showing, from outside to inside, the distribution of CENH3 ChIP-seq signal, genes, tandem repeats, Ty3/Gypsy, CRM, Ty1/Copia, LINE and DNA transposons along the chromosomes.

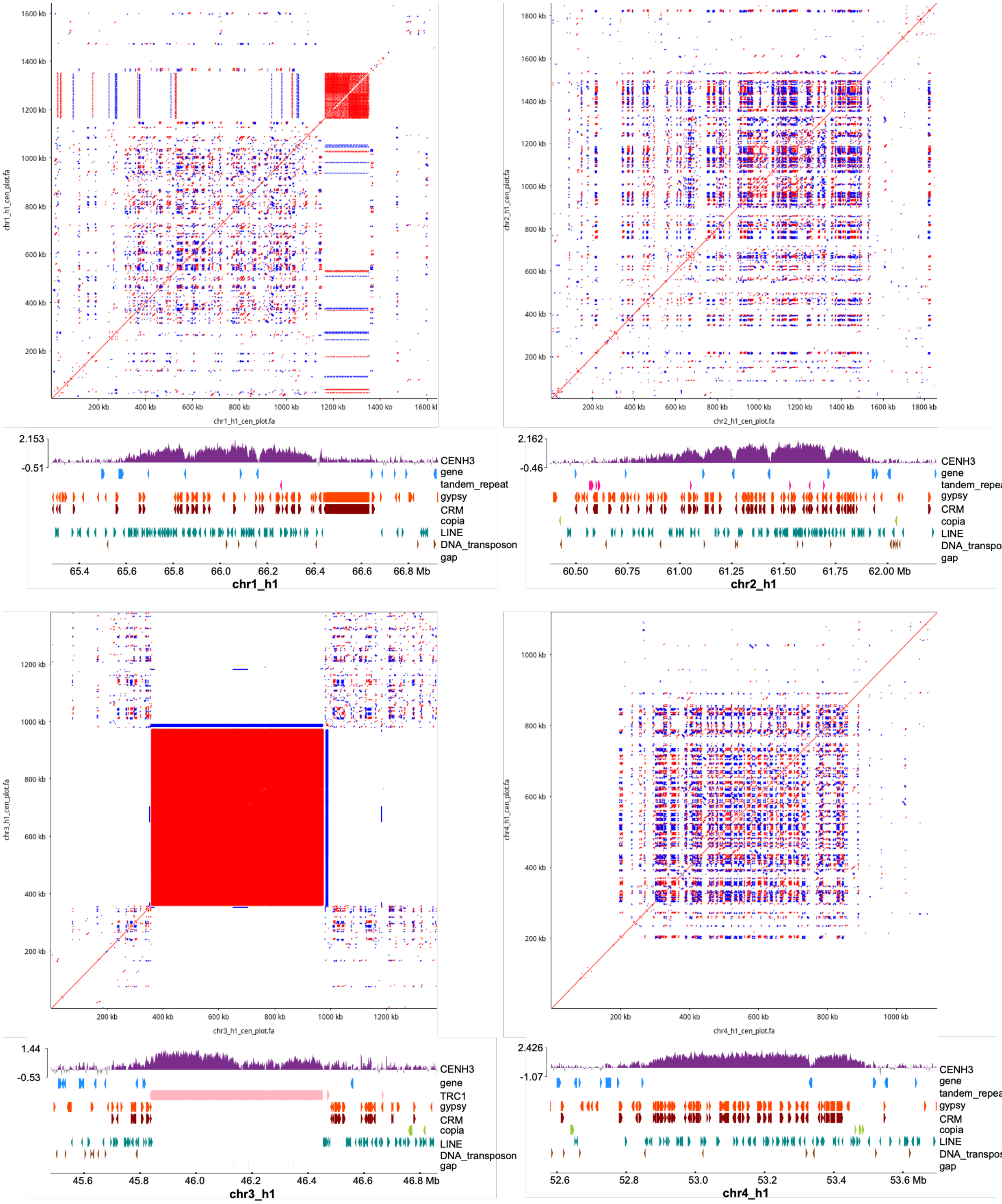

chr1_h1_cen_plot.fa
chr1_h1_cen_plot.fa
200 kb
400 kb
600 kb
800 kb
1000 kb
1200 kb
1400 kb
1600 kb
2.153
-0.51
CENH3
gene
tandem_repeat
gypsy
CRM
copia
LINE
DNA_transposon
gap
65.4
65.6
65.8
66.0
66.2
66.4
66.6
66.8 Mb
chr1_h1
chr2_h1_cen_plot.fa
chr2_h1_cen_plot.fa
1800 kb
2.162
-0.46
CENH3
gene
tandem_repeat
gypsy
CRM
copia
LINE
DNA_transposon
gap
60.50
60.75
61.00
61.25
61.50
61.75
62.00 Mb
chr2_h1
chr3_h1_cen_plot.fa
chr3_h1_cen_plot.fa
1.44
-0.53
CENH3
gene
TRC1
gypsy
CRM
copia
LINE
DNA_transposon
gap
45.6
45.8
46.0
46.2
46.4
46.6
46.8 Mb
chr3_h1
chr4_h1_cen_plot.fa
chr4_h1_cen_plot.fa
2.426
-1.07
CENH3
gene
tandem_repeat
gypsy
CRM
copia
LINE
DNA_transposon
gap
52.6
52.8
53.0
53.2
53.4
53.6 Mb
chr4_h1

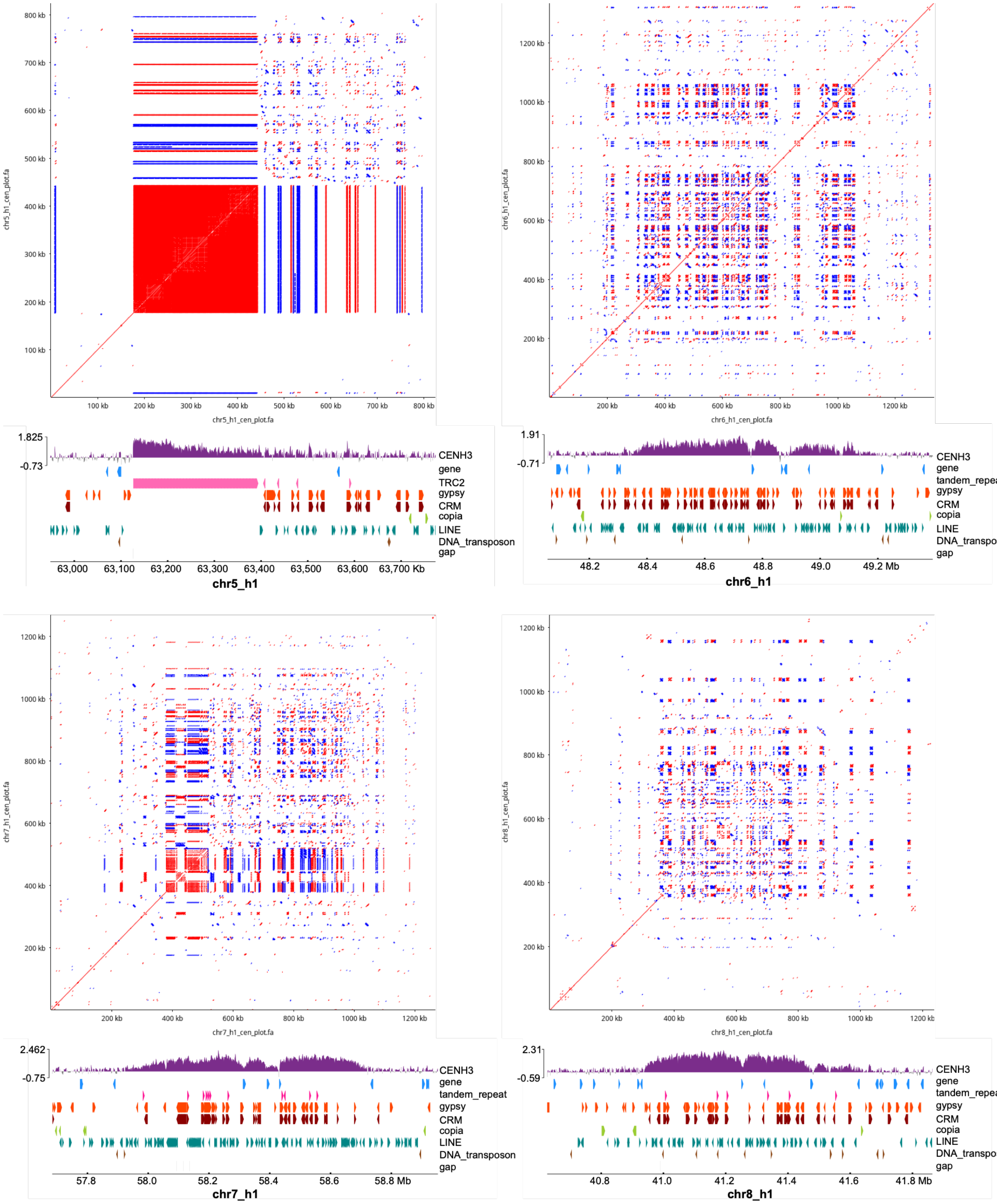

chr5_h1_cen_plot.fa
800 kb
700 kb
600 kb
500 kb
400 kb
300 kb
200 kb
100 kb
CENH3
gene
TRC2
gypsy
CRM
copia
LINE
DNA_transposon
gap
1.825
-0.73
63,000
63,100
63,200
63,300
63,400
63,500
63,600
63,700 Kb
chr5_h1
chr6_h1_cen_plot.fa
1200 kb
1000 kb
800 kb
600 kb
400 kb
200 kb
1.91
-0.71
tandem_repeat
48.2
48.4
48.6
48.8
49.0
49.2 Mb
chr6_h1
chr7_h1_cen_plot.fa
2.462
-0.75
57.8
58.0
58.2
58.4
58.6
58.8 Mb
chr7_h1
chr8_h1_cen_plot.fa
2.31
-0.59
40.8
41.0
41.2
41.4
41.6
41.8 Mb
chr8_h1

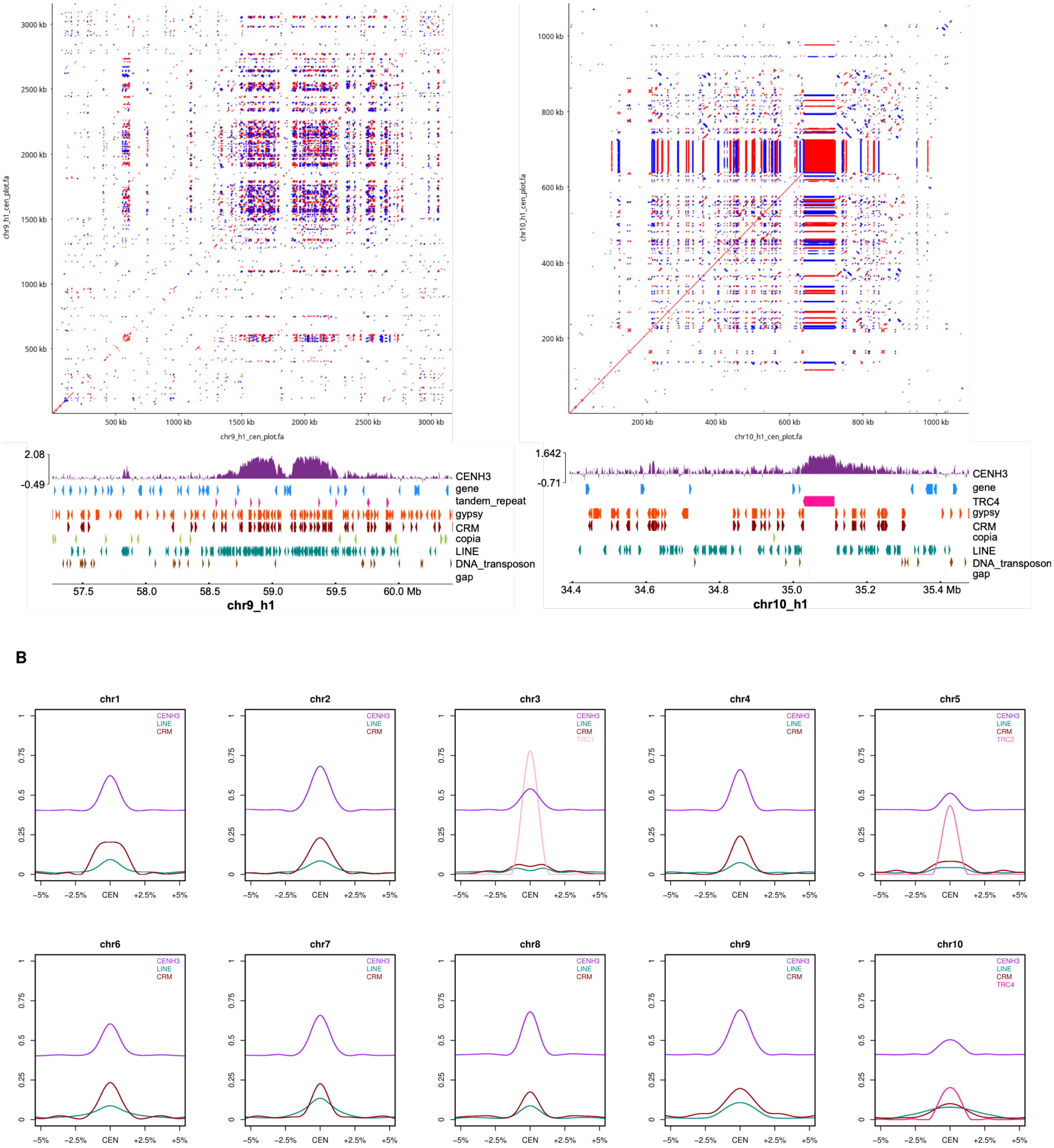


**Figure S11. Centromere organization and CENH3 enrichment in the *Banisteriopsis caapi* genome (Tucunacá haplotype 1), related to Figure 3. (A)** Self-similarity dotplots and genomic features at the centromere of each chromosome. **(B)** Metaplots of CENH3/H3 enrichment and of LINE, CRM and TRC coverage across the centromere of each chromosome, from 5% of chromosome length on either side of the centromere (CEN), in 1-kb windows. See also Figure 3E and Tables S7 and S8.

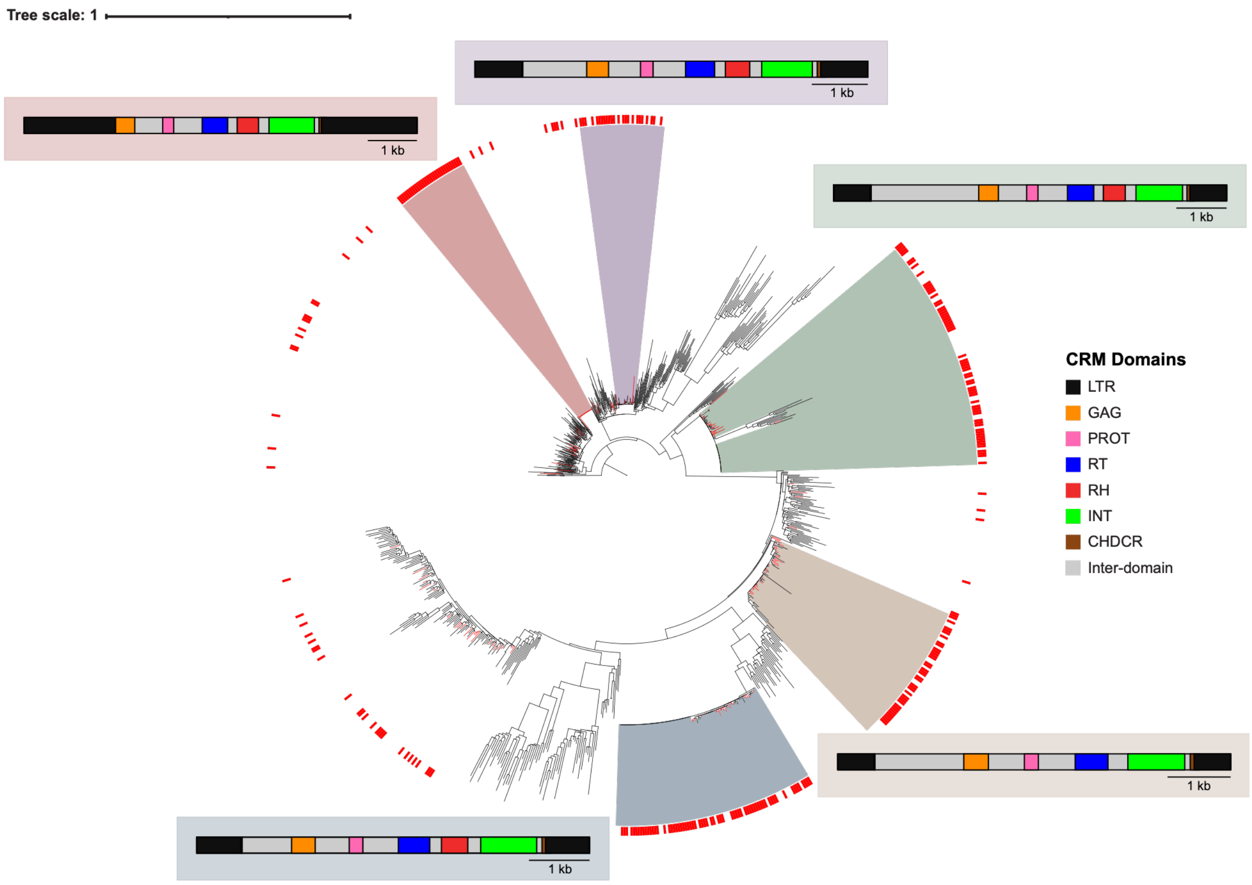


**Figure S12. Phylogenetic tree of CRM elements in the *Banisteriopsis caapi* genome (Tucunacá haplotype 1), related to Figure 3.** Maximum-likelihood phylogeny based on the RT domain of CRM elements. Red colorstrips mark CRM copies located within centromeres. Color ranges mark phylogenetic clades significantly enriched for centromeric CRMs (one-sided Fisher's exact test, BH-adjusted $q < 0.05$, enrichment ratio $> 0.6$). Boxed diagrams show the domain structure of a representative, full-length CRM element from each enriched cluster.

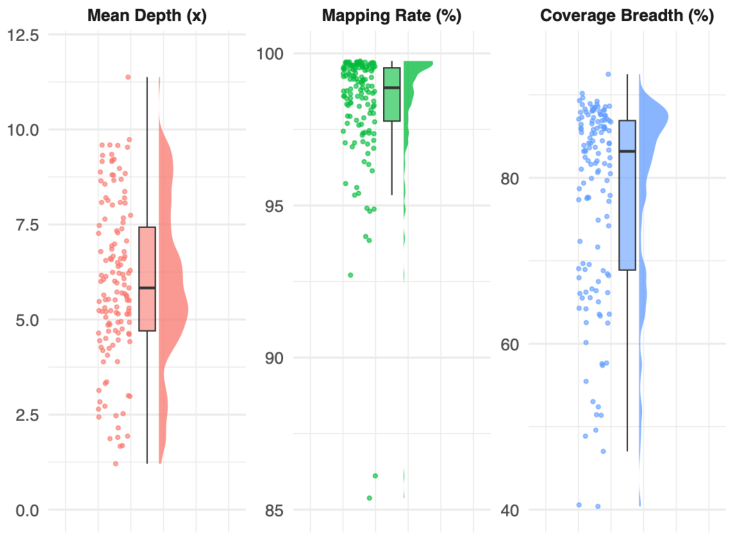


**Figure S13. Population resequencing statistics for *Banisteriopsis caapi*, related to Figure 4.** Reads from 135 individuals were mapped to the *B. caapi* Tucunacá haplotype 1 reference genome.

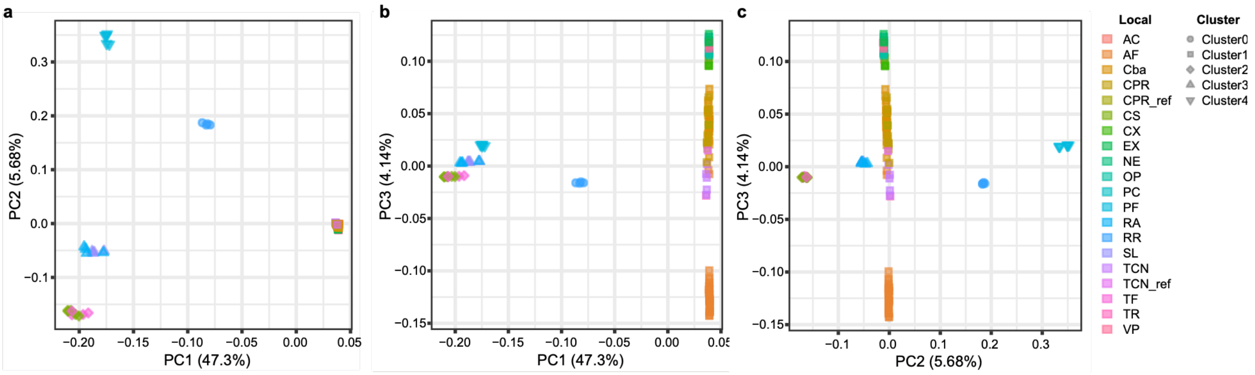


**Figure S14. Two-dimensional projections of PCA of *Banisteriopsis caapi* populations, related to Figure 4.** Pairwise projections of the first three principal components: **(A)** PC1 vs. PC2 (as in Figure 4B); **(B)** PC1 vs. PC3; **(C)** PC2 vs. PC3. Points are colored by sampling locality and shaped by k-means cluster assignment. The CPR and TCN reference genotypes are included in addition to the 135 resequenced accessions (137 samples).

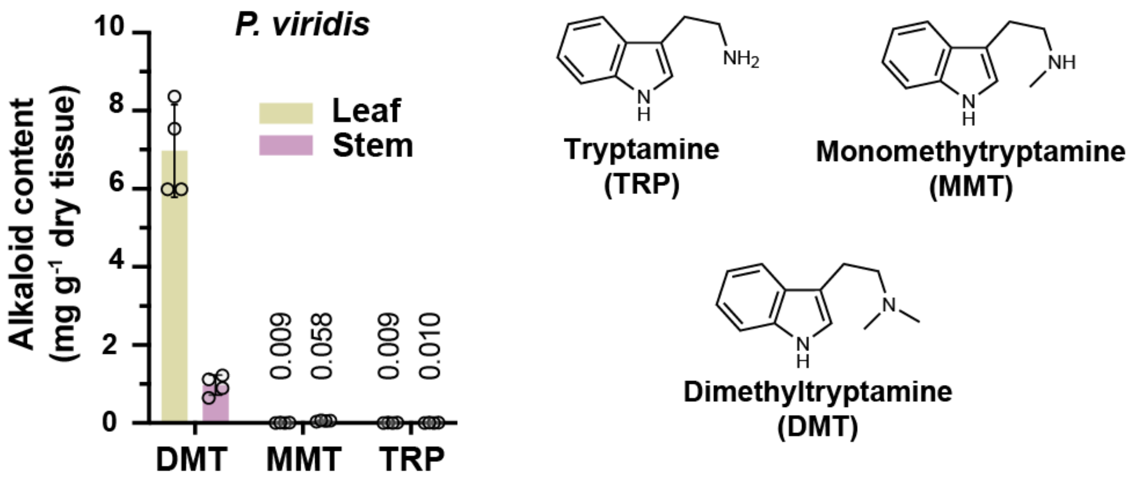


**Figure S15. Quantification of di-methyltryptamine (DMT), mono-methyltryptamine (MMT) and tryptamine (TRP) levels in dry *Psychotria viridis* leaf and stem tissues, related to Figure 5.** Data represent mean values from four biological replicates (circles). Error bars indicate standard deviations.

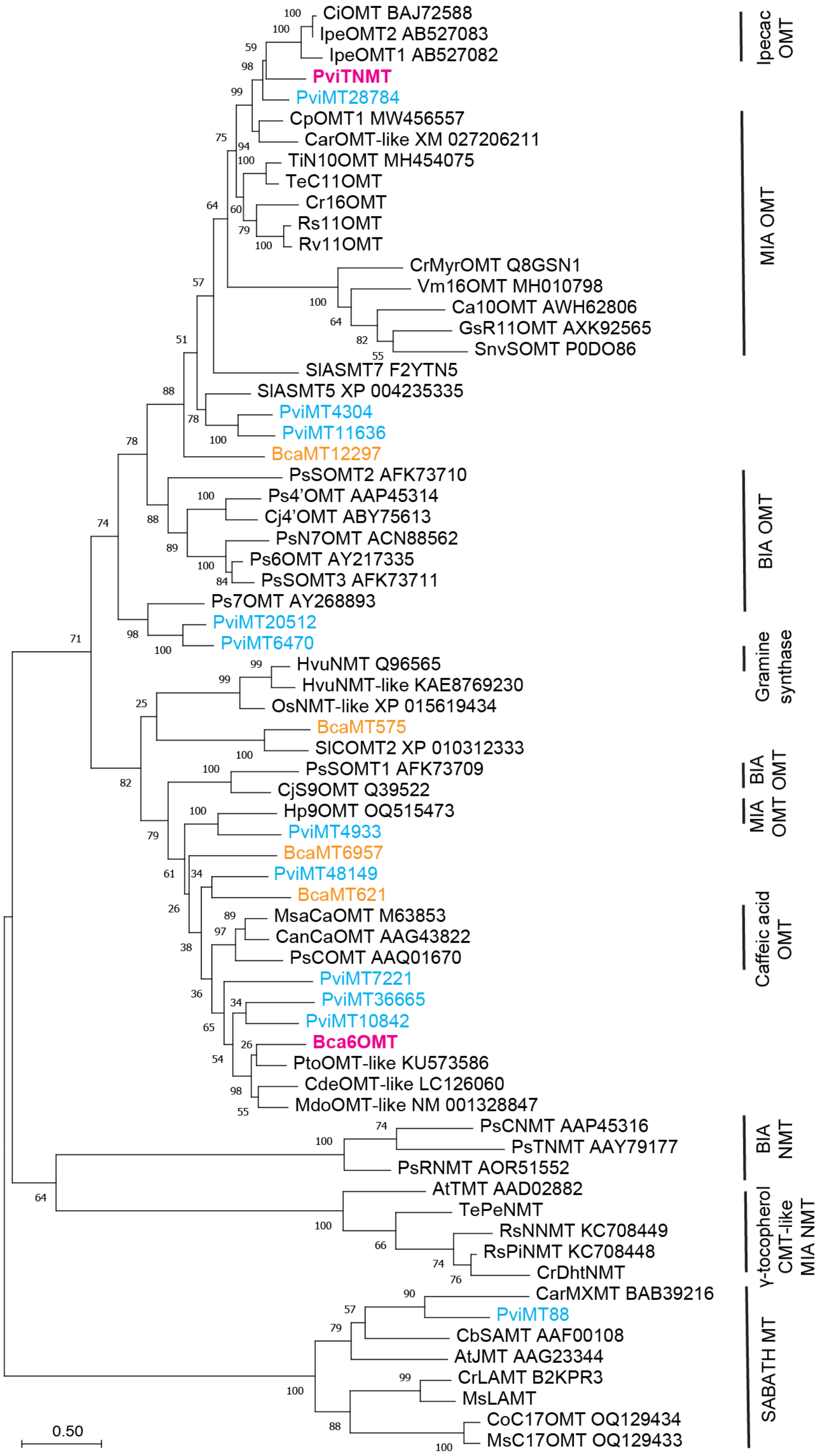


**Figure S16. Phylogenetic analysis of the methyltransferases (MT) candidates' protein sequences from *Psychotria viridis* and *Banisteriopsis caapi*, related to Figures 5 and 6.** The enzymes investigated in this study are color-labeled. The MT types and corresponding metabolite classes catalyzed by them are also indicated, including those involved in monoterpenoid indole alkaloid (MIA), benzylisoquinoline alkaloid (BIA), and ipecac alkaloid biosynthesis. PviTNMT clusters with characterized OMTs involved in MIA and ipecac alkaloid biosynthesis from the Rubiaceae and Apocynaceae families within the order Gentianales. In contrast, it is distantly related to gramine synthase from *Hordeum vulgare* (barley), which sequentially installs two methyl groups on the amine of aminomethylindole, an analog of tryptamine. Bca6OMT groups with numerous uncharacterized methyltransferases as well as several caffeic acid OMTs, although the amino acid sequence identity among these enzymes is low (<55%). Evolutionary analyses were conducted in MEGA11 using protein sequences. The evolutionary history was inferred by using the Maximum Likelihood method and JTT matrix-based model. The percentage of trees in which the associated taxa clustered together is shown next to the branches (100 bootstrap replicates). Initial tree(s) for the heuristic search were obtained automatically by applying Neighbor-Join and BioNJ algorithms to a matrix of pairwise distances estimated using the JTT model, and then selecting the topology with superior log likelihood value. The tree is drawn to scale, with branch lengths measured in the number of substitutions per site (scale bar). At: *Arabidopsis thaliana*; Bca: *Banisteriopsis caapi*; Ca: *Camptotheca acuminata*; Can: *Capsicum annuum*; Car: *Coffea arabica*; Cb: *Clarkia breweri*; Cde: *Citrus dep*ressa; Ci/Ipe: *Carapichea ipecacuanha*; Cj: *Coptis chinensis/japonica*; Co: *Cephalanthus occidentalis*; Cp: *Cinchona pubescens*; Cr: *Catharanthus roseus*; Gs: *Gelsemium sempervirens*; Hp: *Hamelia patens*; Hvu: *Hordeum vulgare*; Mdo: *Malus domestica*; Ms: *Mitragyna speciosa*; Msa: *Medicago sativa*; Os: *Oryza sativa*; Ps: *Papaver somniferum*; Pto: *Populus tomentosa*; Pvi: *Psychotria viridis*; Rs: *Rauvolfia serpentina*; Rv: *Rauvolfia verticillata*; Sl: *Solanum lycopersicum*; Snv: *Strychnos nux-vomica*; Ti: *Tabernanthe iboga*; Te: *Tabernaemontana elegans*; Vm: *Vinca minor*. The plant species, amino acid sequences are included in data S1.

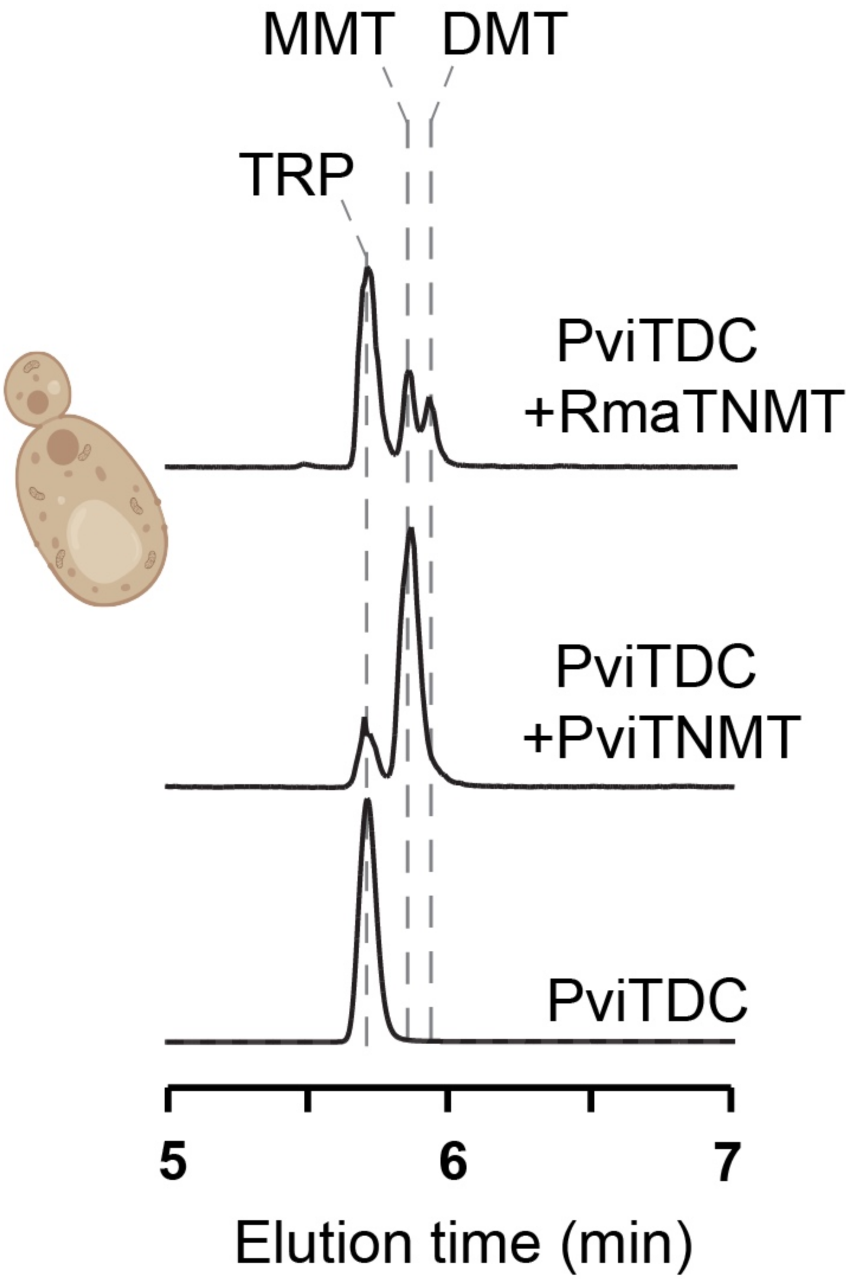


**Figure S17. Expressing *Psychotria viridis* tryptophan decarboxylase (TDC), tryptamine *N*-methyltransferases (TNMT), and a characterized NMT from *Rhinella marina* (cane toad) in *Saccharomyces cerevisiae* led to the de novo biosynthesis of mono-methyltryptamine (MMT) and di-methyltryptamine (DMT) in yeast media, related to Figure 5.** LC-MS/MS analysis shows extracted-ion chromatograms (EICs) corresponding to the following $[M+H]^+$ ions: *m/z* 161 for tryptamine (TRP), 175 for monomethyltryptamine (MMT), and 189 for dimethyltryptamine (DMT).

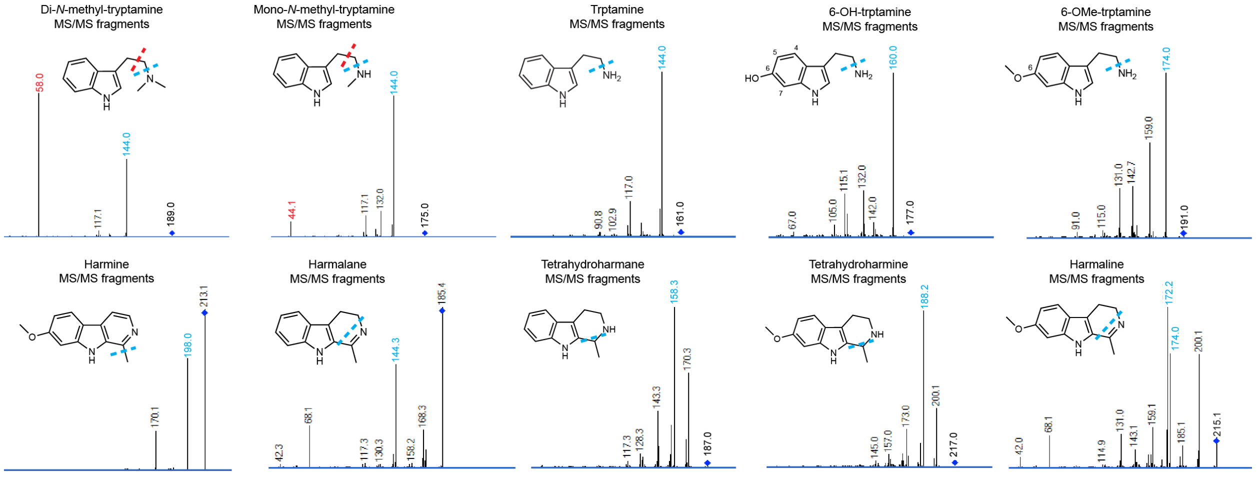


**Figure S18. MS/MS fragmentation patterns for alkaloids in this study, related to Figures 5 and 6.** Colored dashed lines indicate the specific fragment ions generated during MS/MS fragmentation, corresponding to the identically colored *m/z* values shown in the fragmentation spectra.

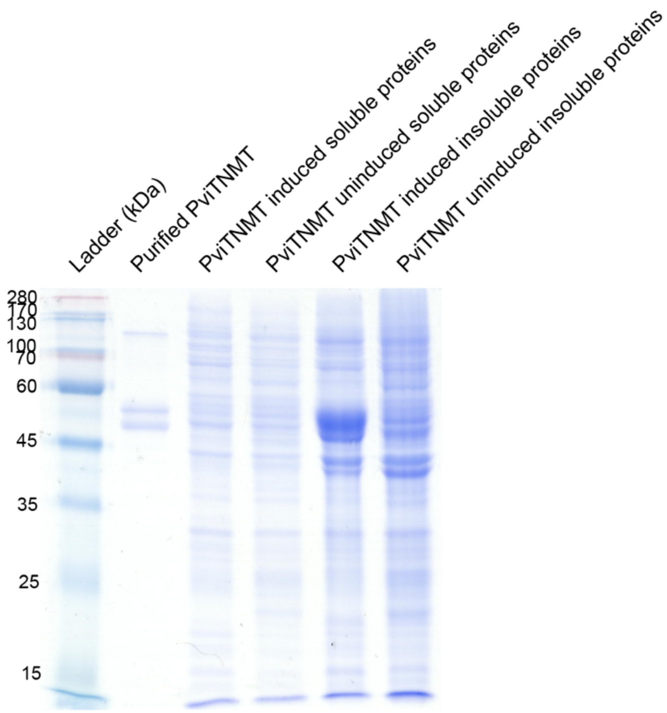


**Figure S19. SDS-PAGE showing the purification of his-tagged *Psychotria viridis* tryptamine *N*-methyltransferases (PviTNMT) from *E. coli* BL21DE3, related to Figure 5.**

**A**

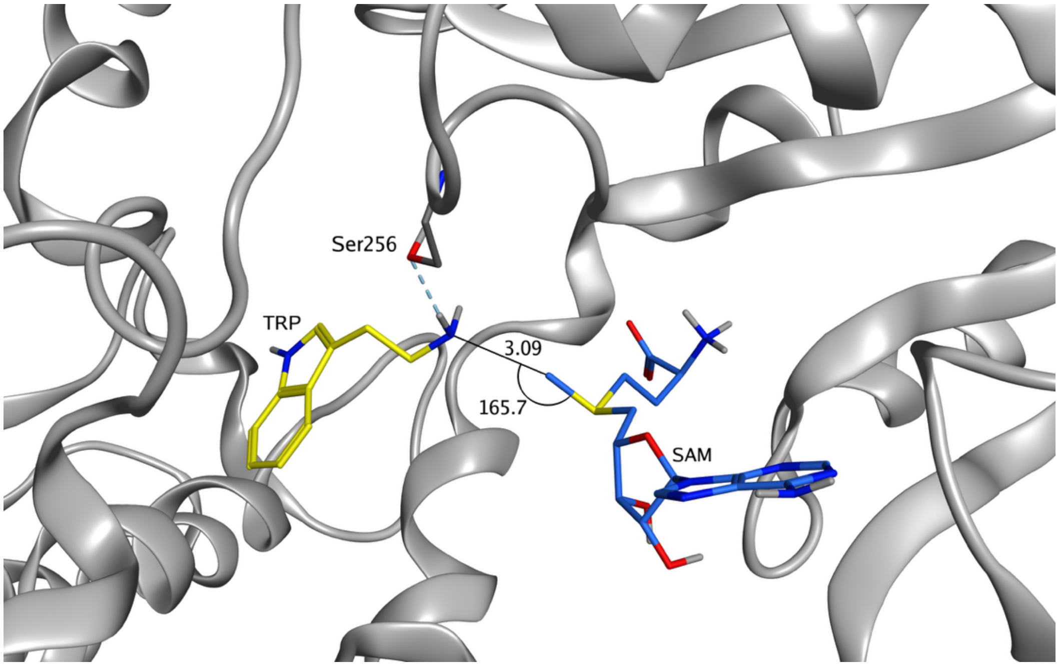


**B**

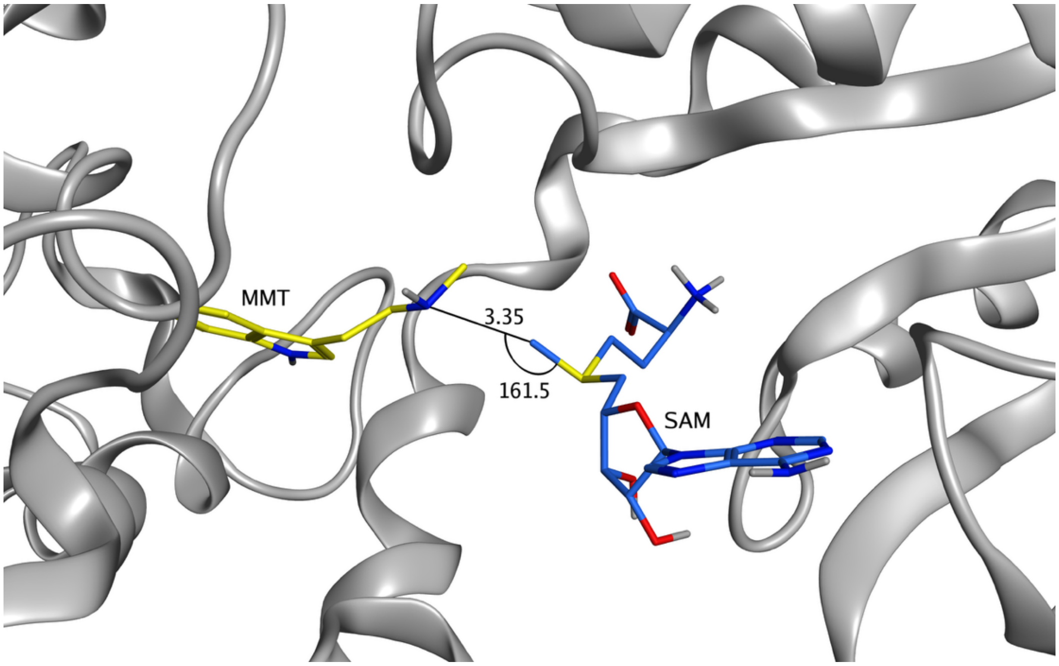

**Figure S20. Homology modeling illustrates substrate binding at the active site of *Psychotria viridis* tryptamine *N*-methyltransferases (PviTNMT), related to Figure 5. (A)** Docking of tryptamine (TRP) into the PviTNMT active site positioned the amine nitrogen optimally toward the sulfonium methyl group of S-adenosylmethionine (SAM), with a distance of 3.09 Å and a dihedral angle of 165.7°, consistent with an $S_N2$ nucleophilic attack from the amine to the sulfonium methyl group. The calculated docking score was −4.8324 kcal/mol. **(B)** Docking of monomethyltryptamine (MMT) revealed a less favorable geometry, with a reduced dihedral angle (161.5°) and an increased distance between the amine nitrogen and the sulfonium methyl group of SAM, resulting in a weaker docking score (−4.5486 kcal/mol). These structural differences are consistent with the experimentally determined kinetic parameters of PviTNMT, which show a higher catalytic efficiency toward tryptamine ($K_M$ = 4.56 µM; $k_{cat}$ = 28.8 × $10^{-3}$ $s^{-1}$) compared to monomethyltryptamine ($K_M$ = 100.4 µM; $k_{cat}$ = 0.84 × $10^{-3}$ $s^{-1}$). Homology modeling and molecular docking were performed using the Molecular Operating Environment (MOE) 2022.02. Docking poses were manually inspected to identify configurations conducive to the relevant catalytic reactions, and the optimal pose defined by the lowest docking energy and a biochemically plausible orientation was selected.

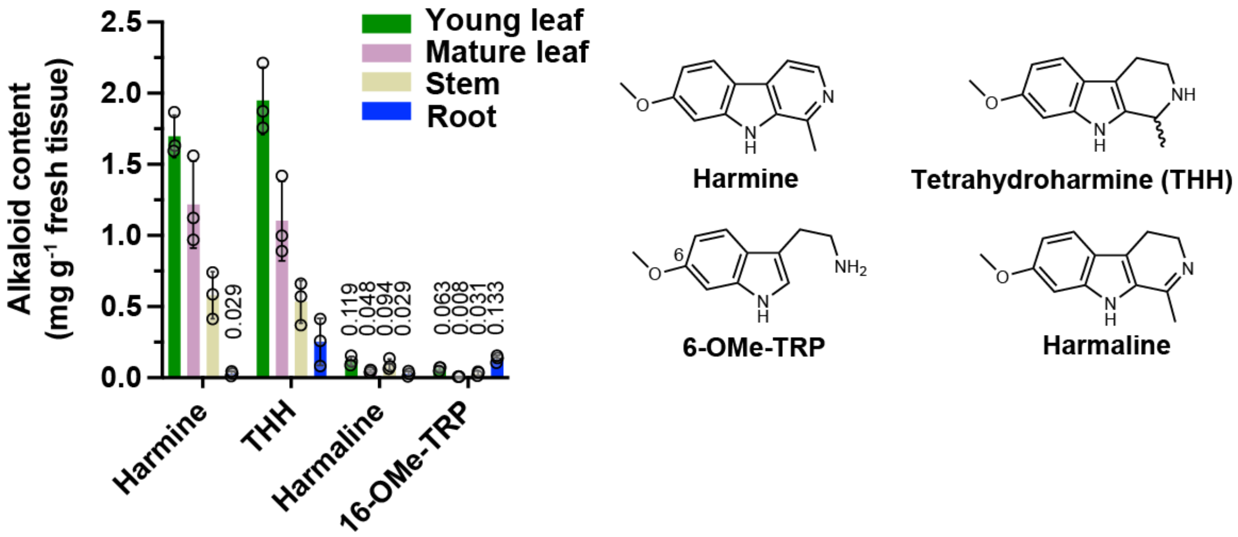


**Figure S21. Quantification of the four most abundant alkaloids harmine, tetrahydroharmine (THH), harmaline, and 6-methoxy-tryptamine levels in fresh *Banisteriopsis caapi* young leaf, old leaf, stem and root tissues, related to Figures 6 and 7.** Data represent mean values from three biological replicates (circles). Error bars indicate standard deviations.

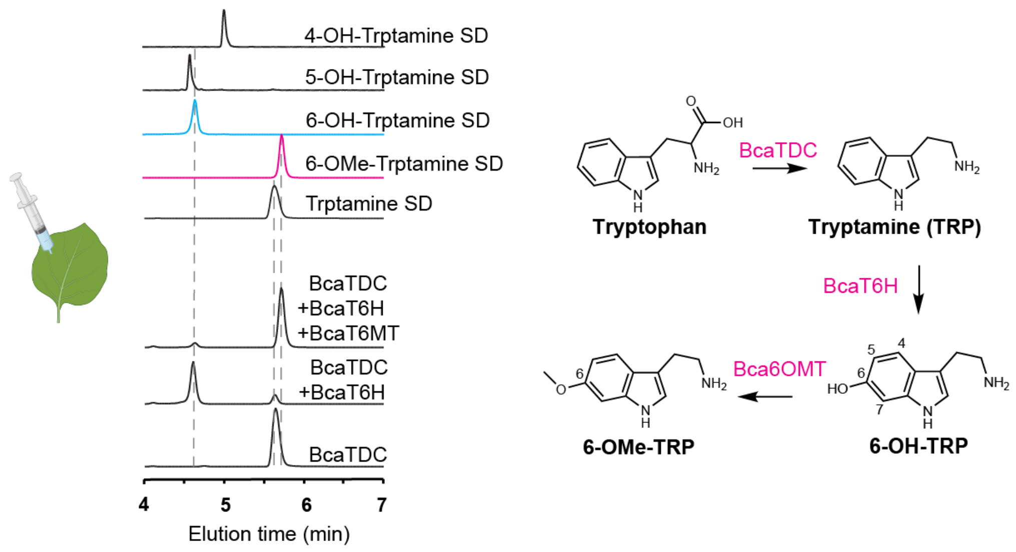


**Figure S22. Transient expression of *Banisteriopsis caapi* tryptophan decarboxylase (BcaTDC), tryptamine 6-hydroxylase (BcaT6H), and 6-hydroxy-tryptamine *O*-methyltransferase (Bca6OMT) results in 6-hydroxy-tryptamine and 6-methoxy-tryptamine de novo biosynthesis in *Nicotiana benthamiana* leaves, related to Figure 6.** Metabolites in *N. benthamiana* leaves were analyzed 3 days after multiplexed *Agrobacterium*-mediated transient gene expression. LC-MS/MS analysis shows extracted-ion chromatograms (EICs) corresponding to the following $[M+H]^+$ ions: *m/z* 161 for tryptamine, *m/z* 177 for 6-hydroxy-tryptamine, and *m/z* 191 for 6-methoxy-tryptamine.

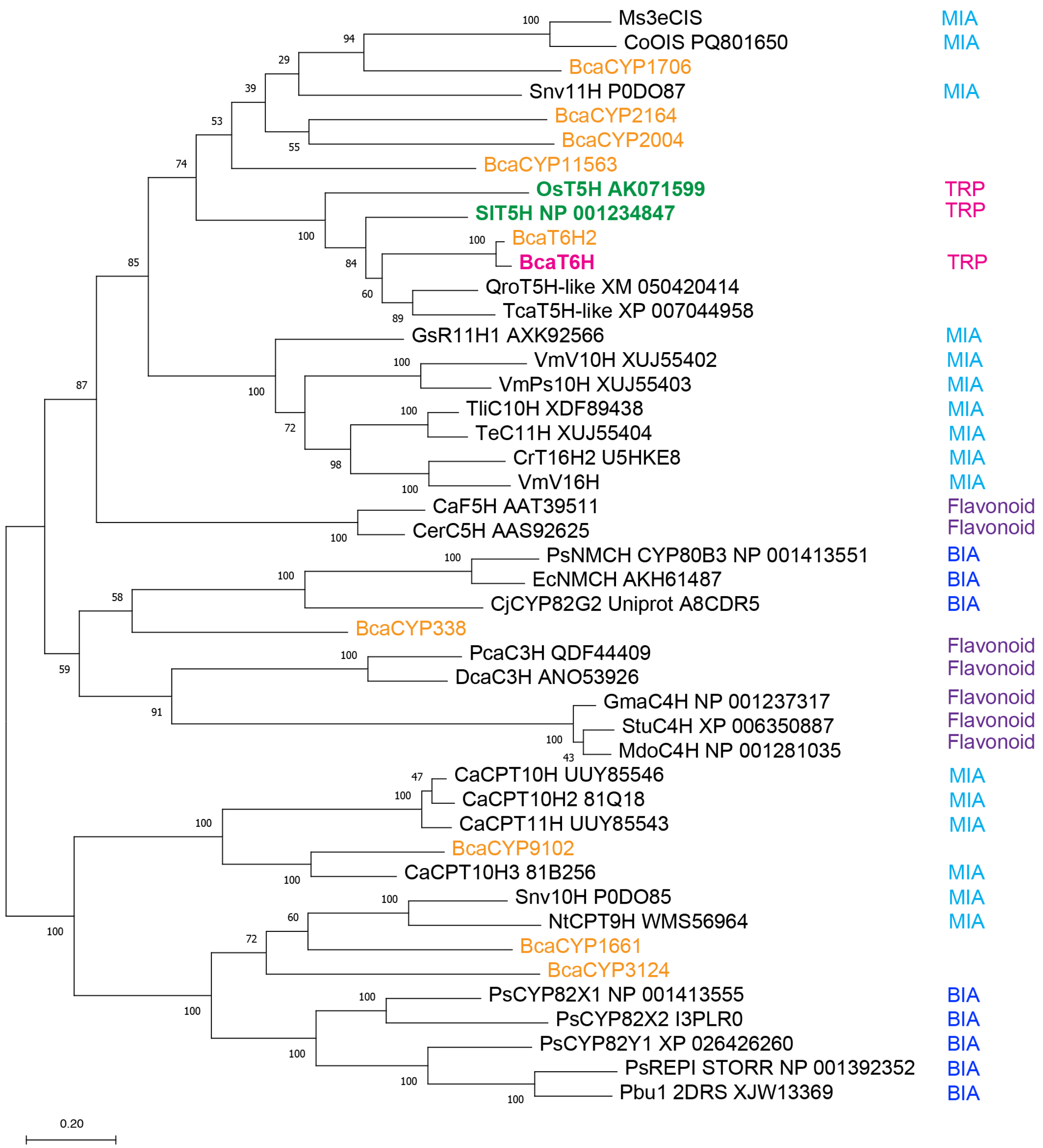


**Figure S23. Phylogenetic analysis of *Banisteriopsis caapi* tryptamine 6-hydroxylase (BcaT6H) protein sequence among other characterized indole alkaloid hydroxylases and benzylisoquinoline alkaloid hydroxylases, related to Figure 6.** The enzymes investigated in this study are color-labeled. Phylogenetically, BcaT6H and its homolog BcaT6H2 (95% amino acid identity) are closely related with two biochemically characterized tryptamine 5-hydroxylases (T5H, labeled in green) from *Solanum lycopersicum* (tomato) and *Oryza sativa* (rice), sharing 65% and 53% amino acid identity, respectively. The classes of metabolites catalyzed by the biochemically characterized cytochrome P450 monooxygenases are labeled as follows: MIA, monoterpenoid indole alkaloids; TRP, tryptamine derivatives; BIA, benzylisoquinoline alkaloids; and flavonoids. The evolutionary history was inferred by using the Maximum Likelihood method and JTT matrix-based model. The percentage of trees in which the associated taxa clustered together is shown next to the branches (100 bootstrap replicates). Initial tree(s) for the heuristic search were obtained automatically by applying Neighbor-Join and BioNJ algorithms to a matrix of pairwise distances estimated using the JTT model, and then selecting the topology with superior log likelihood value. The tree is drawn to scale, with branch lengths measured in the number of substitutions per site (scale bar). Bca: *Banisteriopsis caapi*; Ca: *Camptotheca acuminata*; Cj: *Coptis chinensis/japonica*; Co: *Cephalanthus occidentalis*; Cr: *Catharanthus roseus*; Dca: *Daucus carota*; Ec: *Eschscholzia californica*; Gma: *Glycine max*; Gs: *Gelsemium sempervirens*; Mdo: *Malus domestica*; Ms: *Mitragyna speciosa*; Nt: *Nothapodytes tomentosa*; Os: *Oryza sativa*; Pbu: *Papaver burseri*; Pca: *Phacelia campanularia*; Ps: *Papaver somniferum*; Qro: *Quercus robur*; Sl: *Solanum lycopersicum*; Snv: *Strychnos nux-vomica*; Stu: *Solanum tuberosum*; Tca: *Theobroma cacao*; Te: *Tabernaemontana elegans*; Tli: *Tabernaemontana litoralis*; Vm: *Vinca minor*. The plant species, amino acid sequences are included in data S1.

**A**

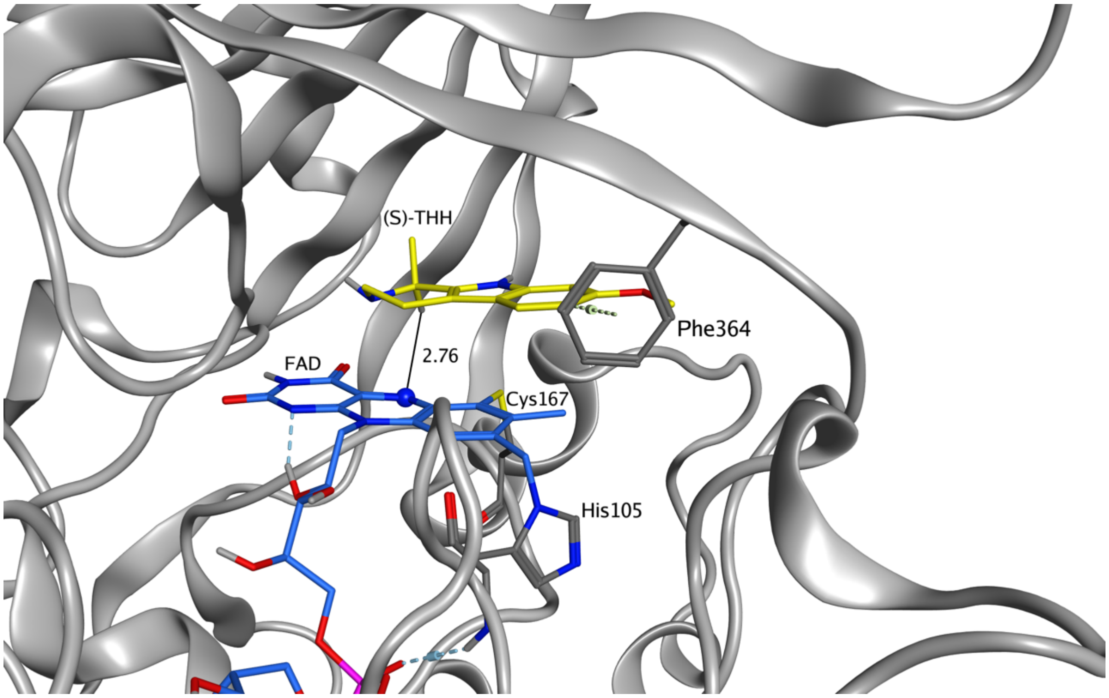


**B**

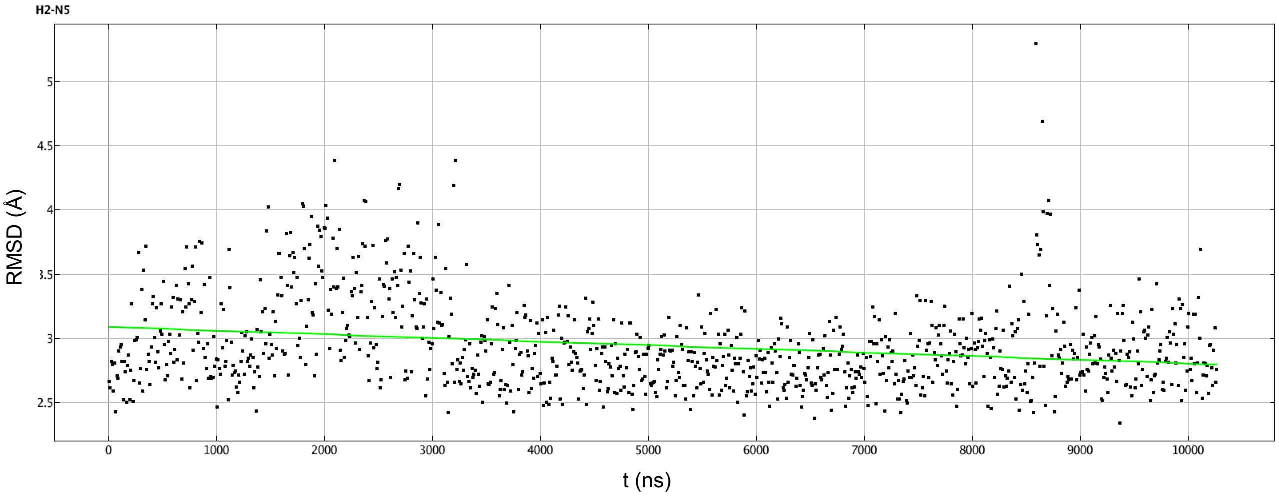


**C**

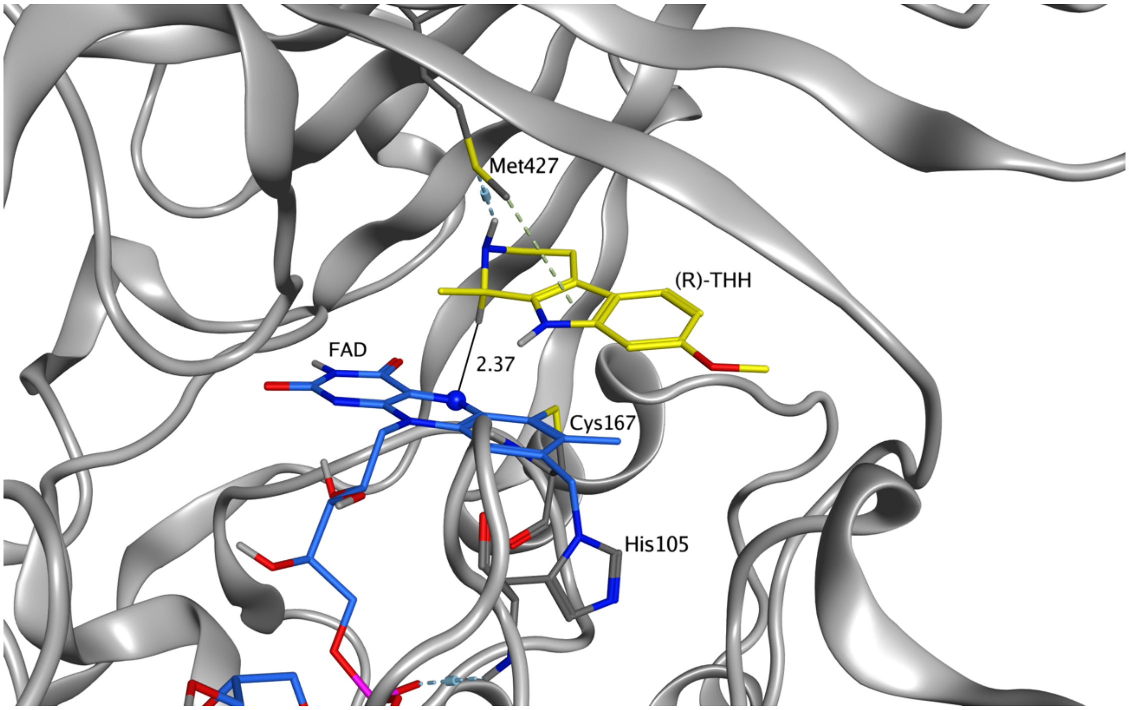


**D**

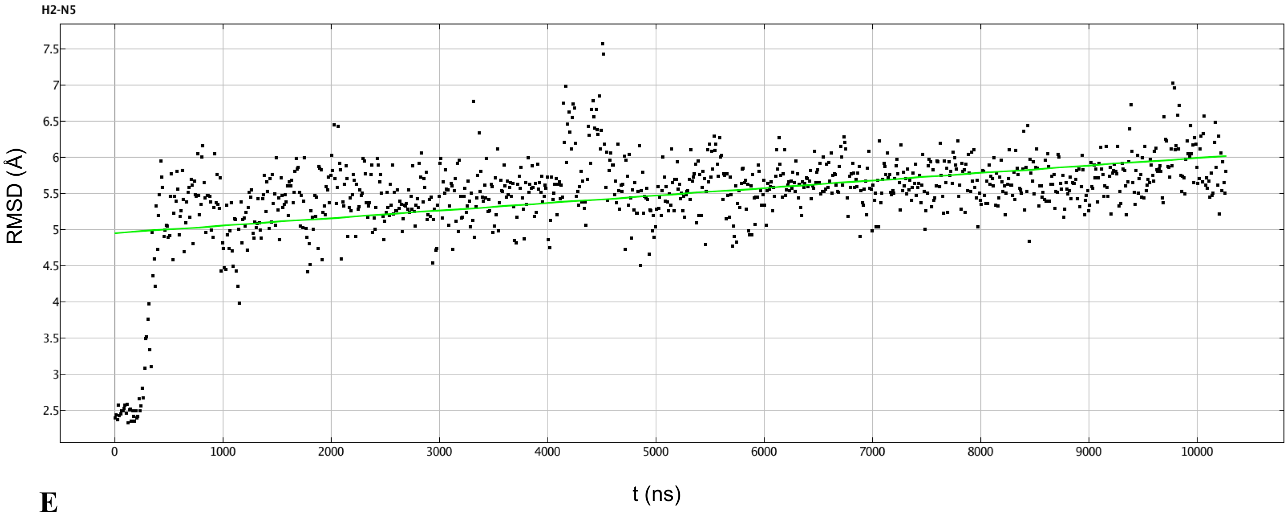


**E**

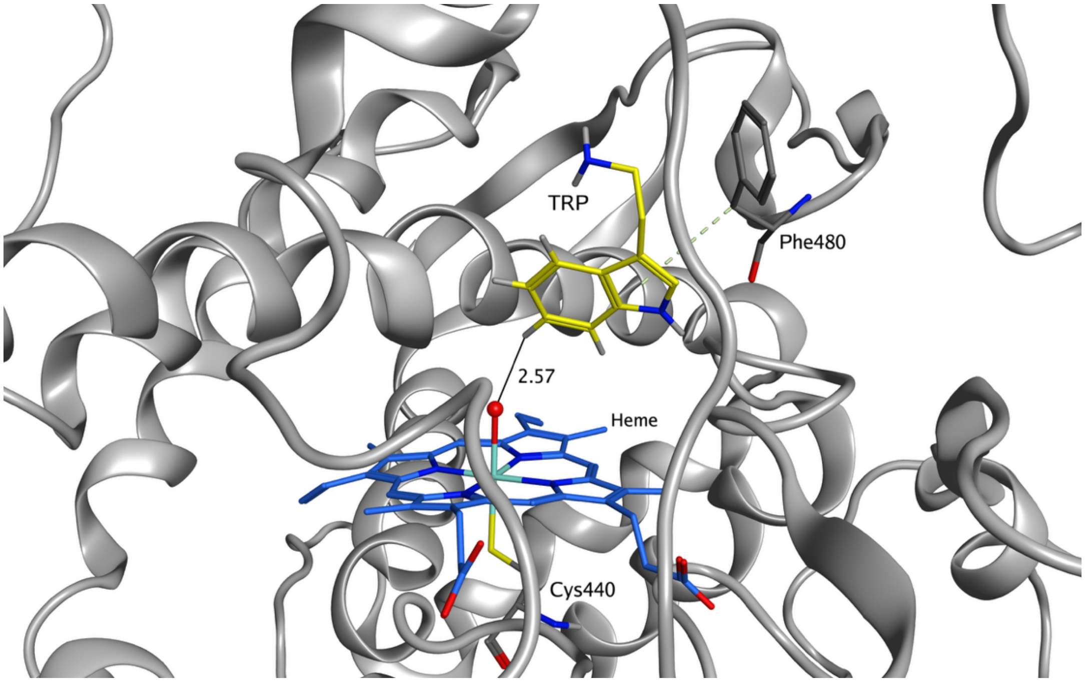


F

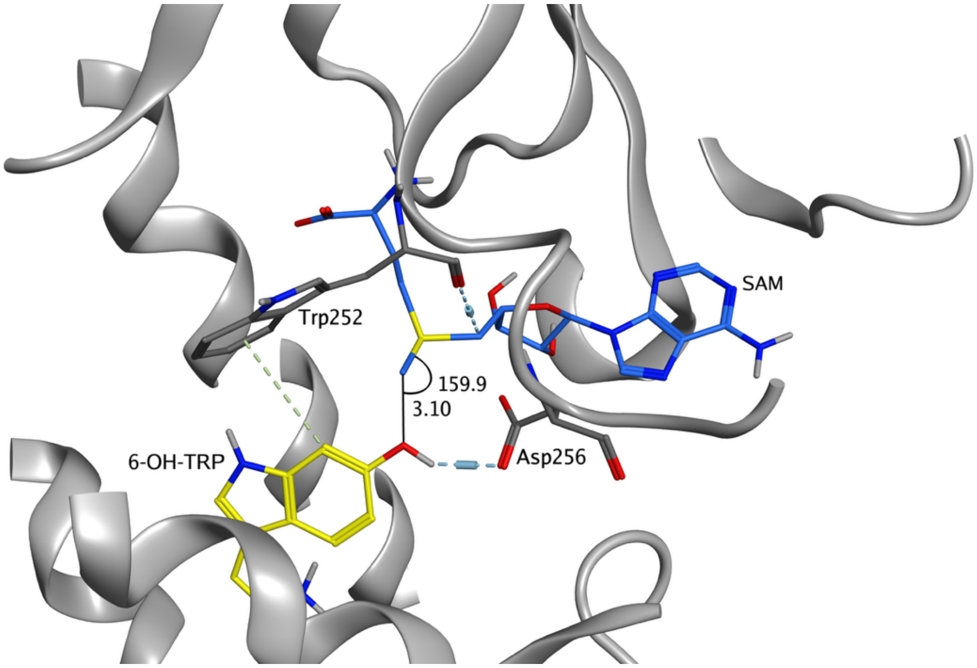


**Figure S24. Homology modeling and molecular dynamics (MD) analyses of *Banisteriopsis caapi* harmaline synthase (BcaHS), tryptamine 6-hydroxylase (BcaT6H), and 6-hydroxytryptamine O-methyltransferase (Bca6OMT) reveal enzyme enantio- and regio-selectivity, related to Figures 6 and 7. (A)** The most favorable binding pose of *S*-tetrahydroharmine (THH) in the active site of BcaHS. *S*-THH π-stacks with the flavin adenine dinucleotide (FAD) cofactor, positioning the C2 hydrogen toward the reactive N5 of FAD (2.76 Å) for hydride abstraction. The FAD is covalently linked to His105 and Cys167, a hallmark of berberine bridge enzyme (BBE)-like oxidases. An excellent docking score (−6.3731 kcal/mol), ranked first among the top 25 poses, indicates optimal binding. **(B)** MD simulation of the optimal *S*-THH binding pose at the BcaHS active site over 10,000 ns. The root mean square deviation (RMSD) remains consistently around 3 Å throughout the simulation, indicating a stable binding conformation. **(C)** Although a binding pose conducive to hydride abstraction can be identified for *R*-THH in the BcaHS active site, it displays a weaker docking score (−5.8509) and ranks 11$^{th}$ among the top 25 poses. **(D)** MD simulation of *R*-THH binding reveals instability of this pose, as the RMSD rapidly increases beyond 5 Å after ~300 ns, suggesting insufficient stability to support catalysis. These modeling results are consistent with biochemical data showing that BcaHS oxidizes only approximately half of a racemic THH mixture, supporting *S*-THH as the preferred substrate. **(E)** Docking of tryptamine (TRP) into the BcaT6H active site reveals optimal positioning of the C6–H bond toward the ferryl oxo (Fe=O) moiety of the heme cofactor, consistent with regio-selective hydroxylation at the C6 position. **(F)** Docking of 6-hydroxytryptamine (6-OH-TRP) into the Bca6OMT active site reveals regio-selectivity for *O*-methylation at the 6-hydroxy group. Asp256 forms a hydrogen bond with the 6-hydroxy group and is positioned to function as a catalytic base during nucleophilic attack on the sulfonium methyl group of the *S*-adenosyl-L-methionine (SAM) cofactor. The observed distance (3.10 Å) and dihedral angle (159.9°) between the substrate oxygen and the SAM methyl group are conducive to an optimal $S_N2$ reaction.

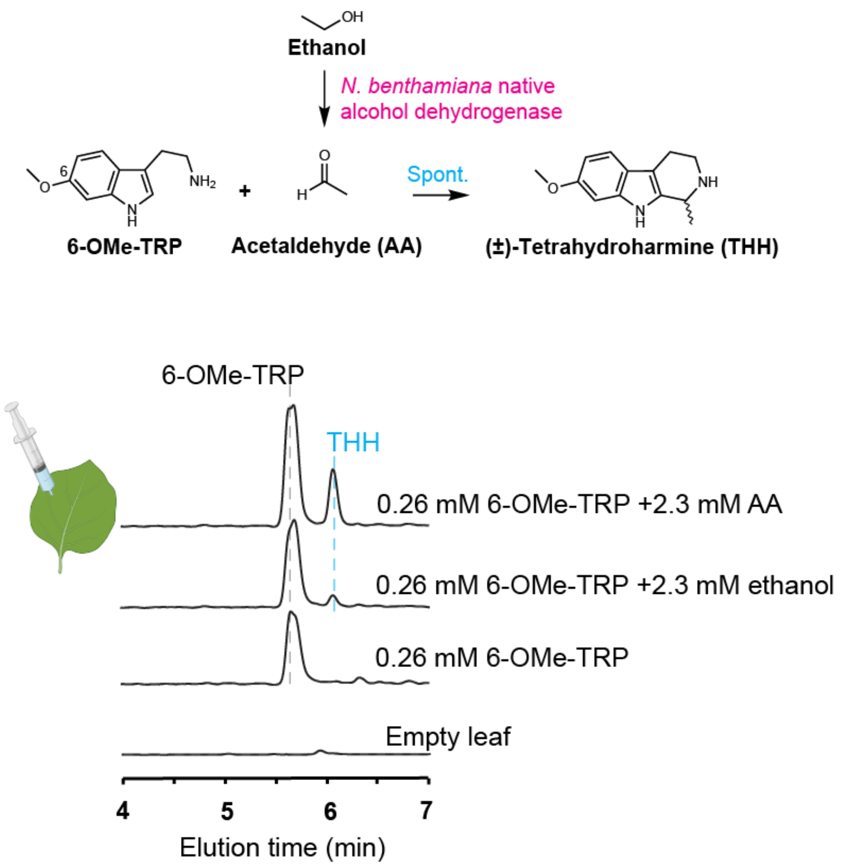


**Figure S25. Co-infiltration of 6-methoxy-tryptamine with acetaldehyde (AA) or ethanol in *Nicotiana benthamiana* leaves results in tetrahydroharmine (THH) formation, related to Figure 6.** Infiltration of 6-methoxytryptamine alone (0.26 mM in 20 mM Tris-HCl, pH 7.5) into *N. benthamiana* leaves did not yield detectable THH. In contrast, co-infiltration with either acetaldehyde or ethanol (2.3 mM in 20 mM Tris-HCl, pH 7.5) resulted in THH formation, supporting a spontaneous reaction between 6-methoxytryptamine and acetaldehyde without the requirement for a *B. caapi* enzyme. Metabolites were detected one day after infiltration. LC-MS/MS analysis shows extracted-ion chromatograms (EICs) corresponding to the following $[M+H]^+$ ions: *m/z* 191 for 6-methoxytryptamine and *m/z* 217 for THH.

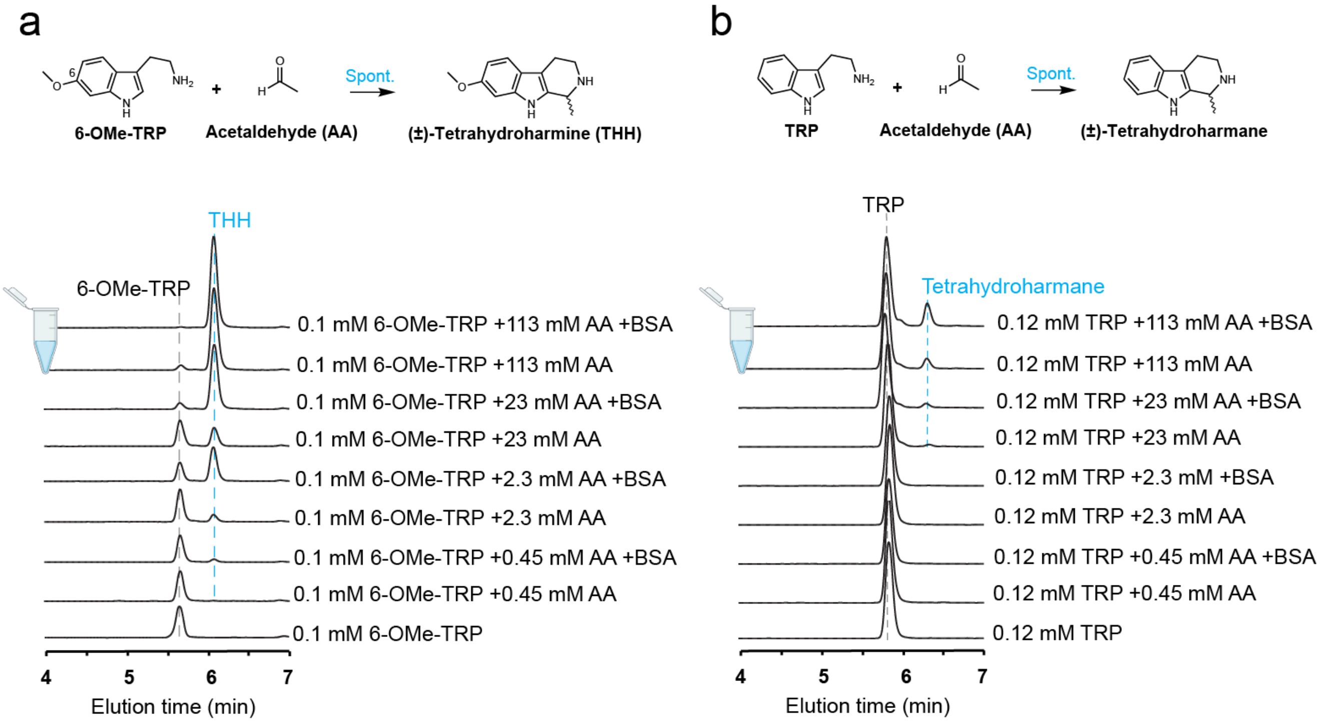


**Figure S26. Acetaldehyde spontaneously reacts with 6-methoxytryptamine or tryptamine in 20 mM Tris HCl (pH 7.5) to form racemic tetrahydroharmine (THH) or tetrahydroharmane, related to Figure 6. (A)** In vitro assays show that THH forms spontaneously upon incubation of 6-methoxytryptamine with acetaldehyde (AA). The amount of THH produced positively correlates with the concentration of acetaldehyde (0.45-113 mM). THH formation was substantially enhanced by the addition of bovine serum albumin (BSA; 250 µg protein per 100 µL assay or 37.6 µM), suggesting reaction rate acceleration through nonspecific protein binding. **(B)** Tetrahydroharmane (eleagnine) formed in a similar manner in in vitro assays, although tryptamine was less reactive than 6-methoxytryptamine, possibly due to the electron-donating effect of the methoxy group increasing indole nucleophilicity. All reactions were incubated at 30 °C overnight to ensure completion. LC-MS/MS analysis shows extracted-ion chromatograms (EICs) corresponding to the following $[M+H]^+$ ions: *m/z* 161 for tryptamine, 191 for 6-methoxytryptamine, *m/z* 187 for tetrahydroharmane, and *m/z* 217 for THH.

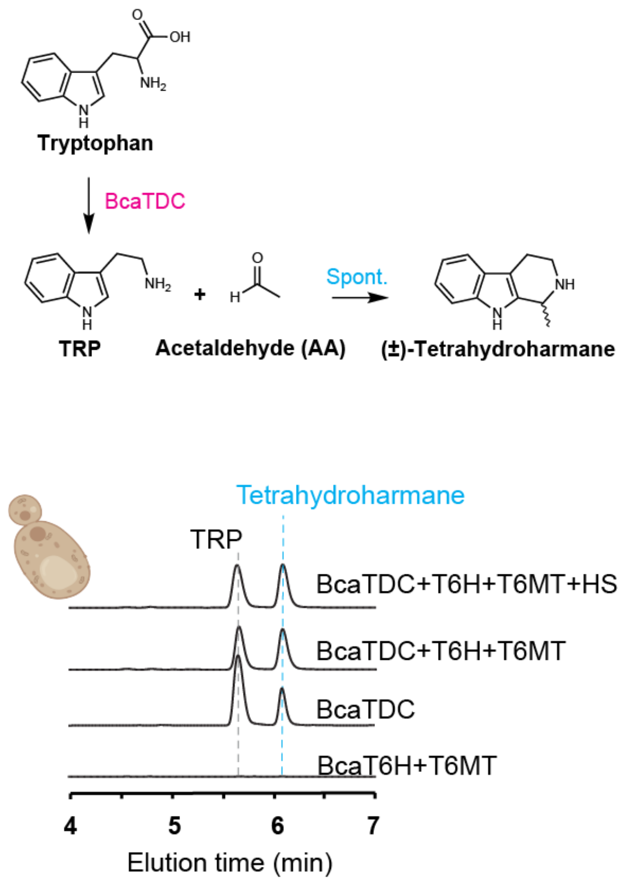


**Figure S27. Tetrahydroharmane forms de novo during yeast fermentation when *Banisteriopsis caapi* tryptophan decarboxylase (BcaTDC) is expressed, related to Figure 6.** LC-MS/MS analysis shows extracted-ion chromatograms (EICs) corresponding to the following $[M+H]^+$ ions: *m/z* 161 for tryptamine and *m/z* 187 for tetrahydroharmane.

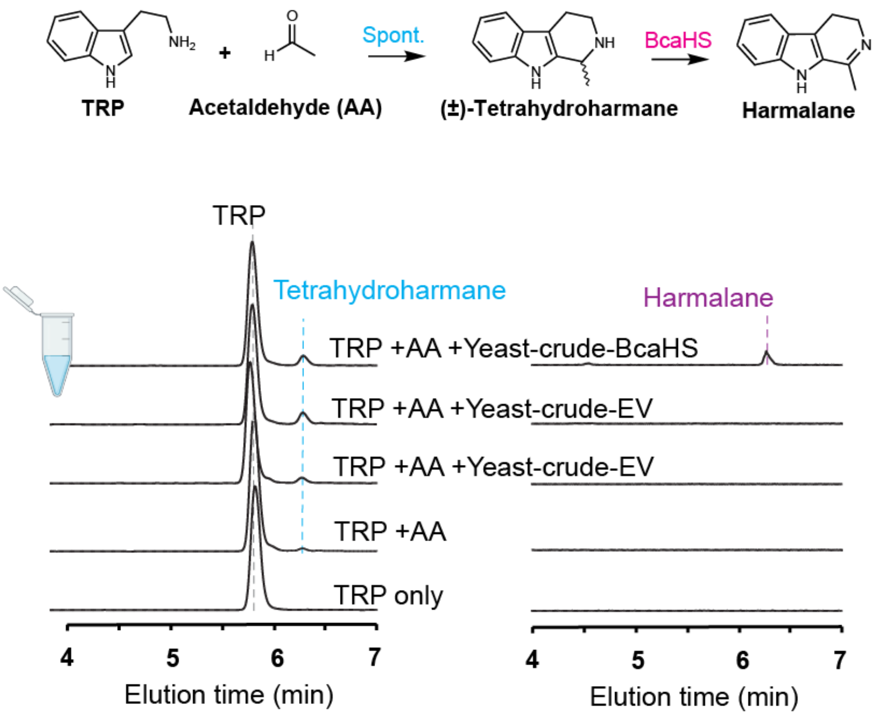


**Figure S28.** ***Banisteriopsis caapi*** **harmaline synthase (BcaHS) oxidizes tetrahydroharmane to harmalane, related to Figure 7.** In vitro assays show that racemic tetrahydroharmane (eleagnine) forms spontaneously upon incubation of tryptamine (TRP) with acetaldehyde (AA), and its formation is substantially enhanced by the addition of yeast crude lysate. Approximately half of the racemic tetrahydroharmane was oxidized to harmalane by yeast crude lysate expressing BcaHS, consistent with the stereoselectivity of BcaHS. All reactions were incubated at 30 °C overnight to ensure completion. LC-MS/MS analysis shows extracted-ion chromatograms (EICs) corresponding to the following $[M+H]^+$ ions: *m/z* 161 for tryptamine and *m/z* 187 for tetrahydroharmane.

**Table S1. Assembly statistics for the *Psychotria viridis* and *Banisteriopsis caapi* genomes, related to Figures 1 and 2.**

| **Feature** | ***Psychotria viridis*** | ***Banisteriopsis caapi* Tucunacá** | ***Banisteriopsis caapi* Caupuri** |
|---|---|---|---|
| **Ploidy** | hexaploid | diploid | diploid |
| **2n / base x** | 66 / 11 | 20 / 10 | 20 / 10 |
| **GenomeScope estimated monoploid size (Gb)** | 1.47 | 1.20 | 1.16 |
| **Heterozygosity (%)** | 8.24 | 1.18 | 0.51 |
| **Total assembly (Gb)** | 8.80 | 2.25 | 2.24 |
| **Assembly span (Gb)** | 1.39-1.51 | 1.10-1.15 | 1.11-1.13 |
| **Number of contigs** | 2649 | 1896 | 1118 |
| **Contig N50** | 19298921 | 4521368 | 11687942 |
| **Contig L50** | 135 | 144 | 54 |
| **Number of scaffolds** | 1952 | 1110 | 737 |
| **Longest scaffold (bp)** | 181506585 | 129066968 | 125969259 |
| **Scaffold N50** | 142595911 | 109940244 | 113503008 |
| **Scaffold L50** | 29 | 10 | 10 |
| **Pseudochromosome size (Gb)** | 8.67 | 2.20 | 2.20 |
| **Anchor rate (%)** | 98.54 | 97.51 | 98.22 |
| **Gap number** | 698 | 787 | 381 |
| **Gap span (bp)** | 69800 | 78700 | 38200 |
| **BUSCO lineage** | embryophyta_odb12 | embryophyta_odb12 | embryophyta_odb12 |
| **BUSCO completeness (%)** | 98.7 | 98.9 | 98.8 |
| **BUSCO completeness for each monoploid/haplotype (%)** | 86.4-94.8 | 98.2 | 98.2-98.3 |
| **Merqury completeness** | 98.33 | 99.22 | 99.30 |
| **Merqury QV** | 56.44 | 62.64 | 63.57 |
| **LAI** | 18.38 | 13.77 | 13.62 |
| **Repeat content** | 70.86 | 56.73 | 57.00 |
| **Total gene counts** | 237279 | 81183 | 88266 |

### Table S2. Per-monoploid and per-haplotype assembly statistics, related to Figures 1 and 2.

| Feature | *P. viridis* g1 | *P. viridis* g2 | *P. viridis* g3 | *P. viridis* g4 | *P. viridis* g5 | *P. viridis* g6 | *B. caapi* TCN hap1 | *B. caapi* TCN hap2 | *B. caapi* CPR hap1 | *B. caapi* CPR hap2 |
|---|---|---|---|---|---|---|---|---|---|---|
| **Assembly (Gb)** | 1.51 | 1.47 | 1.45 | 1.44 | 1.42 | 1.39 | 1.15 | 1.10 | 1.13 | 1.11 |
| **Number of scaffolds** | 11 | 11 | 11 | 11 | 11 | 11 | 939 | 171 | 633 | 104 |
| **Longest scaffold (bp)** | 174156282 | 164838164 | 162997396 | 160591913 | 162741385 | 181506585 | 129066968 | 124841945 | 125152041 | 125969259 |
| **Scaffold N50** | 147588419 | 144108249 | 142595911 | 143371387 | 138237763 | 139442786 | 112484948 | 109940244 | 113503008 | 114320970 |
| **Scaffold L50** | 5 | 5 | 5 | 5 | 5 | 5 | 5 | 5 | 5 | 5 |
| **Gap number** | 130 | 106 | 114 | 121 | 107 | 120 | 417 | 370 | 201 | 180 |
| **Gap span (bp)** | 13000 | 10600 | 11400 | 12100 | 10700 | 12000 | 41700 | 37000 | 20200 | 18000 |
| **BUSCO lineage** | embryophyta_odb12 | | | | | embryophyta_odb12 | | | | |
| **BUSCO completeness (%)** | 93.0 | 94.0 | 94.6 | 94.8 | 93.7 | 86.4 | 98.2 | 98.2 | 98.2 | 98.3 |
| **BUSCO single copy (%)** | 80.8 | 89.9 | 90.8 | 89.7 | 90.1 | 81.7 | 93.7 | 93.7 | 93.4 | 93.5 |
| **LAI** | 17.28 | 18.51 | 19.2 | 18.31 | 18.52 | 18.52 | 13.74 | 13.8 | 13.65 | 13.59 |
| **Repeat content (%)** | 70.96 | 70.38 | 70.39 | 70.9 | 71.12 | 71.46 | 56.45 | 57.02 | 56.8 | 57.21 |
| **Gene counts** | 38221 | 36438 | 36046 | 36759 | 35628 | 32978 | 41019 | 40164 | 47343 | 40923 |

**Table S4. The most highly expressed tryptophan decarboxylase (TDC) and methyltransferase (MT) candidates in *Psychotria viridis* young leaf tissue measured in Transcript Per Million reads (TPM), related to Figure 5.**

| Query | Gene | Young Leaf TPM | Mature Leaf1 TPM | Mature Leaf 2 TPM | Mature leaf 3 TPM | Stem 1 TPM | Stem 2 TPM | Root 1 TPM | Root 2 TPM |
|---|---|---|---|---|---|---|---|---|---|
| TRINITY_DN1174_c0_g2_i1 | PviTDC | 64.49 | 0.65 | 0.89 | 0 | 0 | 0 | 0 | 0 |
| TRINITY_DN213_c0_g1_i1 | PviTNMT | 606.84 | 267.83 | 270.87 | 23.99 | 5.93 | 2.36 | 2.44 | 2.15 |
| TRINITY_DN4304_c0_g1_i9 | PviMT4304 | 52.77 | 9.90 | 9.60 | 4.09 | 3.16 | 0.90 | 13.19 | 15.32 |
| TRINITY_DN88_c0_g1_i5 | PviMT88 | 33.49 | 29.82 | 29.98 | 29.47 | 1.99 | 3.28 | 2.87 | 2.44 |
| TRINITY_DN20512_c0_g1_i1 | PviMT20512 | 8.39 | 110.40 | 121.33 | 21.32 | 399.67 | 387.87 | 43.49 | 50.57 |
| TRINITY_DN7221_c0_g1_i8 | PviMT7221 | 7.38 | 16.02 | 13.70 | 14.92 | 12.04 | 13.89 | 66.39 | 94.41 |
| TRINITY_DN36665_c0_g1_i3 | PviMT36665 | 4.56 | 7.16 | 6.83 | 9.06 | 3.55 | 3.72 | 17.22 | 18.83 |

**Table S5. Kinetics parameters of *Psychotria viridis* tryptamine *N*-methyltransferases (PviTNMT) and a characterized NMT from *Rhinella marina* (cane toad), related to Figure 5.**

| | Substrate | $K_M$ (μM) | $K_M$ 95% CI* (μM) | $k_{cat}$ ($10^{-3}$ $s^{-1}$) | $k_{cat}$ 95% CI ($s^{-1}$) | $k_{cat}$ / $K_M$ ($M^{-1}$ $s^{-1}$) |
|---|---|---|---|---|---|---|
| PviTNMT | TRP | 4.56 | 3.76 - 5.52 | 28.8 | 36.8 - 41.1 | 6312.2 |
| | MMT | 100.4 | 90.19 - 112.1 | 0.840 | 1.08 - 1.20 | 8.40 |
| RmaTNMT** | TRP | 96.3 ± 12.8 | - | 0.520 | - | 5.40 |
| | MMT | 37.0 ± 9.5 | - | 2.10 | - | 56.8 |

*Confidence interval

** Data are adapted from Chen et al.[S1]

**Table S6. The 21 most highly expressed transcripts in *Banisteriopsis caapi* young leaf tissue that share less than 70% amino acid identity with *Arabidopsis thaliana* homologs, related to Figure 6. Expression levels are reported as Transcripts Per Million (TPM).**

| | | *B. caapi* var. Caupuri | | | | | | *B. caapi* var. Tucunaca | | | | | |
|---|---|---|---|---|---|---|---|---|---|---|---|---|---|
| Query | gene | Young Leaf TPM | Mature Leaf 1 TPM | Mature Leaf 2 TPM | Mature Leaf 3 TPM | Stem 1 TPM | Stem 2 TPM | Young Leaf TPM | Mature Leaf 1 TPM | Mature Leaf 2 TPM | Mature Leaf 3 TPM | Stem 1 TPM | Stem 2 TPM |
| TRINITY_DN28178_c0_g1_i1 | Lipid transfer protein | 3077.5 | 393.4 | 1016.0 | 1030.0 | 118.5 | 176.6 | 2186.0 | 717.6 | 1057.0 | 1050.6 | 32.8 | 26.4 |
| TRINITY_DN1623_c0_g1_i2 | BcaTDC | 1866.2 | 0.0 | 1.3 | 0.6 | 197.8 | 166.1 | 468.2 | 0.0 | 0.0 | 0.0 | 329.0 | 307.5 |
| TRINITY_DN2205_c0_g2_i5 | Bca6OMT | 1544.6 | 2.4 | 23.8 | 30.2 | 174.4 | 160.1 | 397.5 | 8.2 | 12.7 | 13.3 | 303.2 | 322.6 |
| TRINITY_DN5042_c0_g2_i1 | Pathogenesis-related 4 | 1309.8 | 0.0 | 1.7 | 2.0 | 178.8 | 193.1 | 1234.4 | 1.4 | 2.0 | 1.9 | 410.5 | 376.5 |
| TRINITY_DN9498_c0_g1_i11 | Lipoxygenase | 1117.0 | 1799.6 | 630.9 | 615.3 | 250.1 | 251.7 | 1596.3 | 1917.5 | 581.4 | 567.3 | 192.6 | 196.0 |
| TRINITY_DN4481_c0_g1_i13 | Dioxygenase | 790.5 | 25.9 | 339.8 | 327.6 | 294.5 | 293.0 | 611.6 | 235.4 | 73.2 | 77.2 | 445.0 | 456.6 |
| TRINITY_DN546_c0_g2_i1 | NADPH-dependent reductase | 737.5 | 42.1 | 341.6 | 351.1 | 275.9 | 285.1 | 481.8 | 255.2 | 173.2 | 175.8 | 452.0 | 485.4 |
| TRINITY_DN195_c0_g1_i4 | Anthocyanidin reductase like | 694.1 | 85.1 | 1058.4 | 1064.6 | 845.0 | 938.7 | 573.0 | 442.8 | 428.6 | 432.7 | 1129.1 | 1154.4 |
| TRINITY_DN1720_c0_g1_i21 | Very-long-chain aldehyde decarbonylase-like | 608.9 | 62.7 | 0.0 | 0.2 | 1.0 | 1.0 | 880.8 | 112.8 | 0.2 | 0.9 | 0.0 | 0.0 |
| TRINITY_DN30017_c0_g1_i1 | BcaHS | 603.7 | 16.8 | 32.3 | 31.4 | 275.0 | 272.5 | 516.1 | 21.9 | 18.8 | 15.4 | 461.3 | 458.2 |
| TRINITY_DN7_c0_g2_i2 | Photosystem_II_reaction_center_W | 591.6 | 828.9 | 978.4 | 979.7 | 229.7 | 240.0 | 564.8 | 310.9 | 677.0 | 658.2 | 182.5 | 193.8 |
| TRINITY_DN14910_c1_g1_i2 | Acid_phosphatase | 577.1 | 1.9 | 2.2 | 3.5 | 8.5 | 19.2 | 150.7 | 2.3 | 2.3 | 2.7 | 0.8 | 2.0 |
| TRINITY_DN89_c0_g1_i2 | Protodermal_factor_1 | 570.5 | 0.0 | 0.0 | 0.0 | 0.8 | 0.0 | 28.9 | 0.4 | 0.0 | 0.0 | 0.0 | 0.0 |
| TRINITY_DN435_c0_g1_i1 | Unknown protein | 535.0 | 1318.9 | 831.2 | 863.1 | 2889.6 | 2759.9 | 1129.9 | 864.6 | 1968.1 | 1975.1 | 1698.6 | 1631.0 |
| TRINITY_DN4230_c0_g1_i1 | Photosystem_I_reaction_center_subunit | 475.5 | 710.3 | 1169.1 | 1193.5 | 434.4 | 423.2 | 448.0 | 128.7 | 565.1 | 551.7 | 353.1 | 338.4 |
| TRINITY_DN33_c0_g3_i1 | Sulfotransferase-like | 454.5 | 305.7 | 180.5 | 193.8 | 24.8 | 11.7 | 228.9 | 343.4 | 76.8 | 73.6 | 0.6 | 0.6 |
| TRINITY_DN5322_c0_g1_i6 | Fe_superoxide_dismutase_2 | 368.8 | 1188.8 | 196.3 | 210.9 | 48.8 | 67.4 | 4.8 | 15.9 | 94.8 | 97.1 | 46.1 | 41.7 |
| TRINITY_DN13887_c0_g1_i3 | BcaT6H | 351.2 | 24.4 | 118.4 | 120.0 | 207.0 | 195.6 | 206.1 | 143.2 | 80.5 | 83.8 | 321.4 | 338.2 |
| TRINITY_DN9837_c0_g3_i2 | Pollen_Ole_e_1_allergen | 336.6 | 17.9 | 56.2 | 49.8 | 355.5 | 291.5 | 266.3 | 35.4 | 31.2 | 30.2 | 350.3 | 360.4 |
| TRINITY_DN575_c0_g1_i13 | BcaMT575 | 328.6 | 901.1 | 1189.2 | 1194.5 | 287.9 | 298.0 | 739.1 | 787.9 | 652.1 | 646.9 | 167.1 | 136.5 |
| TRINITY_DN2168_c1_g1_i1 | Major latex protein | 317.3 | 24.7 | 56.1 | 63.5 | 14.7 | 22.2 | 162.9 | 42.0 | 64.2 | 61.2 | 20.9 | 19.5 |

**Table S7. CENH3/H3 enrichment over TRC arrays versus CRM elements within satellite-containing centromeres (chromosomes 3, 5 and 10) in *Banisteriopsis caapi* Tucunacá haplotype 1, related to Figure 3.**

| Chromosome | TRC family | n (TRC) | Median CENH3/H3 (TRC) | n (CRM) | Median CENH3/H3 (CRM) | W | p |
|---|---|---|---|---|---|---|---|
| chr3_h1 | TRC1 | 618 | 0.691 | 89 | 0.420 | 15,788 | $7.94 \times 10^{-11}$ |
| chr5_h1 | TRC2 | 298 | 0.778 | 50 | 0.463 | 4,218 | $9.16 \times 10^{-7}$ |
| chr10_h1 | TRC4 | 171 | 0.782 | 80 | 0.410 | 4,258 | $1.46 \times 10^{-6}$ |

Median CENH3/H3 log2 ratios and Wilcoxon rank-sum test statistics (W, two-sided p) for 1-kb windows classified by repeat element within annotated centromere regions; n, number of 1-kb windows per element class.

**Table S8. CENH3/H3 enrichment over CRM versus LINE elements within the centromeres of all chromosomes in *Banisteriopsis caapi* Tucunacá haplotype 1, related to Figure 3 and Figure S11.**

| Chromosome | n (CRM) | Median CENH3/H3 (CRM) | n (LINE) | Median CENH3/H3 (LINE) | W | p |
|---|---|---|---|---|---|---|
| chr1_h1 | 341 | 0.389 | 218 | 0.983 | 16,570 | $1.98 \times 10^{-28}$ |
| chr2_h1 | 286 | 1.137 | 246 | 1.127 | 36,226 | 0.553 |
| chr3_h1 | 89 | 0.420 | 74 | 0.225 | 4,222 | $1.97 \times 10^{-3}$ |
| chr4_h1 | 242 | 1.292 | 128 | 1.366 | 15,191 | 0.762 |
| chr5_h1 | 50 | 0.463 | 42 | 0.342 | 1,339 | 0.0237 |
| chr6_h1 | 224 | 0.895 | 175 | 0.868 | 20,833 | 0.281 |
| chr7_h1 | 192 | 1.309 | 267 | 1.069 | 34,321 | $5.72 \times 10^{-10}$ |
| chr8_h1 | 139 | 1.284 | 173 | 1.338 | 12,708 | 0.388 |
| chr9_h1 | 301 | 1.095 | 368 | 0.805 | 59,007 | 0.145 |
| chr10_h1 | 80 | 0.410 | 147 | 0.271 | 7,060 | 0.0126 |

Median CENH3/H3 log2 ratios and Wilcoxon rank-sum test statistics (W, two-sided p) for 1-kb windows classified by repeat element within annotated centromere regions; n, number of 1-kb windows per element class.

**Table S3** (collection metadata for the 135 resequenced *Banisteriopsis caapi* accessions and the two reference genotypes, related to Figure 4) and **Data S1** (amino acid sequences used in phylogenetic analysis, related to Figures S16 and S23) are provided as separate files.
