## Supplementary material for "Ayahuasca plant genomes reveal retrotransposon-derived centromeres and contrasting routes to DMT and β-carbolines": Table S3

**Table S3. Collection metadata for all Banisteriopsis caapi samples included in the population genetic analysis (135 resequenced accessions and the TCN and CPR reference genotypes; 137 samples), related to Figure 4.**

| Sample | Locality | Coordinate | Collection year | Geographic location | Country | Variety |
|---|---|---|---|---|---|---|
| NE-01 | NE | 9.645597 S 67.112535 W | 2020 | Lábrea, Amazonas | Brazil | Tucunacá |
| NE-02 | NE | 9.644922 S 67.11991 W | 2020 | Lábrea, Amazonas | Brazil | Tucunacá |
| NE-03 | NE | 9.658993 S 67.116372 W | 2020 | Lábrea, Amazonas | Brazil | Tucunacá |
| NE-04 | NE | 9.664152 S 67.112901 W | 2020 | Lábrea, Amazonas | Brazil | Tucunacá |
| OP-01 | OP | 10.595218 S 62.323018 W | 2020 | Ouro Preto do Oeste, Rondônia | Brazil | Tucunacá |
| OP-02 | OP | 10.595305 S 62.322888 W | 2020 | Ouro Preto do Oeste, Rondônia | Brazil | Tucunacá |
| OP-03 | OP | 10.593244 S 62.330088 W | 2020 | Ouro Preto do Oeste, Rondônia | Brazil | Tucunacá |
| OP-04 | OP | 10.593447 S 62.330064 W | 2020 | Ouro Preto do Oeste, Rondônia | Brazil | Tucunacá |
| OP-05 | OP | 10.589639 S 62.326173 W | 2020 | Ouro Preto do Oeste, Rondônia | Brazil | Tucunacá |
| OP-06 | VP | 10.487775 S 62.061316 W | 2020 | Vale do Paraíso, Rondônia | Brazil | Tucunacá |
| OP-07 | VP | 10.487678 S 62.061344 W | 2020 | Vale do Paraíso, Rondônia | Brazil | Tucunacá |
| OP-08 | VP | 10.487651 S 62.06143 W | 2020 | Vale do Paraíso, Rondônia | Brazil | Tucunacá |
| OP-09 | VP | 10.487991 S 62.060982 W | 2020 | Vale do Paraíso, Rondônia | Brazil | Tucunacá |
| OP-10 | VP | 10.487992 S 62.060889 W | 2020 | Vale do Paraíso, Rondônia | Brazil | Tucunacá |
| PF-01 | PF | 1.772453 S 60.251982 W | 2020 | Presidente Figueiredo, Amazonas | Brazil | Tucunacá |
| PF-02 | PF | 1.773452 S 60.249068 W | 2020 | Presidente Figueiredo, Amazonas | Brazil | Tucunacá |
| PF-03 | PF | 1.773861 S 60.248986 W | 2020 | Presidente Figueiredo, Amazonas | Brazil | Tucunacá |
| PF-04 | PF | 1.774041 S 60.249073 W | 2020 | Presidente Figueiredo, Amazonas | Brazil | Tucunacá |
| PF-05 | PF | 1.77418 S 60.249114 W | 2020 | Presidente Figueiredo, Amazonas | Brazil | Tucunacá |
| RA-01 | RA | 7.789446 S 72.721101 W | 2020 | Rodrigues Alves, Acre | Brazil | Caupuri |
| RA-02 | RA | 7.789872 S 72.721168 W | 2020 | Rodrigues Alves, Acre | Brazil | Caupuri |
| RA-03 | RA | 7.792041 S 72.722651 W | 2020 | Rodrigues Alves, Acre | Brazil | Caupuri |
| RA-04 | RA | 7.792179 S 72.723708 W | 2020 | Rodrigues Alves, Acre | Brazil | Caupuri |
| RB-01 | EX | 9.737048 S 66.448136 W | 2020 | Extrema, Rondônia | Brazil | Tucunacá |
| RB-02 | EX | 9.74949 S 66.451298 W | 2020 | Extrema, Rondônia | Brazil | Tucunacá |

| | | | | | | |
|---|---|---|---|---|---|---|
| RB-04 | EX | 9.741727 S 66.443393 W | 2020 | Extrema, Rondônia | Brazil | Tucunacá |
| RB-05 | EX | 9.741586 S 66.443875 W | 2020 | Extrema, Rondônia | Brazil | Tucunacá |
| RB-09 | AC | 9.905748 S 66.878441 W | 2020 | Acrelândia, Acre | Brazil | Tucunacá |
| RB-11 | PC | 10.19152 S 67.153715 W | 2020 | Plácido de Castro, Acre | Brazil | Tucunacá |
| RB-12 | PC | 10.190523 S 67.14985 W | 2020 | Plácido de Castro, Acre | Brazil | Tucunacá |
| RB-13 | PC | 10.189584 S 67.152385 W | 2020 | Plácido de Castro, Acre | Brazil | Tucunacá |
| RB-14 | PC | 10.186507 S 67.155034 W | 2020 | Plácido de Castro, Acre | Brazil | Tucunacá |
| RB-15 | PC | 10.187243 S 67.153623 W | 2020 | Plácido de Castro, Acre | Brazil | Tucunacá |
| RB-16 | CX | 10.615314 S 67.989271 W | 2020 | Capixaba, Acre | Brazil | Tucunacá |
| RB-17 | CX | 10.615051 S 67.98944 W | 2020 | Capixaba, Acre | Brazil | Tucunacá |
| RB-18 | CX | 10.615749 S 67.989538 W | 2020 | Capixaba, Acre | Brazil | Tucunacá |
| RB-19 | CX | 10.616143 S 67.990084 W | 2020 | Capixaba, Acre | Brazil | Tucunacá |
| RR-01 | RR | 1.049355 N 59.938753 W | 2020 | São João da Baliza, Roraima | Brazil | Tucunacá |
| RR-03 | RR | 1.049677 N 59.935619 W | 2020 | São João da Baliza, Roraima | Brazil | Tucunacá |
| RR-04 | RR | 1.049865 N 59.937332 W | 2020 | São João da Baliza, Roraima | Brazil | Tucunacá |
| RR-05 | RR | 1.049731 N 59.936261 W | 2020 | São João da Baliza, Roraima | Brazil | Tucunacá |
| SL-01 | SL | 7.756986 S 72.424344 W | 2020 | Santa Luzia, Acre | Brazil | Tucunacá |
| SL-02 | SL | 7.758357 S 72.422821 W | 2020 | Santa Luzia, Acre | Brazil | Tucunacá |
| SL-03 | SL | 7.792389 S 72.392262 W | 2020 | Santa Luzia, Acre | Brazil | Tucunacá |
| SL-04 | SL | 7.792977 S 72.392601 W | 2020 | Santa Luzia, Acre | Brazil | Tucunacá |
| TF-01 | TF | 3.538836 S 64.620749 W | 2020 | Tefé, Amazonas | Brazil | Tucunacá |
| TF-03 | TF | 3.537856 S 64.619456 W | 2020 | Tefé, Amazonas | Brazil | Tucunacá |
| TF-04 | TF | 3.535679 S 64.619961 W | 2020 | Tefé, Amazonas | Brazil | Tucunacá |
| TF-05 | CS | 3.500707 S 64.80326 W | 2020 | Tefé, Amazonas | Brazil | Tucunacá |
| TF-06 | CS | 3.497782 S 64.802991 W | 2020 | Tefé, Amazonas | Brazil | Tucunacá |
| TF-07 | CS | 3.49777 S 64.801501 W | 2020 | Tefé, Amazonas | Brazil | Tucunacá |
| TF-08 | CS | 3.497363 S 64.775906 W | 2020 | Tefé, Amazonas | Brazil | Tucunacá |
| TF-09 | CS | 3.495965 S 64.775247 W | 2020 | Tefé, Amazonas | Brazil | Tucunacá |

| | | | | | |
|---|---|---|---|---|---|
| TF-10 | TF | 3.53099 S 64.624027 W | 2020 Tefé, Amazonas | Brazil | Tucunacá |
| TR-02 | TR | 8.073288 S 71.246915 W | 2020 Tarauacá, Acre | Brazil | Tucunacá |
| Cbá 1 | Cba | 15.447902 S 55.978346 W | 2022 Cuiabá, Mato Grosso | Brazil | |
| Cbá 2 | Cba | 15.447825 S 55.978044 W | 2022 Cuiabá, Mato Grosso | Brazil | |
| Cbá 3 | Cba | 15.447824 S 55.97811 W | 2022 Cuiabá, Mato Grosso | Brazil | |
| Cbá 4 | Cba | 15.447815 S 55.978197 W | 2022 Cuiabá, Mato Grosso | Brazil | |
| Cbá 5 | Cba | 15.447812 S 55.978333 W | 2022 Cuiabá, Mato Grosso | Brazil | |
| Cbá 6 | Cba | 15.447804 S 55.97823 W | 2022 Cuiabá, Mato Grosso | Brazil | |
| Cbá 7 | Cba | 15.447798 S 55.978225 W | 2022 Cuiabá, Mato Grosso | Brazil | |
| Cbá 8 | Cba | 15.447817 S 55.978236 W | 2022 Cuiabá, Mato Grosso | Brazil | |
| Cbá 9 | Cba | 15.447816 S 55.978235 W | 2022 Cuiabá, Mato Grosso | Brazil | |
| Cbá 10 | Cba | 15.447792 S 55.978226 W | 2022 Cuiabá, Mato Grosso | Brazil | |
| Cbá 11 | Cba | 15.447797 S 55.978407 W | 2022 Cuiabá, Mato Grosso | Brazil | |
| Cbá 12 | Cba | 15.447839 S 55.978298 W | 2022 Cuiabá, Mato Grosso | Brazil | |
| Cbá 13 | Cba | 15.448115 S 55.978216 W | 2022 Cuiabá, Mato Grosso | Brazil | |
| Cbá 14 | Cba | 15.447988 S 55.977971 W | 2022 Cuiabá, Mato Grosso | Brazil | |
| Cbá 15 | Cba | 15.448082 S 55.978203 W | 2022 Cuiabá, Mato Grosso | Brazil | |
| Cbá 16 | Cba | 15.447864 S 55.977841 W | 2022 Cuiabá, Mato Grosso | Brazil | |
| Cbá 17 | Cba | 15.448055 S 55.97789 W | 2022 Cuiabá, Mato Grosso | Brazil | |
| Cbá 18 | Cba | 15.448027 S 55.977546 W | 2022 Cuiabá, Mato Grosso | Brazil | |
| Cbá 19 | Cba | 15.448207 S 55.978346 W | 2022 Cuiabá, Mato Grosso | Brazil | |
| Cbá 20 | Cba | 15.448101 S 55.978262 W | 2022 Cuiabá, Mato Grosso | Brazil | |
| Cbá 21 | Cba | 15.447883 S 55.978084 W | 2022 Cuiabá, Mato Grosso | Brazil | |
| Cbá 22 | Cba | 15.447979 S 55.978549 W | 2022 Cuiabá, Mato Grosso | Brazil | |
| Cbá 23 | Cba | 15.447715 S 55.977927 W | 2022 Cuiabá, Mato Grosso | Brazil | |
| Cbá 24 | Cba | 15.44762 S 55.977961 W | 2022 Cuiabá, Mato Grosso | Brazil | |
| Cbá 25 | Cba | 15.447395 S 55.977383 W | 2022 Cuiabá, Mato Grosso | Brazil | |
| Cbá 26 | Cba | 15.447396 S 55.977177 W | 2022 Cuiabá, Mato Grosso | Brazil | |

| | | | | | |
|---|---|---|---|---|---|
| Cbá 27 | Cba | 15.447464 S 55.977974 W | 2022 | Cuiabá, Mato Grosso | Brazil |
| AF1 | AF | 9.87859 S 56.09733 W | 2021 | Alta Floresta, Mato Grosso | Brazil |
| AF2 | AF | 9.87858 S 56.09725 W | 2021 | Alta Floresta, Mato Grosso | Brazil |
| AF3 | AF | 9.87504 S 56.06598 W | 2021 | Alta Floresta, Mato Grosso | Brazil |
| AF4 | AF | 9.797576 S 55.862469 W | 2021 | Alta Floresta, Mato Grosso | Brazil |
| AF5 | AF | 9.87171 S 56.10136 W | 2021 | Alta Floresta, Mato Grosso | Brazil |
| AF6 | AF | 9.91492 S 55.913048 W | 2021 | Alta Floresta, Mato Grosso | Brazil |
| AF7 | AF | 9.86991 S 56.0953 W | 2021 | Alta Floresta, Mato Grosso | Brazil |
| AF8 | AF | 9.820971 S 55.949661 W | 2021 | Alta Floresta, Mato Grosso | Brazil |
| AF9 | AF | 9.915494 S 55.91319 W | 2021 | Alta Floresta, Mato Grosso | Brazil |
| AF10 | AF | 9.914716 S 55.913742 W | 2021 | Alta Floresta, Mato Grosso | Brazil |
| AF11 | AF | 9.90977 S 55.913642 W | 2021 | Alta Floresta, Mato Grosso | Brazil |
| AF12 | AF | 9.910132 S 55.913693 W | 2021 | Alta Floresta, Mato Grosso | Brazil |
| AF13 | AF | 9.909962 S 55.913717 W | 2021 | Alta Floresta, Mato Grosso | Brazil |
| AF14 | AF | 9.909474 S 55.913602 W | 2021 | Alta Floresta, Mato Grosso | Brazil |
| AF15 | AF | 9.909598 S 55.913858 W | 2021 | Alta Floresta, Mato Grosso | Brazil |
| AF16 | AF | 9.913309 S 55.914653 W | 2021 | Alta Floresta, Mato Grosso | Brazil |
| AF17 | AF | 9.912382 S 55.914518 W | 2021 | Alta Floresta, Mato Grosso | Brazil |
| AF18 | AF | 9.914904 S 55.913056 W | 2021 | Alta Floresta, Mato Grosso | Brazil |
| AF19 | AF | 9.915494 S 55.91319 W | 2021 | Alta Floresta, Mato Grosso | Brazil |
| AF20 | AF | 9.915732 S 55.91194 W | 2021 | Alta Floresta, Mato Grosso | Brazil |
| AF21 | AF | 9.91515 S 55.91123 W | 2021 | Alta Floresta, Mato Grosso | Brazil |
| AF22 | AF | 9.915101 S 55.911134 W | 2021 | Alta Floresta, Mato Grosso | Brazil |
| AF23 | AF | 9.914551 S 55.911536 W | 2021 | Alta Floresta, Mato Grosso | Brazil |
| AF24 | AF | 9.914793 S 55.912813 W | 2021 | Alta Floresta, Mato Grosso | Brazil |
| AF25 | AF | 9.914093 S 55.913856 W | 2021 | Alta Floresta, Mato Grosso | Brazil |
| AF26 | AF | 9.914041 S 55.913824 W | 2021 | Alta Floresta, Mato Grosso | Brazil |
| AF27 | AF | 9.80125 S 55.892417 W | 2021 | Alta Floresta, Mato Grosso | Brazil |

| | | | | | | |
|---|---|---|---|---|---|---|
| AF28 | AF | 9.82109 S 55.9492 W | 2021 | Alta Floresta, Mato Grosso | Brazil | |
| AF29 | AF | 9.82129 S 55.9494 W | 2021 | Alta Floresta, Mato Grosso | Brazil | |
| AF30 | AF | 9.82098 S 55.9503 W | 2021 | Alta Floresta, Mato Grosso | Brazil | |
| AF31 | AF | 9.81671 S 55.958 W | 2021 | Alta Floresta, Mato Grosso | Brazil | |
| AF32 | AF | 9.915676 S 55.91286 W | 2022 | Alta Floresta, Mato Grosso | Brazil | |
| AF33 | AF | 9.915986 S 55.912565 W | 2022 | Alta Floresta, Mato Grosso | Brazil | |
| AF34 | AF | 9.916215 S 55.910737 W | 2022 | Alta Floresta, Mato Grosso | Brazil | |
| AF35 | AF | 9.916239 S 55.910929 W | 2022 | Alta Floresta, Mato Grosso | Brazil | |
| AF36 | AF | 9.915775 S 55.909336 W | 2022 | Alta Floresta, Mato Grosso | Brazil | |
| AF37 | AF | 9.915566 S 55.90909 W | 2022 | Alta Floresta, Mato Grosso | Brazil | |
| AF38 | AF | 9.911625 S 55.911157 W | 2022 | Alta Floresta, Mato Grosso | Brazil | |
| TCN1 | TCN | 7.907837 S 34.870609 W | 2025 | Abreu e Lima, Pernambuco | Brazil | Tucunacá |
| TCN2 | TCN | 7.907837 S 34.870609 W | 2025 | Abreu e Lima, Pernambuco | Brazil | Tucunacá |
| TCN3 | TCN | 7.907837 S 34.870609 W | 2025 | Abreu e Lima, Pernambuco | Brazil | Tucunacá |
| TCN4 | TCN | 7.907837 S 34.870609 W | 2025 | Abreu e Lima, Pernambuco | Brazil | Tucunacá |
| TCN5 | TCN | 7.907837 S 34.870609 W | 2025 | Abreu e Lima, Pernambuco | Brazil | Tucunacá |
| TCN6 | TCN | 7.907837 S 34.870609 W | 2025 | Abreu e Lima, Pernambuco | Brazil | Tucunacá |
| CPR1 | CPR | 7.907837 S 34.870609 W | 2025 | Abreu e Lima, Pernambuco | Brazil | Caupuri |
| CPR2 | CPR | 7.907837 S 34.870609 W | 2025 | Abreu e Lima, Pernambuco | Brazil | Caupuri |
| CPR3 | CPR | 7.907837 S 34.870609 W | 2025 | Abreu e Lima, Pernambuco | Brazil | Caupuri |
| CPR4 | CPR | 7.907837 S 34.870609 W | 2025 | Abreu e Lima, Pernambuco | Brazil | Caupuri |
| CPR5 | CPR | 7.907837 S 34.870609 W | 2025 | Abreu e Lima, Pernambuco | Brazil | Caupuri |
| CPR6 | CPR | 7.907837 S 34.870609 W | 2025 | Abreu e Lima, Pernambuco | Brazil | Caupuri |
| CPR7 | CPR | 7.907837 S 34.870609 W | 2025 | Abreu e Lima, Pernambuco | Brazil | Caupuri |
| CPR8 | CPR | 7.907837 S 34.870609 W | 2025 | Abreu e Lima, Pernambuco | Brazil | Caupuri |
| CPR9 | CPR | 7.907837 S 34.870609 W | 2025 | Abreu e Lima, Pernambuco | Brazil | Caupuri |
| TCN_ref | TCN | 7.907837 S 34.870609 W | 2022 | Abreu e Lima, Pernambuco | Brazil | Tucunacá |
| CPR_ref | CPR | 7.907837 S 34.870609 W | 2022 | Abreu e Lima, Pernambuco | Brazil | Caupuri |

Highlighted rows: reference-genome accessions (HiFi/Hi-C); collection year and coordinates to be confirmed. Coordinates of the TCN/CPR plantatic

ɔn are close to the Abreu e Lima/Paulista municipal border.
